# Deep reinforcement learning-driven discovery of a MsbA-targeted small-molecule antibiotic for the treatment of *Acinetobacter baumannii* infection

**DOI:** 10.64898/2026.09.12.751098

**Authors:** Lei Wang, Baixin Jing, Qingxin Liu, Jiale Zhong, Zunsheng Han, Yingying Yang, Hongwei Jin, Yanyan Li, Liang Peng, Song Wu, Jie Xia

## Abstract

Antibiotics with new mechanisms are highly pursued to address the threat of infections caused by drug-resistant Gram-negative bacteria. Targeting MsbA, a key protein of the lipopolysaccharide biosynthesis pathway, represents a promising strategy to discover new classes of antibiotics. However, currently available MsbA-targeted molecules either lack sufficient potency or have unfavorable properties, necessitating expansion of chemical space. In this study, we chose the most promising cerastecin **Cpd 4** as the template, and used two Artificial Intelligence (AI)-based tools, i.e. Link-INVENT and AutoMolDesigner for molecular design, performed chemical derivatization and antibacterial activity evaluation, which led to the discovery of **Y-11** (MIC for *A. baumannii*: 0.5 μg/mL). Encouragingly, **Y-11** showed equivalent potency to **Cpd4** for carbapenem-resistant *A. baumannii*, and less cytotoxicity and hemolysis as well as lower spontaneous resistance frequency. *In vivo* efficacy study demonstrated that **Y-11** could effectively reduce bacterial loads in the mice infected by *A. baumannii*. The following mechanism study including molecular dynamics simulation, biochemical assay, and transmission electron microscope (TEM) analysis suggested that **Y-11** inhibited the transport of lipooligosaccharide and impaired the formation of outer membrane, probably by competitively binding to the substrate binding site of MsbA and modulating ATPase activity. Taken together, we have discovered a MsbA-targeted small molecule **Y-11** via AI-driven drug design, which provides a foundation for future antibiotic development.

**Graphical abstract:** 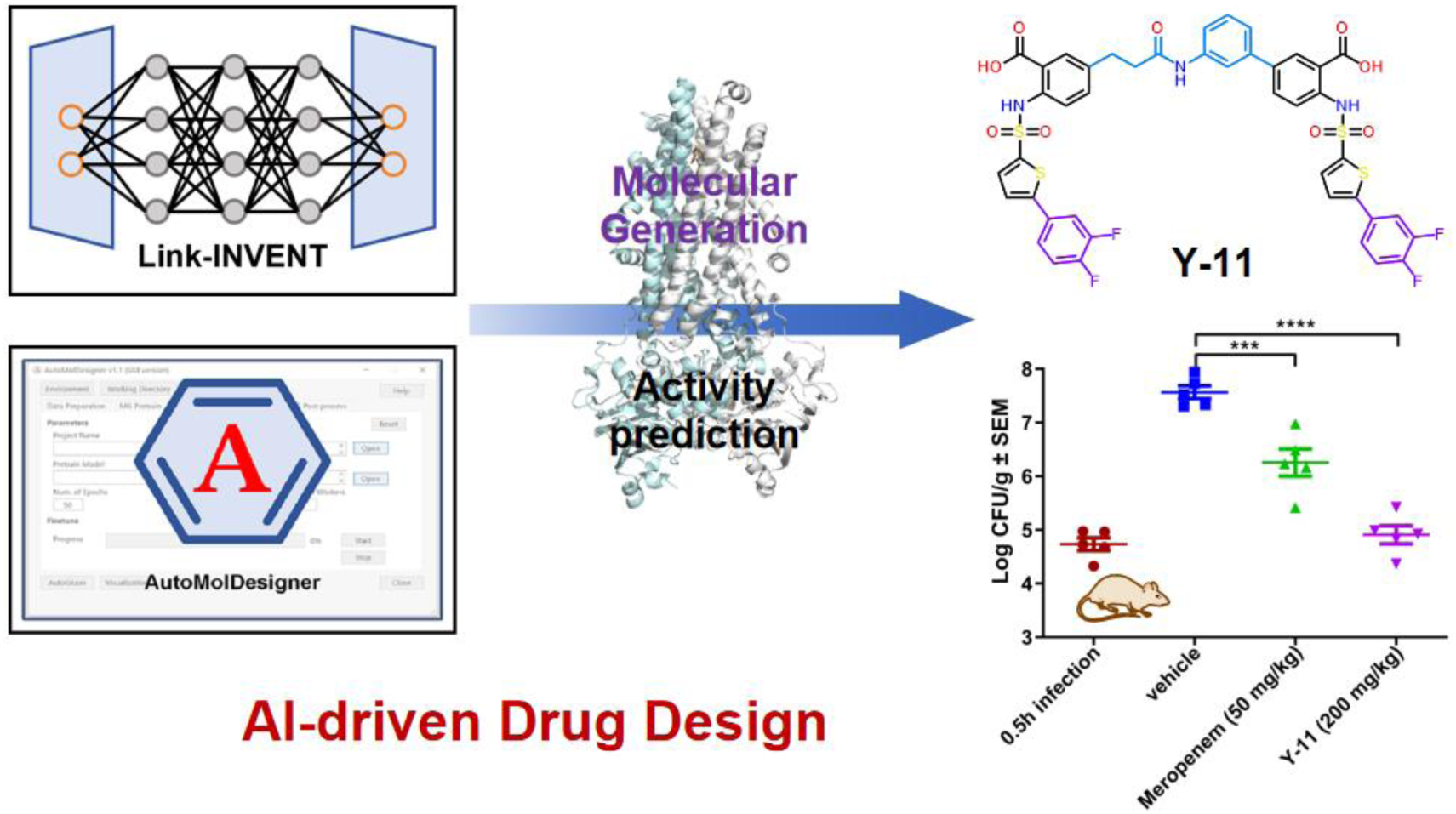

## 1. Introduction

Antibiotic resistance has been a serious threat to public health worldwide. ^1^ *Enterococcus faecium*, *Staphylococcus aureus*, *Klebsiella pneumoniae*, *Acinetobacter baumannii* (*A. baumannii*), *Pseudomonas aeruginosa*, and *Enterobacter species*, i.e. ‘ESKAPE’, are the pathogens that are easy to develop resistance to antibiotics, and thus are of high concern. ^2, 3^ Among them, *A. baumannii* is on the top of WHO Bacterial Priority Pathogens List for novel therapies and enhanced prevention measures, as carbapenem-resistant *A. baumannii* (CRAB) may cause serious infections, e.g. ventilator-associated pneumonia and bloodstream infections, that are difficult to treat and threaten human life. ^4, 5^ Since small-molecule antibiotics are commonly used and highly effective therapeutics, and new chemotypes are promising to address antibiotic resistance, new targets for drug discovery attract much interest from academia and industry. ^6^

MsbA, a member of the ATP binding cassette (ABC) transporter superfamily, is one of the interesting targets. It is located at the inner membrane of gram-negative bacteria, and mediates the transport of lipooligosaccharide (LOS), a precursor of lipopolysaccharide (LPS), from the inner leaflet to the outer leaflet of the inner membrane. ^7–10^ In terms of molecular mechanism, the transport depends on the binding of ATP to MsbA and hydrolysis to provide energy for the transition from open inward-facing conformation to open outward-facing one. ^11, 12^ MsbA-targeted small molecules that affect the conformational transition are able to block LOS transport, which eventually impairs integrity of bacterial outer membrane and thus cause bacterial death.

Although there are three classes of MsbA inhibitors, including tetrahydrobenzothiophenes (TBTs), quinoline acrylic acids, and cerastecins, none of them has entered clinical trials. TBTs are weak small-molecule antibiotics, for instance, **TBT1** (**1**) only has its MIC value of 20 μg/mL for *A. baumannii*. ^13^ Quinoline acrylic acids could potently inhibit MsbA, but are rather weak in terms of antibacterial activity (e.g. **G907** (**2**), cf. **Figure 1**). ^14^ Cerastecins are the most promising class. Since the identification of **cerastecin A** (**3**) from high-throughput screening, hit-to-lead optimization efforts have been taken, which led to two MsbA inhibitors with strong antibacterial activity, **cerastecin C** (**4**) and **Cpd 4** (**5**) (cf. **Figure 1**). ^15, 16^ Nevertheless, the reported cerastecins lack favorable ADME/T properties, in particular plasma protein binding (PPB). ^15^ Accordingly, chemical space of MsbA inhibitors are necessary to expand.

**Figure 1.**
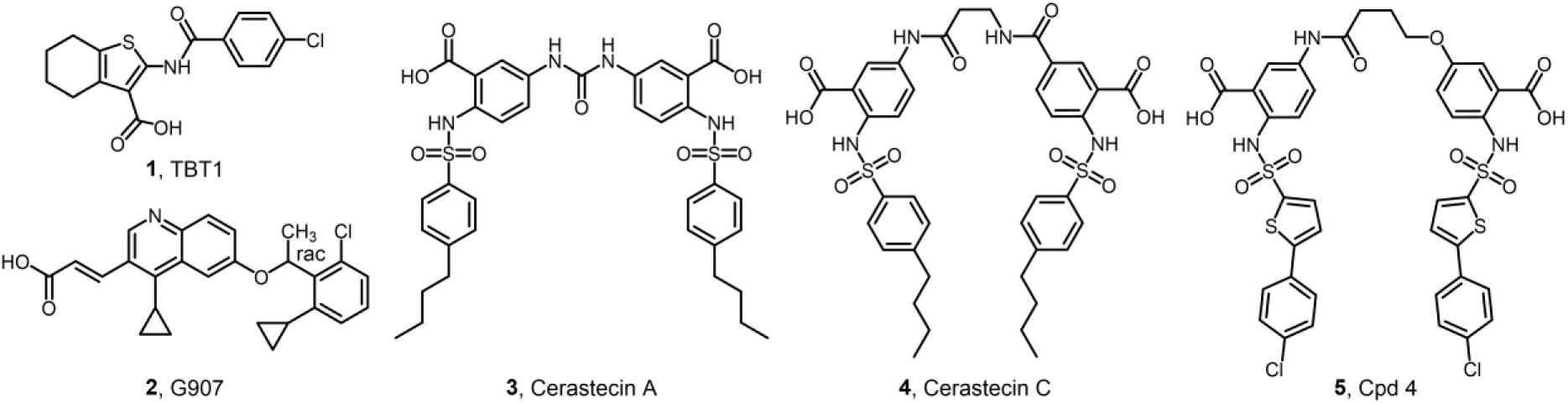
Chemical structures and antibacterial activity of currently available MsbA inhibitors:

### TBT1, G907, cerastecin A, cerastecin C, Cpd 4

Artificial intelligence (AI) is a powerful technique to accelerate drug discovery. In the field of small-molecule antibiotic discovery, it could be used for virtual screening or *de novo* drug design and has let to cases of success. For instance, Stokes et al. developed AI-based property prediction tool, Chemprop, and discovered Halicin and Abaucin as new chemotypes of small-molecule antibiotics by virtual screening. ^17, 18^ They further developed CReM and VAE, AI-based tools for *de novo* antibiotic design, and discovered NG1 for *N. gonorrhoeae* and DN1 for MRSA. ^19^ To the best of our knowledge, AI has not been applied to MsbA-targeted drug discovery.

This study is aimed to use AI to discover novel MsbA inhibitors that are structurally different from the previously reported ones for the treatment of *A. baumannii* infection. Firstly, we constructed an AI-integrated computational workflow for *de novo* design of MsbA-targeted small-molecule antibiotics. The workflow mainly included: (1) the generation of a focused library of potential MsbA inhibitors by Link-INVENT, a reinforcement learning-based tool for linker design, (2) molecular docking against the substrate binding site of MsbA, (3) anti-*A. baumannii* activity prediction by machine learning model built with AutoMolDesigner. ^20, 21^ The hit identified by the workflow was further derivatized, which provided **Y-11** with improved antibacterial activity and lower toxicity. Additionally, we demonstrated **Y-11** was effective to treat the mice infected by *A. baumanni*, with the mechanism of modulating MsbA ATPase activity, and impairing bacterial LOS transport and outer membrane integrity. Our results suggest N-phenylpropionamide–linked cerastecins constitute a promising new class of small-molecule antibiotics.

## 2. Results and Discussion

### 2.1. AI-driven hit identification

#### 2.1.1. Virtual chemical library of cerastecins generated by Link-INVENT

By analyzing the structures and anti-*A. baumannii* activity of representative cerastecins from the patents of Merck (cf. **Figure S1**), we noted that the type of linkers is the key to anti-*A. baumannii* activity of cerastecins. ^22, 23^ To be specific, the length and conformation of the linker determine the distance between the two carboxyl groups of cerastecins, thus may affect the formation of salt bridge between the carboxyl group of cerastecins and Arg-72 of MsbA, as observed in the Cryo-EM structure of MsbA (cf. **Figure S2**). Accordingly, we focused on the design of new linkers in this work.

Link-INVENT, an extension of REINVENT v3.2, is a computational tool based on recurrent neural network (RNN) and Reinforcement learning (RL) specifically for linker design. ^20^ In Link-INVENT, a prior network (RNN) is available to generate syntactically valid linkers, while the Agent gradually learns to generate favorable linkers connecting molecular subunits that satisfy multi-parameter optimization (MPO) objectives. Thus, this tool is applicable for our purpose. As **Cpd 4** (cf. **Figure 1**) is the most active MsbA-targeted antibiotic reported so far, we selected it as a template molecule. Firstly, we chose its identical fragments of 2-((5-(4-chlorophenyl)thiophene)-2-sulfonamido)benzoic acid at both sides as the warheads and the 5-position of the benzoic acid ring as the attachment points (cf. **Figure 2A**). Then, we defined four property constraints for the linker design: effective length from 4 to 7 bonds, 0 or 1 aliphatic ring, 0 or 1 aromatic ring, molecular weight between 70 and 200. If a generated linker met all four constraints, a Finalscore of 1 was assigned to the linked molecule. The prior network (RNN) of Link-INVENT was trained for a total of 200 RL steps. As shown in **Figure S3**, the learning curve of the average Finalscore converged after 14 RL steps. Three components, i.e. the counts of aliphatic rings, the counts of aromatic rings, the molecular weight of the generated linkers, converged rapidly, while the length of the linkers exhibited slow convergence. After 14 steps, the proportion of molecules meeting the design criteria (FinalScore =1) exceeded 80% in each cycle of molecular generation, and the average Finalscore was maintained between 0.74 and 1. During the whole RL process, 20,466 linked molecules were generated. **Figure 2B** shows frequency distribution of four properties of the linkers generated by Link-INVENT, including linker_effective_length, aliphatic_rings, aromatic_rings and linker_molecular_weight. Properties of most linkers are within the ranges predefined in the RL process. Among all the generated molecules, 18,372 compounds (89.77%) had a FinalScore of 1, i.e. meeting all the predefined criteria, which constituted the virtual chemical library of cerastecins.

**Figure 2.**
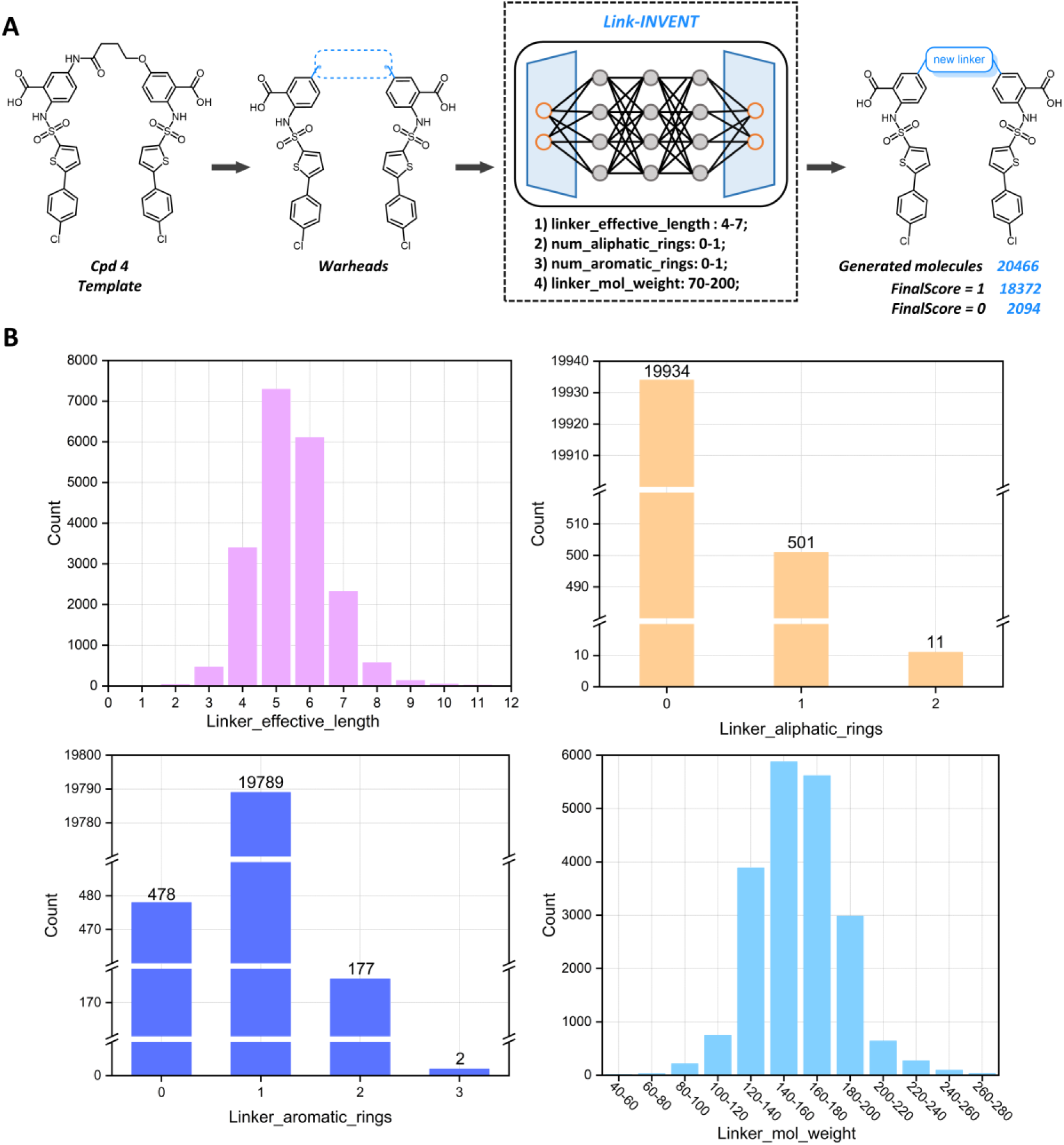
Generation of a virtual chemical library of cerastecins with diverse linkers using Link-INVENT. (A) The workflow of linker generation. With the SMILES notations of the bilateral warheads of **Cpd 4** as the input, the Agent generated 18,372 molecules that met the predefined criteria (FinalScore = 1). (B) Property distribution of the generated linkers.

#### 2.1.2 In-1 as a hit identified from the virtual library

##### 2.1.2.1 Computational workflow

As known cerastecins are MsbA-targeted antibiotics, it is natural to take both MsbA binding and antibacterial activity into consideration. With this aim, we firstly performed molecular docking against the substrate binding site of MsbA with the FRED program (cf. **Figure 3A**). ^24, 25^ We only selected 2,636 linked molecules that had better scores than **Cpd 4** (Chemgauss4 score: -12.77) as potential MsbA binders. To select potential antibacterial compounds from them, we used AutoMolDesigner (V1.1) to build machine learning models and predict anti-*A. baumannii* activity. ^21, 26^ With the diverse dataset composed of 393 actives and 504 inactives against *A. baumannii*, we built 5 ensemble models based on RDKit_2D_norm, RDKit_2D, ECFP_4, FCFP_6 and MACCS, respectively. As shown in **Table S1**, the ECFP_4 based model (**M3**) achieved the best predictive performance, with the highest accuracy (0.956), MCC (0.910) and F1 score (0.950). This model suggested 1,387 molecules with potential anti-*A. baumannii* activity (i.e. possibility > 0.5). To narrow down the number of compounds to synthesize, we first used AutoMolDesigner (V1.1) to evaluate synthetic accessibility (SA) and selected top 10% molecules (SA Score: 3.0 ∼ 3.3), then visually inspected their binding modes and found only 19 linked molecules could form salt bridges between their two carboxyl groups and Arg-72 (cf. **Table S2**). ^27^ With expertise on synthetic chemistry, we selected **In-1** as the candidate compound for experimental validation.

**Figure 3.**
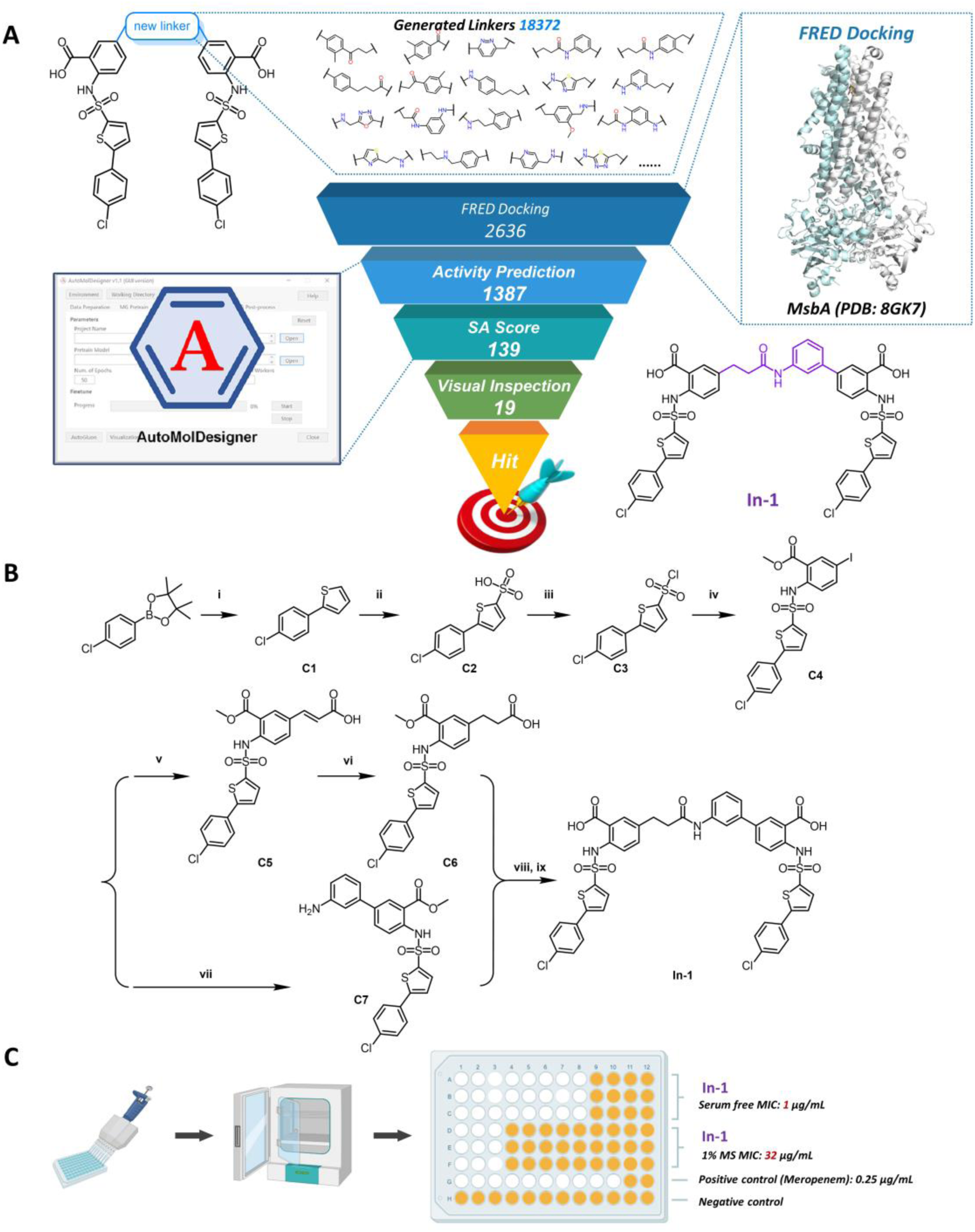
Identification of the hit **In-1** via virtual screening from the library of AI-generated cerastecins, chemical synthesis and anti-*A. baumannii* activity evaluation. (A) The workflow of virtual screening, including molecular docking of the virtual chemical library against MsbA (PDB ID: 8GK7), anti-*A. baumannii* activity prediction and SA Score calculation with AutoMolDesigner, visual inspection of binding mode and synthetic feasibility evaluation by human experts on medicinal chemistry, which suggested **In-1** as the most promising molecule. (B) The synthetic route of **In-1**. i) 2-iodothiophene, Pd(PPh_3_)_4_, K_2_CO_3_, 1,4-dioxane, H_2_O, reflux, 2h; ii) Ac_2_O, H_2_SO_4_, EtOAc, 24h; iii) SOCl_2_, DMF, reflux, 8h; iv) methyl 2-amino-5-iodobenzoate, pyridine, DCM, 12h; v) acrylic acid, Pd(dppf)Cl_2_, TEA, DMF, 100℃, 2h; vi) H_2_, Pd/C, CH_3_OH, DCM, 2h; vii) (3-aminophenyl)boronic acid, Pd(PPh_3_)_4_, K_2_CO_3_, 1,4-dioxane, H_2_O, 60℃, 2h; viii) TCFH, NMI, DCM, 12h; ix) LiOH (aq), THF, CH_3_OH, 12h. (C) Evaluation of **In-1** for its activity against *A. baumannii* (ATCC 19606). Serum free MIC: MIC(triplicate) of **In-1** without mouse serum (MS); 1% MS MIC: MIC (triplicate) of **In-1** with 1% mouse serum; Positive control: Meropenem.

##### 2.1.2.2 Chemical synthesis of In-1

Compound **In-1** was synthesized as outlined in **Figure 3B**. With 4-chlorophenylboronic acid pinacol ester as the starting material, Suzuki coupling with 2-iodothiophene afforded intermediate **C1**. Treated with concentrated sulfuric acid and acetic anhydride, **C1** was converted to **C2**, which was then reacted with thionyl chloride to give **C3**. Condensation with methyl 2-amino-5-iodobenzoate yielded **C4**. Subsequent Heck coupling of **C4** with acrylic acid provided **C5**. Catalytic hydrogenation over Pd/C reduced **C5** to **C6**. Suzuki coupling of **C4** with 3-aminophenylboronic acid provided **C7**. Amide coupling between **C6** and **C7** using N,N,N’,N’-Tetramethylchloroformamidinium hexafluorophosphate (TCFH) followed by hydrolysis afforded the target compound **In-1**. The following facts in the NMR spectra (cf. **Figure S4**) validated the structures of the synthesized compound **In-1**: (1) In the ^1^H NMR spectra, the protons of two alkyl carbons (i.e. CH_2_-CH_2_) of the linker, revealed peaks at *δ* 2.59 and 2.86 ppm, and both of them appeared as triplets; (2) In the ^13^C NMR spectra, the above groups revealed peaks at *δ* 29.9 and 37.8 ppm in DMSO-*d_6_*.

##### 2.1.2.3 Anti-A. baumannii activity of **In-1**

We tested compound **In-1** for its activity against *A. baumannii* (ATCC19606), and its MIC value was 1 μg/mL. It indicated our computational workflow of AI-integrated *de novo* design was effective for hit identification. To explore whether PPB affected the activity of **In-1**, we determined MICs of **In-1** for *A. baumannii* in the presence of 1% mouse serum (MS). As a result, MIC of **In-1** was enhanced from 1 μg/mL to 32 μg/mL, indicating **In-1**, similar to those reported cerastecins, had the issue of a high PPB rate.

### 2.2. Chemical derivatization of In-1

#### 2.2.1 Synthesis of In-1 derivatives

In order to improve antibacterial activity for *A. baumannii*, we performed derivatization on the hydrophobic 4-chlorophenyl moieties at both ends of **In-1**, by simultaneously replacing them with substituted phenyl groups, e.g. 3-fluorophenyl, 4-fluorophenyl, 3,4-difluorophenyl (cf. **Figure 4A**). A total of 18 derivatives (**Y-1** to **Y-17**) were synthesized according to the synthetic route outlined in **Figure 4B**. Starting from ethyl 2-amino-5-iodobenzoate, Heck coupling with *tert*-butyl acrylate afforded **C8**, which was hydrogenated over Pd/C to give **C9**. **C9** was condensed with 5-bromothiophene-2-sulfonyl chloride, followed by deprotection of the *tert*-butyl group under trifluoroacetic acid (TFA) treatment to afford the carboxylic acid intermediate. In parallel, Suzuki coupling of ethyl 2-amino-5-iodobenzoate with ((tert-butoxycarbonyl)amino)phenyl)boronic acid provided **C10**, which was condensed with 5-bromothiophene-2-sulfonyl chloride and deprotected to give amine intermediate. Key amide coupling using TCFH produced **C11**. Lastly, Suzuki coupling of **C11** with various arylboronic acid, followed by hydrolysis, afforded the target compounds **Y-1**∼**Y-17**.^16^ In NMR spectra, compound **Y-1**∼**Y-17** exhibited peaks similar to those of **In-1** and thus were easy to validate.

**Figure 4.**
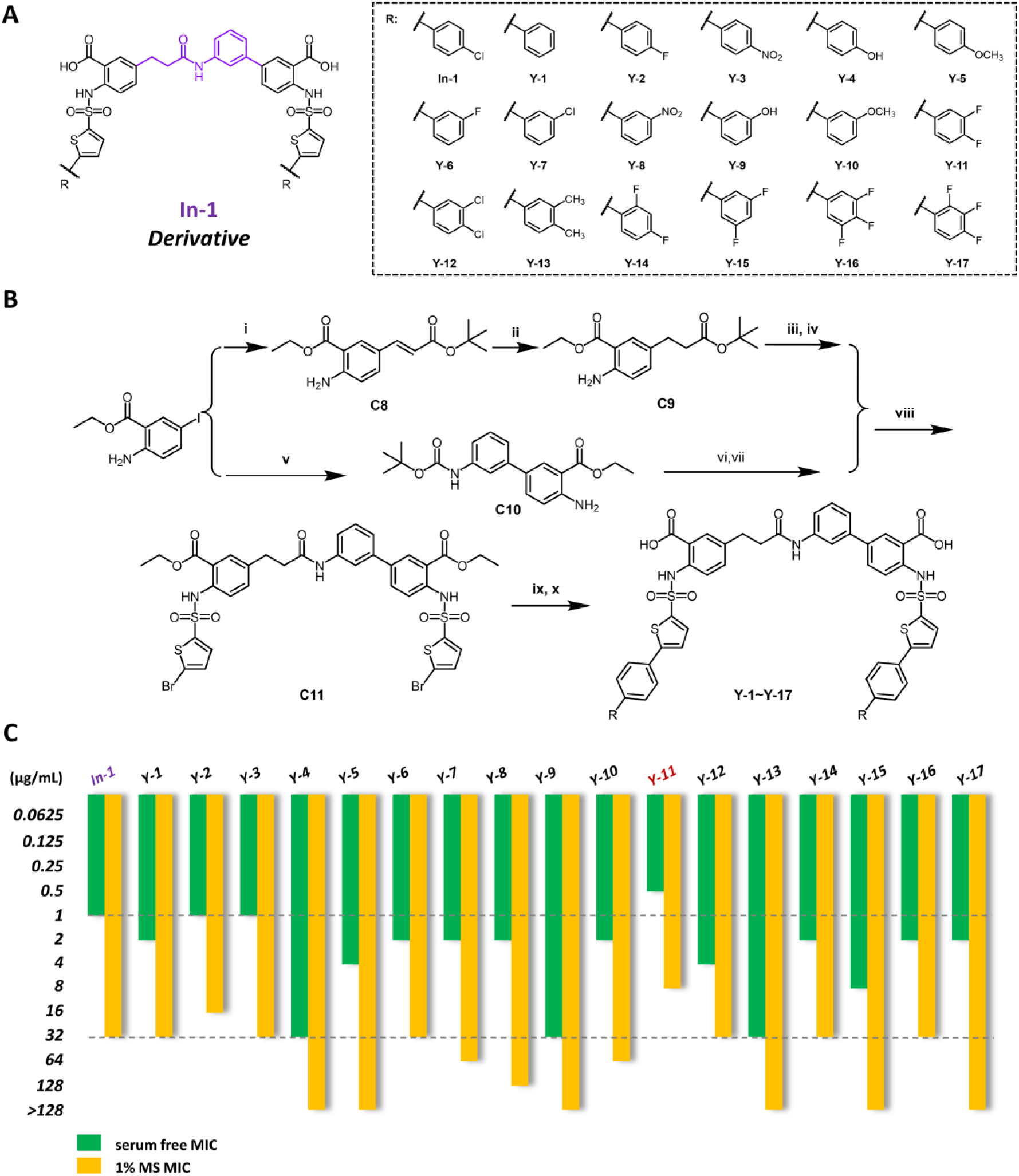
Derivatization of **In-1** and anti-*A. baumannii* activity evaluation. (A) Chemical structures of the 17 derivatives (**Y-1** ∼ **Y-17**). (B) The general synthetic route of **Y-1**∼**Y-17**. i) *tert*-butyl acrylate, Pd(dppf)Cl_2_, TEA, DMF, 100℃, 2h; ii) H_2_, Pd/C, CH_3_OH, DCM, 2h; iii) 5-bromothiophene-2-sulfonyl chloride, pyridine, DCM, 12h; iv) TFA, DCM, 2h; v) (3-((tert-butoxycarbonyl)amino)phenyl)boronic acid, Pd(PPh_3_)_4_, K_2_CO_3_, 1,4-dioxane, H_2_O, reflux, 2h; vi) 5-bromothiophene-2-sulfonyl chloride, pyridine, DCM, 12h; vii) 4M HCl 1,4-dioxane, 2h; viii) TCFH, NMI, DCM, 12h; ix) arylboronic acid, Pd(PPh_3_)_4_, K_2_CO_3_, 1,4-dioxane, H_2_O, reflux, 2h; x) LiOH (aq), THF, CH_3_OH, 12h. (C) Antibacterial activity for *A. baumannii* (ATCC 19606) of **In-1** and its 17 derivatives. Serum free MIC (green): MIC of compounds without mouse serum; 1% MS MIC (yellow): MIC of compounds with 1% mouse serum.

#### 2.2.2 Anti-A. baumannii activity of In-1 derivatives

We tested all the synthesized derivatives for their antibacterial activity (i.e. MIC) for *A. baumannii* (ATCC19606), without or with 1% MS. The chemical structures of 18 compounds (**In-1** and **Y-1**∼**Y-17**) and MIC values are shown in **Figure 4C**. Without 1% MS (i.e. under the serum-free condition), the MIC values ranged from 0.5 μg/mL to 32 μg/mL, indicating that all the derivatives inhibited the growth of *A. baumannii* (ATCC19606). Encouragingly, under the serum-free condition, **Y-11** was more potent than **In-1** (0.5 μg/mL vs. 1 μg/mL). In the presence of 1% MS, MIC of **Y-11** became 8 μg/mL while MIC of **In-1** was 32 μg/mL. The above data demonstrated the chemical derivatization used here was effective for hit-to-lead optimization. In addition, it is very likely that the derivatization improved PPB property because MIC decreased by 16 fold from the serum-free condition to the condition with 1% MS for **Y-11**, compared with 32 fold for **In-1** (cf. **Figure 4C**).

#### 2.2.3 SAR of the In-1 derivatives

Although other derivatives were not as potent as **Y-11** at both conditions (without or with 1% MS), we got insights into the SAR of these derivatives with the N-phenylpropionamide linker.

Compared with the derivative without substitution (**Y-1**, 2 μg/mL), the substitution of *para*-fluro (**Y-2**), *para*-nitro (**Y-3**) as well as *para*-chloro (**In-1**) improved antibacterial activity (MIC: 1 μg/mL), while the *para*-hydroxyl (**Y-4**, 32 μg/mL), *para*-methoxy (**Y-5,** 4 μg/mL) substitution caused reduced activity. Accordingly, we could conclude that the introduction of electron-withdrawing groups at the *para*-position of the phenyl group was favorable for antibacterial activity, while the electron-donating group had unfavorable effect. However, this effect seemed lost when the substitution at the *meta*-position, as the MIC value of **Y-6** (*meta*-fluro), **Y-7** (*meta*-chloro) and **Y-8** (*meta*-nitro), and **Y-10** (meta-methoxy) was the same as that of **Y-1**, i.e. 2 μg/mL. Besides, for most of the derivatives with one *meta*-substituent (**Y-6**, **Y-7**, **Y-8**), they showed weaker activity than the derivatives with one *para*-substituent (**Y-2**, **In-1** and **Y-3**), indicating the *para-*substitution was favorable for antibacterial activity. The derivative substituted by 3,4-difluorine (**Y-11,** 0.5 μg/mL) was the most potent. Those compounds with 3,4-dichloro (**Y-12,** 4 μg/mL), 3,4-dimethyl (**Y-13**, 32 μg/mL) were less potent than **Y-11**, indicating the importance of fluorine. However, when the position of di-difluorine substitution changed, e.g. 2,4-difluorine (**Y-14**, 2 μg/mL), 3,5-difluorine (**Y-15**, 8 μg/mL), antibacterial activity decreased. As the derivatives with three fluorines (MIC for **Y-16**, **Y-17**: 2 μg/mL) showed weaker activity than **Y-11** (0.5 μg/mL) and **Y-2** (1 μg/mL), antibacterial activity of fluorine-substituted compounds was sensitive to the number of fluorine atoms. As for the SAR between these chemical structures and anti-*A. baumannii* activity in 1% MS, in general it was consistent with the above mentioned SAR, though minor difference also exists.

### 2.3. Biological activity of Y-11

#### 2.3.1. Y-11 as the most promising lead compound

##### 2.3.1.1 Antibacterial activity for carbapenem-resistant A. baumannii

Since **Y-11** was highly active against carbapenem-sensitive *A. baumannii* (CSAB, ATCC 19606) (MIC: 0.5 μg /mL), we were curious to see whether it was effective for carbapenem-resistant *A. baumannii* (CRAB). **Table 1** and **Table S3** confirmed that **Y-11** was still active for CRAB. This compound had the MIC_range_ of 1 μg/mL for CRAB (NCTC 13304, CRAB24-1, CRAB24-2, CRAB 24-3). We also compared **Y-11** with the hit compound **In-1** in terms of activity against CRAB. Consistently, its anti-CRAB activity remained 2-fold that of **In-1** (MIC_range_: 2 μg/mL). Compared with **Cpd 4** (MIC_range_:0.5∼1 μg/mL), the potency of **Y-11** was also very close, which was pretty encouraging.

**Table 1.** Anti-CRAB activity, cytotoxicity and hemolysis of Y-11.

| Compound | CRAB<sup>a</sup><br>MIC<sub>range</sub><br>($\mu\text{g/mL}$) | CC<sub>50</sub><sup>b</sup><br>($\mu\text{g/mL}$) | Serum-free<br>CC<sub>50</sub><sup>c</sup> ($\mu\text{g/mL}$) | HC<sub>50</sub><sup>d</sup><br>($\mu\text{g/mL}$) | Selectivity index<sup>e</sup><br>CC<sub>50</sub>/MIC or HC<sub>50</sub>/MIC |
| --- | --- | --- | --- | --- | --- |
| Cpd 4 | 0.5~1 | > 100 | 13.4 | 11.4 | 11.4 |
| **In-1** | 2 | > 100 | 83.7 | 16.6 | 8.3 |
| **Y-11** | 1 | > 100 | 21.2 | 18.6 | 18.6 |
| Meropenem | $\geq 32$ | n.d.<sup>f</sup> | n.d. | n.d. | n.d. |

<sup>a</sup> CRAB, carbapenem-resistant *A. baumannii* (NCTC 13304, CRAB24-1, CRAB24-2, CRAB 24-3).
<sup>b</sup> Values represent the concentration of compounds that causes 50% cytotoxicity (HEK293) under 10% FBS after 72 h treatment.
<sup>c</sup> Values represent the concentration of compounds that causes 50% cytotoxicity (HEK293) under serum-free condition after 12 h treatment.
<sup>d</sup> Values represent concentration of compounds that causes 50% hemolysis after 3 h treatment.
<sup>e</sup> Selectivity index, the smaller value of CC<sub>50</sub>/MIC and HC<sub>50</sub>/MIC.
<sup>f</sup> n.d., not determined.

##### 2.3.1.2 Cytotoxicity and Hemolysis

We further evaluated *in vitro* toxicity of **Y-11**, including cytotoxicity and hemolysis. Human embryonic kidney 293 (HEK293) cells were used as a representative cell line for cytotoxicity assay. At first, we tested cytotoxicity according to the routine, i.e. cells cultured with 10% fetal bovine serum (FBS) were treated with compounds for 72 h, but did not observe cytotoxicity (CC_50_: >100 μg/mL, cf. **Table1**). It was expected as cerastecins usually had high PPB rates. In order to reduce the effect of PPB, we followed Skudlarek et al. to carry out the assay under the serum-free condition. ^16^ As shown in **Table 1**, the replacement of the linker of **Cpd 4** led to significant reduction in cytotoxicity (Serum-free CC_50_: **Cpd 4**, 13.4 μg/mL vs. **In-1**, 83.7 μg/mL), indicating the effectiveness of our linker design strategy to maintain antibacterial activity while reduce cytotoxicity. Unfortunately, the introduction of two halogens (**Y-11**) to the terminal phenyl group restored cytotoxicity (Serum-free CC_50_: 21.2 μg/mL). Nevertheless, **Y-11** still showed improvement compared with **Cpd 4**.

**Table 1** lists the hemolysis of **Y-11**, **In-1** and the previously reported **Cpd 4**. The 50% hemolysis concentration (HC_50_) value of **Cpd 4** was 11.4 μg/mL. Change of the linker from 4-hydroxybutanamide to N-phenylpropionamide (**In-1**) attenuated hemolytic activity (HC_50_: 16.6 μg/mL). Structural optimization further reduced hemolytic activity (**Y-11,** 18.6 μg/mL).

To comprehensively evaluate these compounds, we calculated selectivity index (CC_50_/MIC and HC_50_/MIC) for further comparison, with greater value indicating higher safety. Accordingly, the selectivity index of **Y-11** was the highest, with the value of 18.6 (cf. **Table1**). These findings highlighted the importance of linker design and chemical derivatization in improving drug-like properties.

#### 2.3.2. Anti-A. baumannii effect of Y-11 in vitro and in vivo

##### 2.3.2.1 Mode of action illustrated by Time-kill curves

In order to determine its mode of action, we studied time-kill kinetics of **Y-11** against *A. baumannii* (ATCC 19606) at three concentrations, i.e., 1 × MIC, 4 × MIC and 8 × MIC, with **Cpd 4** as a comparison and meropenem as a positive control. As shown in **Figure 5A**, at 1 × MIC, 4 × MIC and 8 × MIC, **Y-11** reduced the starting log_10_ CFU/mL (bacteria concentration) by more than 3 log units within 24 h. Therefore, **Y-11** was a bactericidal small-molecule antibiotic. As **Cpd4** showed the same result as **Y-11**, change of the linker and derivatization did not change mode of action of cerastecins.

**Figure 5.**
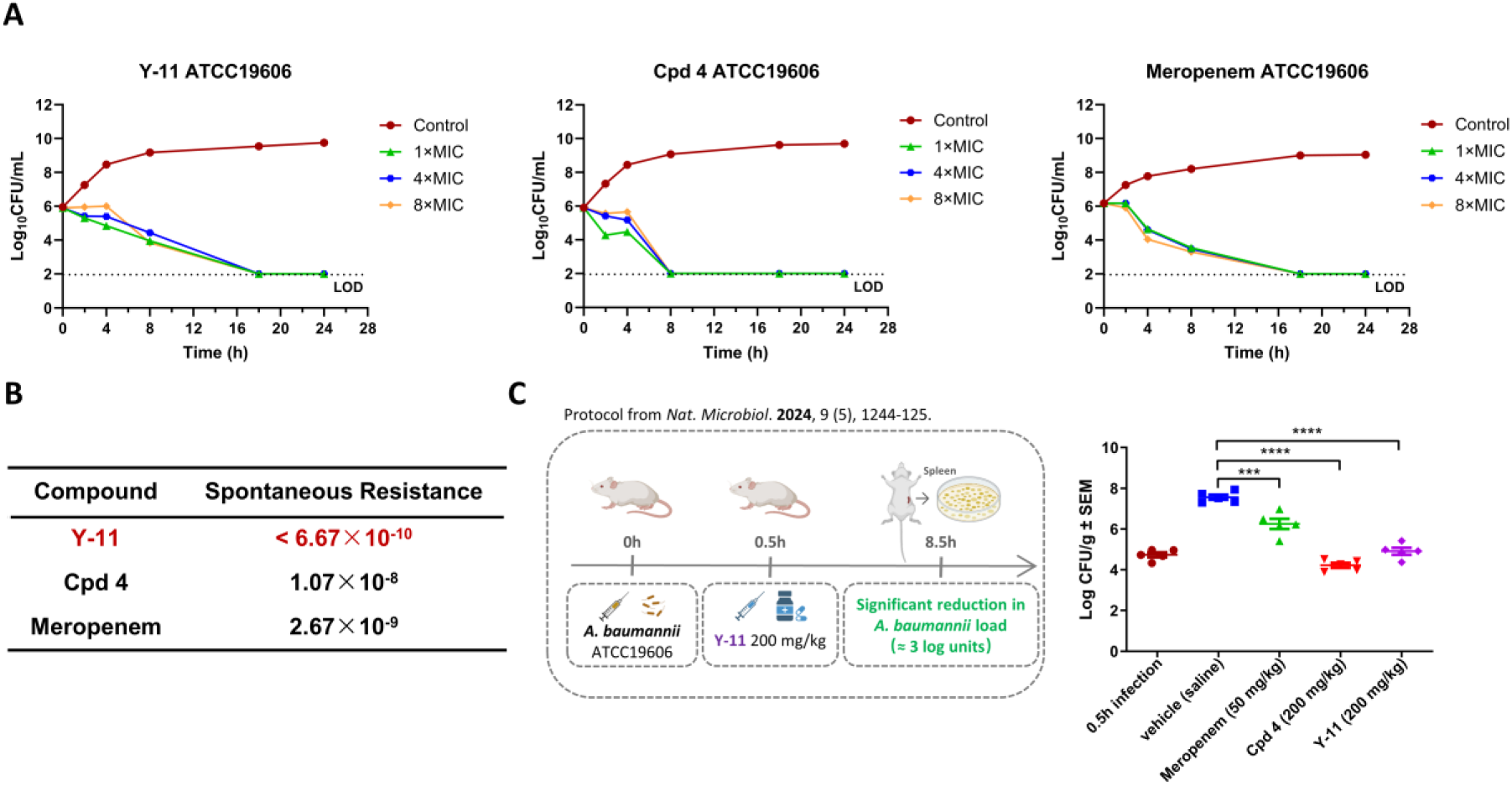
Evaluation of the anti-*A. baumannii* effect of **Y-11**. (A) Time-killing curves of compound **Y-11**, with Meropenem as positive control. (B) Assessment of the spontaneous resistance frequency of *A. baumannii* (ATCC19606) when treated with **Y-11** or Meropenem (positive control). (C) Efficacy was evaluated by using a mouse model infected with *A. baumannii* (ATCC19606) and intraperitoneally injecting **Y-11** at a single dose of 200 mg/kg. Meropenem (positive control) was dosed at 50 mg/kg intraperitoneally as well). Spleen tissue was collected at 8.5 h after infection. Data are presented as mean ± SEM (n = 5). *** p < 0.001, **** p < 0.0001 vs. the vehicle group.

##### 2.3.2.2 Low spontaneous resistance

To evaluate the likeliness to induce resistance, *A. baumannii* (ATCC 19606) was treated with **Y-11** at the concentration of 16 × MIC, with **Cpd 4** as a comparison and meropenem as a control (cf. **Figure 5B**). ^28^ As a result, **Y-11** displayed low spontaneous resistance frequency (< 6.67×10^-10^), a value less than that of both **Cpd 4** (1.07×10^-8^) and meropenem (2.67×10^-9^).

##### 2.3.2.3 Reduced bacterial loads in the infected mice

The *in vivo* pharmacokinetic study in mice indicated the favorable metabolic stability of **Y-11** (cf. **Table S4**), which prompted us to further evaluate its efficacy to treat the mice infected by *A. baumannii* (ATCC 19606). We followed the protocol previously published by Wang, H. *et al* for cerastecins.^15^ At 0.5 h after bacterial infection, the mice were injected intraperitoneally with **Y-11** at the dose of 200 mg/kg. After another 8 h, spleens of the mice were harvested and bacterial loads was quantified. As a result, **Y-11** at a single dose of 200 mg/kg significantly reduced the bacterial loads from 7.6 log_10_(CFU/g) to 4.9 log_10_(CFU/g). Encouragingly, the bacterial load was restored to that status right after infection (4.7 log_10_(CFU/g)), and also similar to that of **Cpd 4** (4.2 log_10_(CFU/g)), indicating we have successfully identified a compound to treat *A.baumanni* infection *in vivo* by AI-based linker design strategy, and this compound was as effective as **Cpd 4**, though it had a linker different from that of **Cpd 4.**

### 2.4. Molecular mechanism

#### 2.4.1. Plausible binding mode to MsbA

To analyze the interactions between **Y-11** and MsbA, we conducted molecular docking, followed by optimization with a 100-ns MD simulation. For the MsbA/**Y-11** complex, it reached the stable state after approximately 30 ns (cf. **Figure S5**). The protein-ligand binding complex after equilibrium was extracted and analyzed. As shown in **Figure 6A**, the key interactions between **Y-11** and MsbA included: (1) the two carboxyl groups formed hydrogen bonds with Arg-72 and Lys-290, respectively; (2) The oxygen atom of the linker formed hydrogen bond with Lys-290; (3) The two hydrophobic fragments at the tails were located at the hydrophobic pocket of MsbA. In general, **Y-11** displayed a serpentine binding pose.

**Figure 6.**
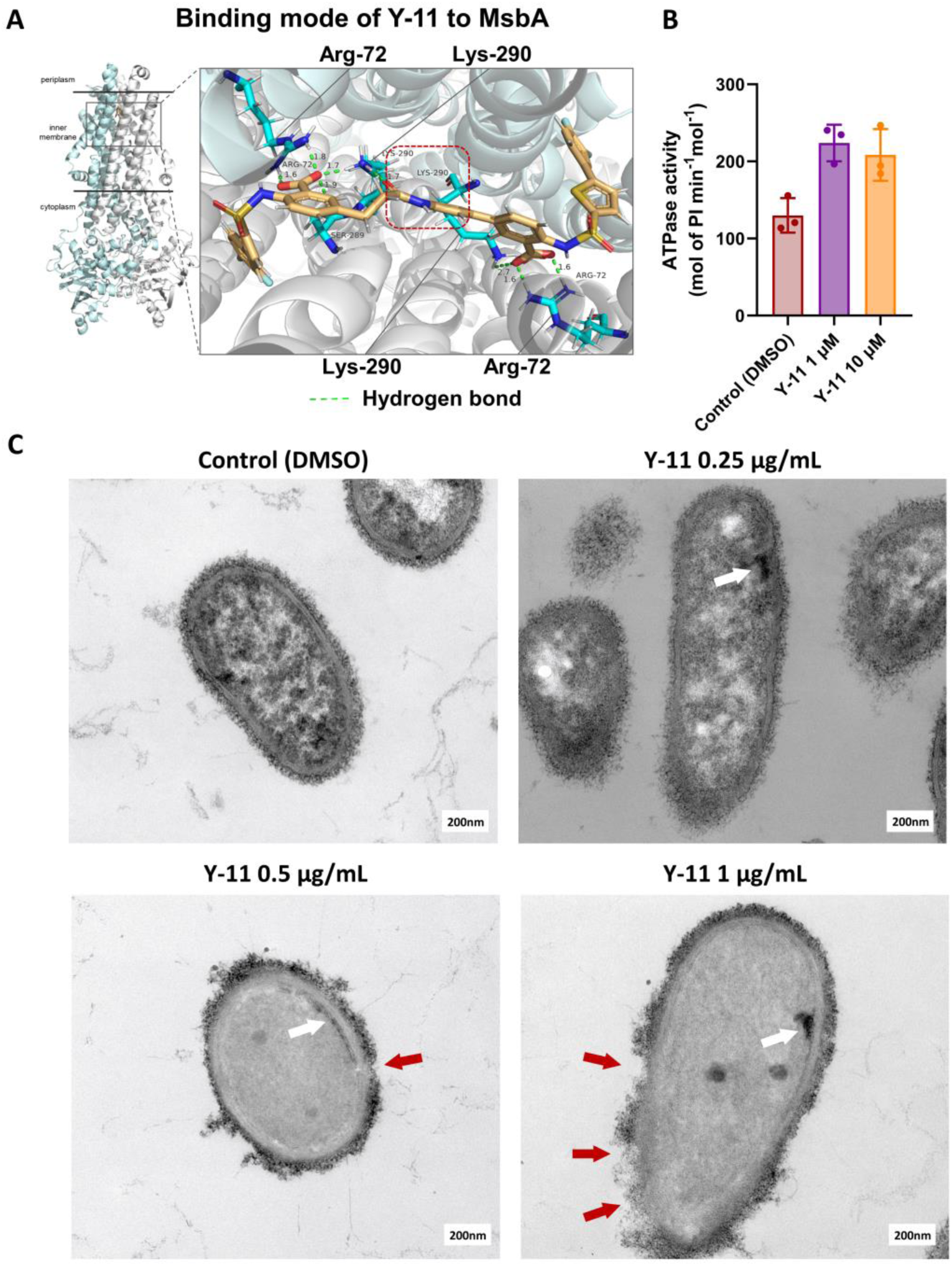
Molecular mechanism of **Y-11** to inhibit *A. baumannii* (ATCC19606). (A) Plausible binding mode of **Y-11** to *Ab*MsbA, predicted by molecular docking and molecular dynamics simulation. (B) ATPase activity assay of *Ab*MsbA in the presence of **Y-11**. (C) The morphology of the outer membrane of *A. baumannii* (ATCC19606), treated with **Y-11** at the concentration of 0.25 µg/mL, 0.5 µg/mL, and 1 µg/mL, detected by transmission electron microscope (TEM). Scale bars, 200 nm. White arrow: accumulation of lipooligosaccharide (LOS); Red arrow: Loss of bacterial outer membrane.

#### 2.4.2. Modulation of MsbA ATPase activity

Recombinant *Ab*MsbA was efficiently overexpressed and purified from *E. coli*, followed by reconstitution into nanodiscs using MSP1D1 and POPG lipids. Detergent was subsequently removed, and the assembled complexes were purified by size-exclusion chromatography for downstream functional characterization. To assess the effects of small molecules on *Ab*MsbA ATPase activity, we used molybdenum blue-based colorimetric assay to measure ATP hydrolysis. ^12^ As shown in **Figure 6B**, **Y-11** exhibited agonistic activity of *Ab*MsbA ATPase at the concentration of 1 μM,. The fold of activation was 1.72. At the concentration of 10 μM, **Y-11** retained the activation activity of *Ab*MsbA ATPase (1.6 fold of control).

#### 2.4.3. Impairment of the outer membrane

We used transmission electron microscope (TEM) to observe the outer membrane of *A. baumannii* that was treated with **Y-11** at the concentration of 0.25 µg/mL (0.5× MIC), 0.5 µg/mL (1 × MIC), and 1 µg/mL (2 × MIC), respectively. As illustrated in **Figure 6C**, the untreated *A. baumannii* (negative control) exhibited a dense and intact outer membrane. Treated with **Y-11** at 0.25 µg/mL, intracellular accumulation of LOS was observed in *A. baumannii* (cf. **Figure 6C**), accompanied by sparse and uneven distribution of LPS in the outer membrane. When the concentration increased to 0.5 µg/mL, intracellular LOS accumulation in the bacteria was consistently detected, and the irregular distribution of LPS in the outer membrane became obvious, with several regions lacking LPS. ^29^ At the concentration of 1 µg/mL, **Y-11** caused marked intracellular LOS accumulation, and significantly impaired the formation of bacterial outer membrane.

These findings preliminarily suggested that **Y-11** could exert its activity against *A. baumannii*, probably because it modulated the activity of MsbA ATPase, and inhibited the transport of LOS and the formation of outer membrane by competitively binding to the substrate binding site of MsbA.

## 3. Conclusion

Antibiotic resistance is a serious threat and urgent issue worldwide. Small-molecule antibiotics with new molecular mechanisms are promising to treat infectious diseases caused by resistant bacteria, e.g. CRAB, while MsbA-targeted compounds belong to the promising class. However, the development of currently available MsbA-targeted compounds is faced with some obstacles including the lack of favorable ADME/T properties.

In this study, we developed a deep reinforcement learning-driven drug design approach by comprehensively using Link-INVENT, AutoMolDesigner as well as traditional CADD technique, and identified **In-1** as a hit compound with anti-*A. baumannii* activity. By chemical derivatization, we obtained **Y-11** with more potent antibacterial activity than **In-1** for both CSAB and CRAB. In terms of antibacterial potency, this compound was similar to the most active MsbA inhibitor, **Cpd 4**. However, this compound had a safety profile and spontaneous resistance frequency better than **Cpd 4**. By *in vivo* efficacy study, **Y-11** could treat bacterial infections caused by *A. baumannii*, showing similar efficacy to **Cpd 4**. Mechanism study has confirmed that **Y-11** is a MsbA-targeted compound. We observed it impaired the bacterial outer membrane and predicted that it competitively bound to the substrate binding site of MsbA. Nevertheless, more mechanism study is needed to understand the interactions between **Y-11** and MsbA.

In summary, this work describes the comprehensive use of AI-driven drug design strategy, chemical synthesis, *in vitro* and *in vivo* bioassays to discover a promising small-molecule antibiotic, **Y-11**. Next work will be the hit-to-lead optimization to improve antibacterial potency and PPB rate.

## 4. Materials and Methods

### 4.1. Computational Modeling

#### 4.1.1. AI-based drug design

##### 4.1.1.1 Generation of a virtual chemical library by Link-INVENT

Link-INVENT (https://github.com/MolecularAI/Reinvent) was used for the linker generation and molecular design. ^20^ Link-INVENT is RL-based tool, thus the MPO objective should be defined for the Agent to train models. Herein, the MPO objective was ComboScore, i.e. the sum of four component scores: effective length, molecular weight, number of aromatic rings, number of aliphatic rings. Firstly, each component (e.g. effective length) was calculated and then converted by step function. To be specific, the component score of a linker was assigned with 1, if (1) the effective length was between 4 and 7, (2) the molecular weight was in the range from 70 to 200, (3) the number of aromatic rings was no greater than 1, or (4) the number of aliphatic rings was 0 or 1. Otherwise it was set to 0. If all the constraints were met, i.e. the ComboScore of this generated linker was 4, then the FinalScore was 1. If not all the constraints were met, i.e., the ComboScore was less than 4, the FinalScore was 0.

The input of Link-INVENT was the SMILES strings of two warheads, i.e., the symmetric terminal groups of **Cpd 4**. The ComboScore was set as the MPO objective of the Agent. The Agent proposed linkers/linked molecules in SMILES strings at a batch size of 128. The Agent was updated for 200 RL cycles, with a learning rate of 1×10⁻⁴.

##### 4.1.1.2. Building of antibacterial activity prediction model by AutoMolDesigner

The compounds with activity data for *A. baumannii* were collected from ChEMBL34 (https://www.ebi.ac.uk/chembl/). The compounds were uncharged and their SMILES strings were standardized. The compounds with exact MIC values, molecular weights less than 1500, and rotatable bonds no more than 20 were retained. For those compounds with multiple MIC values, the average MIC was used to represent its antibacterial activity. Each compound was labeled as active or inactive, based on whether its MIC (or average MIC) was ≤ 1 μg/mL.

AutoMolDesigner (V1.1) (https://zenodo.org/record/10097899) was used to build a classifier for anti-*A. baumannii* activity prediction. ^21^ To be specific, models were constructed with the ‘AutoGluon’ module of AutoMolDesigner (V1.1). The ‘AutoGluon’ module comprises two sub-modules: ‘Model Training’ and ‘Model Prediction’. In the ‘Model Training’ sub-module, all the five molecular descriptors/fingerprints implemented in the ‘Molecular Features’ section, i.e., RDKit_2D_norm, RDKit_2D, ECFP_4, FCFP_6, and MACCS, were selected. The ‘Model Quality Preset’ was set to ‘best quality’. Random split was chosen to partition the data set into two parts, with 80% of the data for model training and the other 20% for model evaluation. ‘ROC_AUC’ was selected as the ‘Evaluation Metric’ during model training. In the ‘Model Prediction’ sub-module, accuracy (ACC), area under the ROC curve (AUROC), Matthews correlation coefficient (MCC), and F1_score based on the test set were automatically calculated for each model. The model with the highest ACC was selected for the following virtual screening.

##### 4.1.1.3. Virtual screening of the generated chemical library

###### General

The workflow mainly includes molecular docking of the molecules from the virtual chemical library for potential binders of MsbA, and anti-*A. baumannii* activity prediction with the best-performing machine learning model, calculation of SA score, and binding mode analysis by visual inspection.

###### Molecular docking

The cyro-EM structure of MsbA in complex with Cerastecin C (PDB Entry: 8GK7) was obtained from PDB (https://www.rcsb.org/structure/8GK7). Cerastecin C was stripped from the cyro-EM structure and used to define the binding site. The MsbA protein was cleaned by the module of ‘Clean the Protein’ implemented in Discovery Studio (version 2018, BIOVIA, Inc). ^30^ OMEGA (version 2.5.1.4, OpenEye Scientific Software, Santa Fe, NM.) was used to generate all plausible poses of each molecule. ^31^ FRED (version 2.5.1.4, OpenEye Scientific Software, Santa Fe, NM.) was used for molecular docking, in which Chemgauss4 was the function to score docking poses. ^24, 25^ Only the top-scoring pose was saved for each molecule, and when its Chemgauss4 score was better than that of **Cpd 4,** this molecule was regarded as a potential MsbA binder.

###### Anti-A. baumannii activity prediction

The above mentioned machine learning model was used to predict whether those potential MsbA binders may have anti-*A. baumannii* activity, with probability of being active as the output. Herein, molecules with probability > 0.5 were selected as potential antibacterial molecules.

###### Synthetic accessibility calculation

The ‘SA score’ module implemented in AutoMolDesigner was used to measure synthetic accessibility of each molecule. ^27^ Top 10% molecules were saved for further analysis.

###### Visual inspection

The binding mode of each molecule was carefully inspected by using Discovery Studio (version 2018, BIOVIA, Inc). Then, molecules that could form two hydrogen bonds between each of two carboxyl groups and Arg-72 were kept as potential hits.

#### 4.1.2. Molecular dynamic simulation

The simulation environment of the protein-ligand complex was prepared using the ‘System Builder’ module. The POPC (300 K) membrane was embedded in the system, with its position determined by the trans-membrane amino acid residues of MsbA. The protein-ligand complex was solvated in an orthorhombic box using the SPC water model, with a buffer distance of 10 Å. To neutralize the system,7 sodium ions were added.

The simulation was performed using Desmond with the OPLS3 force field. ^32^ Prior to the production simulation, the system was submitted for a multi-stage equilibration simulation. This process included 100-ps Brownian Dynamics NVT (isothermal-isochoric) simulation, 12-ps NVT simulation, 48-ps of multi-step NPT (isothermal-isobaric) simulations. Restraints were applied to the heavy atoms of the protein during these equilibration steps except for the last 24 ps NPT simulation to gradually stabilize the system to 300K and 1.0 bar.

For the production simulation, the system was simulated for 100 ns, during which the temperature and pressure were kept at 300 K and 1.0 bar, respectively. Atom coordinates were recorded every 100 ps, resulting in a total of 1000 frames. The simulation trajectory was analyzed using the ‘Simulation Interaction Diagram’ module in Maestro.

### 4.2 Chemical synthesis

#### 4.2.1 Chemicals and instruments

All of the reagents were obtained from commercial sources and used without further purification. Thin-layer chromatography (TLC) on the silica gel plates GF254 (200–300 mm; Qingdao Haiyang Chemical Co., Ltd., Qingdao, China) with UV light illumination was used to monitor chemical reactions. ^1^H NMR (400, 500 or 700 MHz) and ^13^C NMR (101, 126 or 176 MHz) spectra were measured by Quantum-400 MHz NMR spectrometer (Q.One Instruments Ltd., China) and JEOL 500 MHz NMR spectrometer (JEOL Ltd., Japan). Chemical shifts were reported in *δ* values (ppm), with tetramethylsilane as the internal standard. High-resolution mass spectrometry (HRMS) was performed using the Thermo Scientific^TM^ Exactive^TM^ Plus mass spectrometer (Thermo, USA). The purity was determined by high-performance liquid chromatography (HPLC) of the Waters Acquity machine, with the Bridged Ethylene Hybrid (BEH) C18 column (1.7 μm, 50 × 2.1 mm), water (containing 0.1 % formic acid) as mobile phase A and acetonitrile as mobile phase B at the flow rate of 0.20 mL/min. All the compounds are >95% pure by HPLC analysis.

#### 4.2.2 Procedure for the synthesis of In-1

##### 4.2.2.1 2-(4-chlorophenyl)thiophene (intermediate C1)

Pd(PPh_3_)_4_ (11.6 mg, 0.01mmol) was added to a mixture of pinacol 4-chlorophenylborate (286 mg, 1.2 mmol), 2-iodothiophene (210 mg, 1.0 mmol), K_2_CO_3_ (207 mg, 1.5 mmol), 1,4-dioxane (5 mL) and water (1 mL) under N_2_. The reaction solution was heated and refluxed at 130 ℃ for 2 h, and the solvent was removed by rotary evaporation. Water (20 mL) was added and the product was extracted with EtOAc (3×10 mL). The combined organic phases were washed with saturated aqueous sodium chloride (10 mL), dried over anhydrous sodium sulfate, and filtered. The filtrate was concentrated under reduced pressure using a rotary evaporator. The crude product was purified by silica gradient column chromatography (PE) to afford **C1** (189 mg, 97.4%) as a white solid. ^1^H NMR (500 MHz, Chloroform-*d*) *δ* 7.55 – 7.54 (m, 1H), 7.53 – 7.52 (m, 1H), 7.36 – 7.35 (m, 1H), 7.35 – 7.33 (m, 1H), 7.31 – 7.28 (m, 2H), 7.08 (dd, *J* = 4.8, 3.9 Hz, 1H). ^13^C NMR (126 MHz, Chloroform-*d*) *δ* 143.2, 133.3, 133.1, 129.2, 128.3, 127.3, 125.3, 123.6.

##### 4.2.2.2 5-(4-chlorophenyl)thiophene-2-sulfonic acid (intermediate C2)

The concentrated sulfuric acid (58 μL, 1.0 mmol) was added to the mixture of 2-(4-chlorophenyl)thiophene (100 mg, 0.5 mmol), EtOAc (2 mL), acetic anhydride (97 μL, 1.0 mmol) drop by drop in an ice bath. The mixture was stirred at room temperature for 24 h. Saturated brine (2 mL) and EtOAc (2 mL) were added to the reaction mixture, and solid was observed to precipitate from the organic layer. The organic layer was separated and filtered to afford a white solid, which was dried to obtain **C2** (172 mg, 101.3%) as a white solid. ^1^H NMR (500 MHz, D_2_O) *δ* 7.60 (d, *J* = 8.7 Hz, 2H), 7.45 (d, *J* = 3.9 Hz, 1H), 7.43 (d, *J* = 8.5 Hz, 2H), 7.30 (d, *J* = 3.9 Hz, 1H). ^13^C NMR (126 MHz, D_2_O) *δ* 223.9, 146.4, 142.7, 133.9, 131.4, 130.0, 129.1, 127.3, 123.6.

##### 4.2.2.3 5-(4-chlorophenyl)thiophene-2-sulfonyl chloride (intermediate C3)

A drop of DMF was added to the mixture of **C2** (400 mg, 0.7 mmol) and SOCl_2_ (5 mL). The reaction solution was heated and refluxed at 85℃ for 8 h, cooled to room temperature, and slowly poured into crushed ice. The product was extracted with DCM (3×10 mL). The combined organic phases were washed with saturated aqueous sodium chloride (10 mL), dried over anhydrous sodium sulfate, and filtered. The filtrate was concentrated under reduced pressure to afford **C3** (129.6 mg, 60.6%) as a yellow solid. ^1^H NMR (500 MHz, Chloroform-*d*) *δ* 7.84 (d, *J* = 4.1 Hz, 1H), 7.58 – 7.54 (m, 2H), 7.46 – 7.42 (m, 2H), 7.29 (d, *J* = 4.1 Hz, 1H). ^13^C NMR (126 MHz, Chloroform-*d*) *δ* 154.1, 142.2, 136.5, 135.9, 130.4, 129.9, 127.9, 123.6.

##### 4.2.2.4 Methyl 2-((5-(4-chlorophenyl)thiophene)-2-sulfonamido)-5-iodobenzoate (intermediate **C4**)

Pyridine (158 mg, 2.0 mmol) was added to the mixture of methyl 2-amino-5-iodobenzoate (277 mg, 1.0 mmol) and DCM (5 mL). **C3** (441 mg, 1.5 mmol) was dissolved in DCM (5 mL), and then added drop by drop into the reaction system. The reaction solution was stirred at room temperature for 12 h, and then HCl (5 mL, 1 mol/L) was added. The product was extracted with DCM (3×10 mL). The combined organic phases were washed with saturated aqueous sodium chloride (10 mL), dried over anhydrous sodium sulfate, and filtered. The filtrate was concentrated under reduced pressure. The crude product was purified by silica gradient column chromatography (PE/EA = 5:1) to afford **C4** (532 mg, 98.8%). ^1^H NMR (500 MHz, Chloroform-*d*) *δ* 10.73 (s, 1H), 8.27 (d, *J* = 2.2 Hz, 1H), 7.79 (dd, *J* = 8.8, 2.2 Hz, 1H), 7.60 – 7.55 (m, 2H), 7.46 (d, *J* = 8.6 Hz, 2H), 7.37 (d, *J* = 8.6 Hz, 2H), 7.15 (d, *J* = 3.9 Hz, 1H), 3.90 (s, 3H). ^13^C NMR (126 MHz, Chloroform-*d*) *δ* 167.2, 150.7, 143.3, 140.0, 138.4, 135.5, 134.1, 131.1, 129.6, 127.6, 123.4, 120.9, 117.8, 86.1, 53.0.

##### 4.2.2.5 3-(4-((5-(4-chlorophenyl)thiophene)-2-sulfonamido)-3-(methoxycarbonyl)phenyl)acrylic acid (intermediate **C5**)

Pd(dppf)Cl_2_ (140 mg, 0.2 mmol) was added to the mixture of **C4** (500 mg, 1.0 mmol), acrylic acid (330 μL, 4.8 mmol), triethylamine (1.34 mL) and DMF (10 mL) under N_2_. After the reaction solution was heated at 100℃ for 2 h, HCl (10mL, 1 mol/L) was added. The product was extracted with EtOAc (3×10 mL). The combined organic phases were washed with saturated aqueous sodium chloride (10 mL), dried over anhydrous sodium sulfate, and filtered. The filtrate was concentrated under reduced pressure. The crude product was purified by silica gradient column chromatography (PE/ EtOAc = 1:1) to afford **C5** (245 mg, 53.5%) as a white solid. ^1^H NMR (500 MHz, DMSO-*d*_6_) *δ* 8.07 (s, 1H), 7.93 (d, *J* = 7.5 Hz, 1H), 7.72 (d, *J* = 8.6 Hz, 2H), 7.68 (d, *J* = 3.4 Hz, 1H), 7.57 (d, *J* = 4.3 Hz, 1H), 7.55 (d, *J* = 3.7 Hz, 1H), 7.54 – 7.47 (m, 3H), 6.49 (d, *J* = 16.0 Hz, 1H), 3.83 (s, 3H). ^13^C NMR (126 MHz, DMSO-*D*_6_) *δ* 167.3, 142.0, 133.9, 131.0, 130.7, 129.4, 127.8, 124.8, 120.7, 62.8, 52.7.

##### 4.2.2.6 3-(4-((5-(4-chlorophenyl)thiophene)-2-sulfonamido)-3-(methoxycarbonyl)phenyl)propanoic acid (intermediate **C6**)

Dry Pd/C (50 mg) was added to the mixture of **C5** (477 mg, 1.0 mmol), DCM (5 mL) and MeOH (5 mL). The reaction solution was put it into a hydrogenation reactor (H_2_: 0.4 MPa) and stirred for 4 h. Then, the reaction solution was filtered. The filtrate was concentrated under reduced pressure using a rotary evaporator. The crude product was purified by silica gradient column chromatography (PE/EtOAc = 1:1) to afford **C6** (424 mg, 88.5%) as a white solid. ^1^H NMR (500 MHz, Pyridine-*d*_5_) *δ* 8.08 (d, *J* = 8.5 Hz, 1H), 7.96 (d, *J* = 2.2 Hz, 1H), 7.83 (d, *J* = 4.0 Hz, 1H), 7.61 (dd, *J* = 8.6, 2.3 Hz, 1H), 7.48 (d, *J* = 8.6 Hz, 2H), 7.41 – 7.33 (m, 2H), 7.29 (d, *J* = 4.0 Hz, 1H), 3.74 (s, 3H), 3.06 (t, *J* = 7.5 Hz, 2H), 2.82 (t, *J* = 7.5 Hz, 2H). ^13^C NMR (126 MHz, Pyridine-*d*_5_) *δ* 175.3, 168.9, 139.7, 138.4, 138.1, 135.5, 135.4, 134.9, 131.9, 131.7, 130.0, 128.3, 126.8, 124.8, 121.2, 118.4, 52.9, 36.3, 30.9.

##### 4.2.2.7 Methyl 3’-amino-4-((5-(4-chlorophenyl)thiophene)-2-sulfonamido)-[1,1’-biphenyl]-3-carboxylate (intermediate **C7**)

Pd(PPh_3_)_4_ (115 mg, 0.1 mmol) was added to the mixture of **C4** (533 mg, 1.0 mmol), 3-aminophenylboronic acid (205 mg, 1.5 mmol), K_2_CO_3_ (415 mg, 3.0 mmol), 1,4-dioxane (10 mL) and water (2 mL) under N_2_. After the reaction solution was heated at 60 ℃ for 2 h, HCl (10 mL, 1 mol/L) was added. The product was extracted with EtOAc (3×10 mL). The combined organic phases were washed with saturated aqueous sodium chloride (10 mL), dried over anhydrous sodium sulfate, and filtered. The filtrate was concentrated under reduced pressure. The crude product was purified by silica gradient column chromatography (PE/ EtOAc = 1:1) to afford **C7** (282 mg, 56.6%) as a white solid. ^1^H NMR (500 MHz, DMSO-*d*_6_) *δ* 10.54 (s, 1H), 8.01 (d, *J* = 2.3 Hz, 1H), 7.84 (dd, *J* = 8.6, 2.3 Hz, 1H), 7.72 (d, *J* = 8.3 Hz, 2H), 7.67 (d, *J* = 4.0 Hz, 1H), 7.63 (d, *J* = 8.6 Hz, 1H), 7.58 (d, *J* = 4.0 Hz, 1H), 7.50 (d, *J* = 8.2 Hz, 2H), 7.10 (t, *J* = 7.8 Hz, 1H), 6.82 (s, 1H), 6.76 (d, *J* = 7.6 Hz, 1H), 6.58 (d, *J* = 8.0 Hz, 1H), 3.83 (s, 3H). ^13^C NMR (126 MHz, DMSO-*d*_6_) *δ* 167.4, 149.1, 149.0, 138.7, 137.8, 137.2, 136.6, 134.5, 134.0, 132.0, 130.7, 129.7, 129.4, 128.4, 127.8, 124.8, 121.7, 119.8, 114.1, 113.8, 111.8, 52.8.

##### 4.2.2.8 3’-(3-(3-carboxy-4-((5-(4-chlorophenyl)thiophene)-2-sulfonamido)phenyl)propanamido)-4-((5-(4-chlorophenyl)thiophene)-2-sulfonamido)-[1,1’-biphenyl]-3-carboxylic acid (**In-1**)

TCFH (420 mg, 1.5 mmol) was added to the mixture of **C6** (479 mg, 1.0 mmol), **C7** (498 mg, 1.0 mmol), DCM (10 mL), 1-methylimidazole (NMI) (287 mg, 3.5 mmol). After the reaction solution was stirred at room temperature for 12 h, water (20 mL) was added. The product was extracted with DCM (3×10 mL). The combined organic phases were washed with saturated aqueous sodium chloride (10 mL), dried over anhydrous sodium sulfate, and filtered. After the filtrate was concentrated under reduced pressure to afford the intermediate, THF (10 mL), MeOH (2 mL) and LiOH (1 mL, 5.5 mol/L) were added. After the reaction solution was stirred at room temperature for 12h, HCl (20mL, 1 mol/L) was added to adjust the pH to acidic. The product was extracted with EtOAc (3×10 mL). The combined organic phases were washed with saturated aqueous sodium chloride (10 mL), dried over anhydrous sodium sulfate, and filtered. The filtrate was concentrated under reduced pressure. The crude product was purified by silica gradient column chromatography (DCM/MeOH = 10:1) to afford **In-1** (490 mg, 52.7%) as a white solid. ^1^H NMR (700 MHz, DMSO-*d*_6_) *δ* 10.14 (s, 1H), 8.13 (d, *J* = 2.4 Hz, 1H), 7.82 – 7.81 (m, 2H), 7.72 – 7.69 (m, 1H), 7.68 (d, *J* = 8.6 Hz, 2H), 7.65 (d, *J* = 8.5 Hz, 2H), 7.61 (d, *J* = 8.7 Hz, 1H), 7.58 (d, *J* = 3.9 Hz, 1H), 7.56 – 7.53 (m, 2H), 7.50 (d, *J* = 4.0 Hz, 1H), 7.48 (d, *J* = 5.1 Hz, 2H), 7.46 (s, 2H), 7.45 (d, *J* = 3.3 Hz, 2H), 7.43 (d, *J* = 9.7 Hz, 1H), 7.31 (t, *J* = 7.9 Hz, 1H), 7.25 (d, *J* = 8.2 Hz, 1H), 2.86 (t, *J* = 7.7 Hz, 2H), 2.59 (t, *J* = 7.7 Hz, 2H). ^13^C NMR (176 MHz, DMSO-*d*_6_) *δ* 170.4, 169.6, 169.2, 139.8, 139.3, 133.7, 133.5, 131.1, 130.9, 130.9, 129.3, 129.3, 129.3, 128.8, 127.7, 127.6, 125.9, 124.5, 124.4, 120.8, 118.8, 118.8, 117.9, 116.6, 37.8, 29.9. HRMS calcd for C_43_H_11_Cl_2_N_3_O_9_S_4_, [M-H]^-^, 930.0274; found, 930.0101.

#### 4.2.3 Procedures for the synthesis of Y-1∼Y-17

##### 4.2.3.1 Ethyl -2-amino-5-(3-(tert-butoxy)-3-oxoprop-1-en-1-yl)benzoate (intermediate **C8**)

Pd(dppf)Cl_2_ (500 mg, 0.7 mmol) was added to the mixture of ethyl 2-amino-5-iodobenzoate (2.0 g, 6.9 mmol), tert-butyl acrylate (1.8 g, 13.7 mmol), TEA (1.9 mL, 13.7 mmol), DMF (20 mL) under N_2_. After the reaction solution was heated at 100℃ for 2 h, HCl (10mL, 1 mol/L) was added. The product was extracted with EtOAc (3×20 mL). The combined organic phases were washed with saturated aqueous sodium chloride (20 mL), dried over anhydrous sodium sulfate, and filtered. The filtrate was concentrated under reduced pressure. The crude product was purified by silica gradient column chromatography (PE/EtOAc = 5:1) to afford **C8** (1.2 g, 60.0%) as a colorless oil. ^1^H NMR (500 MHz, Chloroform-*d*) *δ* 8.03 (d, *J* = 2.2 Hz, 1H), 7.49 (d, *J* = 15.9 Hz, 1H), 7.45 (dd, *J* = 8.6, 2.2 Hz, 1H), 6.64 (d, *J* = 8.6 Hz, 1H), 6.18 (d, *J* = 15.8 Hz, 1H), 4.35 (q, *J* = 7.1 Hz, 2H), 1.53 (s, 9H), 1.40 (t, *J* = 7.1 Hz, 3H). ^13^C NMR (126 MHz, Chloroform-*d*) *δ* 167.9, 167.0, 151.8, 143.3, 133.0, 132.5, 123.1, 117.2, 116.3, 110.9, 80.2, 60.8, 28.4, 14.5.

##### 4.2.3.2 Ethyl 2-amino-5-(3-(tert-butoxy)-3-oxopropyl)benzoate (intermediate **C9**)

Dry Pd/C (120 mg) was added to the mixture of **C8** (1.2 g, 4.1 mmol), DCM (6 mL) and MeOH (6 mL). Then, the mixture was put into a hydrogenation reactor (H_2_: 0.4 MPa) and stirred for 4 h. The reaction solution was filtered. The filtrate was concentrated under reduced pressure to afford **C9** (1.0 g, 85.0%) as green oil. ^1^H NMR (500 MHz, Chloroform-*d*) *δ* 7.69 (d, *J* = 2.5 Hz, 1H), 7.11 (dd, *J* = 8.4, 1.8 Hz, 1H), 6.60 (d, *J* = 8.3 Hz, 1H), 4.32 (q, *J* = 7.2 Hz, 2H), 2.79 (t, *J* = 7.7 Hz, 2H), 2.48 (t, *J* = 7.1 Hz, 2H), 1.42 (s, 9H), 1.38 (t, *J* = 7.1 Hz, 3H). ^13^C NMR (126 MHz, Chloroform-*d*) *δ* 172.4, 168.2, 149.0, 134.5, 130.6, 128.6, 117.1, 111.1, 80.4, 60.4, 37.5, 30.3, 28.2, 14.5.

##### 4.2.3.3 Ethyl 4-amino-3’-((tert-butoxycarbonyl)amino)-[1,1’-biphenyl]-3-carboxylate (intermediate **C10**)

Pd(PPh_3_)_4_ (0.8 g, 0.7 mmol) was added to the mixture of ethyl 2-amino-5-iodobenzoate (2.0 g, 6.9 mmol), (3-((tert-butoxycarbonyl) amino) phenyl) boronic acid (1.8 g, 7.6 mmol), K_2_CO_3_ (1.9 g, 13.7 mmol), 1,4-dioxane (20 mL) and water (4 mL) under N_2_. After the reaction solution was heated at 60℃ for 2 h, HCl (20 mL, 1 mol/L) was added. The product was extracted with EtOAc (3×20 mL). The combined organic phases were washed with saturated aqueous sodium chloride (20 mL), dried over anhydrous sodium sulfate, and filtered. The filtrate was concentrated under reduced pressure. The crude product was purified by silica gradient column chromatography (PE/EtOAc = 1:1) to afford **C10** (2.1 g, 90.0%) as a white solid. ^1^H NMR (500 MHz, Chloroform-*d*) *δ* 8.10 (d, *J* = 2.3 Hz, 1H), 7.52 (dd, *J* = 8.6, 2.2 Hz, 1H), 7.50 (s, 1H), 7.32 (d, *J* = 5.6 Hz, 1H), 7.24 – 7.19 (m, 1H), 6.72 (d, *J* = 8.5 Hz, 1H), 6.54 (s, 1H), 4.37 (q, *J* = 7.1 Hz, 2H), 1.54 (s, 9H), 1.40 (t, *J* = 7.1 Hz, 3H). ^13^C NMR (126 MHz, Chloroform-*d*) *δ* 168.3, 150.0, 141.6, 138.9, 133.0, 129.7, 129.5, 129.2, 121.3, 117.3, 111.3, 60.6, 28.5, 14.6.

##### 4.2.3.4 Ethyl 4-((5-bromothiophene)-2-sulfonamido)-3’-(3-(4-((5-bromothiophene)-2-sulfonamido)-3-(ethoxycarbonyl)phenyl)propanamido)-[1,1’-biphenyl]-3-carboxylate (intermediate **C11**)

Pyridine(540 μL, 6.8 mmol) was added to the mixture of **C9** (1.0 g, 3.4 mmol) and DCM(10 mL). 5-bromothiophene-2-sulfonyl chloride (1.3 g, 5.1 mmol) was dissolved in DCM (10 mL), added drop by drop into the reaction system. After the reaction solution was stirred at room temperature for 12 h, HCl (10mL, 1 mol/L) was added. The product was extracted with EtOAc (3×20 mL). The combined organic phases were washed with saturated aqueous sodium chloride (20 mL), dried over anhydrous sodium sulfate, and filtered. The filtrate was concentrated under reduced pressure to afford the intermediate, ethyl 2-((5-bromothiophene)-2-sulfonamido)-5-(3-(tert-butoxy)-3-oxopropyl)benzoate (1.6 g, 94.6%).

The above intermediate (1.6 g, 3.3 mmol) was mixed with DCM (16 mL), and then TFA (10 mL) was added to the mixture. The reaction solution was stirred at room temperature for 4 h and then concentrated under reduced pressure to afford the intermediate, 3-(4-((5-bromothiophene)-2-sulfonamido)-3-(ethoxycarbonyl)phenyl)propanoic acid (1.5 g, 98.5%).

Pyridine (90 µL, 1.1 mmol) was added to the mixture of **C10** (260 mg, 0.7 mmol) and DCM (5 mL). 5-bromothiophene-2-sulfonyl chloride (285 mg, 1.1 mmol) was dissolved in DCM (5 mL), added drop by drop into the reaction mixture, and the reaction solution was stirred at room temperature for 12 h. The reaction solution was concentrated under reduced pressure to afford the intermediate, ethyl 4-((5-bromothiophene)-2-sulfonamido)-3’-((tert-butoxycarbonyl)amino)-[1,1’-biphenyl]-3-carboxylate (369 mg, 93.2%).

This intermediate (369 mg, 0.6 mmol) was mixed with 1,4-dioxane (5 mL), and then HCl (5 mL, 4M in 1,4-dioxane) was added. The reaction solution was stirred at room temperature for 4 h. The reaction solution was concentrated under reduced pressure to afford the intermediate, ethyl 3’-amino-4-((5-bromothiophene)-2-sulfonamido)-[1,1’-biphenyl]-3-carboxylate (269 mg, 96.2%).

TCFH (613 mg, 2.2 mmol) was added to the mixture of ethyl 3’-amino-4-((5-bromothiophene)-2-sulfonamido)-[1,1’-biphenyl]-3-carboxylate (700mg, 1.5 mmol), 3-(4-((5-bromothiophene)-2-sulfonamido)-3-(ethoxycarbonyl)phenyl)propanoic acid (672 mg, 1.5 mmol), DCM(10 mL), NMI (360 µL, 4.5 mmol). After the reaction solution was stirred at room temperature for 12 h, water (20 mL) was added. The product was extracted with DCM (3×20 mL). The combined organic phases were washed with saturated aqueous sodium chloride (20 mL), dried over anhydrous sodium sulfate, and filtered. The filtrate was concentrated under reduced pressure. The crude product was purified by silica gradient column chromatography (PE/ EtOAc = 1:1) to afford **C11** (1.2 g, 90.0%) as a white solid. ^1^H NMR (500 MHz, Chloroform-*d*) *δ* 10.83 (s, 1H), 10.68 (s, 1H), 8.16 (d, *J* = 2.2 Hz, 1H), 7.86 (d, *J* = 2.2 Hz, 1H), 7.79 (d, *J* = 8.6 Hz, 1H), 7.74 (t, *J* = 1.9 Hz, 1H), 7.70 (dd, *J* = 8.7, 2.3 Hz, 1H), 7.65 (d, *J* = 8.5 Hz, 1H), 7.44 (d, *J* = 8.1 Hz, 1H), 7.41 – 7.39 (m, 1H), 7.37 (d, *J* = 2.7 Hz, 1H), 7.35 (d, *J* = 4.0 Hz, 1H), 7.29 – 7.26 (m, 2H), 6.98 (d, *J* = 4.0 Hz, 1H), 6.91 (d, *J* = 4.0 Hz, 1H), 4.38 (q, *J* = 7.1 Hz, 2H), 4.31 (q, *J* = 7.1 Hz, 2H), 3.04 (t, *J* = 7.5 Hz, 2H), 2.68 (t, *J* = 7.6 Hz, 2H), 1.40 (t, *J* = 7.1 Hz, 3H), 1.34 (t, *J* = 7.1 Hz, 3H).

##### 4.2.3.5 General procedures for the synthesis of Y-1∼Y-17

Pd(PPh_3_)_4_ (0.1 eq) was added to the mixture of **C11**(1.0 eq), substituted arylboronic acid (1.5 eq), K_2_CO_3_ (3.0 eq), 1,4-dioxane (10 V), water (2 V) under N_2_. After the reaction solution was heated at 60℃ for 2 h, HCl (10 V, 1 mol/L) was added. The product was extracted with EtOAc (3×10 V). The combined organic phases were washed with saturated aqueous sodium chloride (10 V), dried over anhydrous sodium sulfate, and filtered. The filtrate was concentrated under reduced pressure. THF (10 V), MeOH (2 V), LiOH (1 V, 5.5 mol/L) were added. After the reaction solution was stirred at room temperature for 12h, HCl (10 V, 1 mol/L) was added to make the solution acidic. The product was extracted with EtOAc (3×10 V). The combined organic phases were washed with saturated aqueous sodium chloride (10 V), dried over anhydrous sodium sulfate, and filtered. The filtrate was concentrated under reduced pressure. The crude product was purified by silica gradient column chromatography (DCM/MeOH = 10:1) to afford **Y-1∼Y-17**.

###### 3’-(3-(3-carboxy-4-((5-phenylthiophene)-2-sulfonamido)phenyl)propanamido)-4-((5-phenylthiophene)-2-sulfonamido)-[1,1’-biphenyl]-3-carboxylic acid (Y-1)

Yield 62%, white solid. ^1^H NMR (500 MHz, DMSO-*d*_6_) *δ* 9.96 (s, 1H), 8.17 (d, *J* = 2.4 Hz, 1H), 7.81 (d, *J* = 2.2 Hz, 2H), 7.65 – 7.63 (m, 2H), 7.62 – 7.59 (m, 3H), 7.57 (d, *J* = 8.6 H z, 1H), 7.51 – 7.48 (m, 2H), 7.44 (dd, *J* = 6.5, 3.4 Hz, 2H), 7.43 – 7.40 (m, 3H), 7.40 – 7.37 (m, 3H), 7.37 – 7.33 (m, 2H), 7.32 – 7.26 (m, 2H), 7.24 (d, *J* = 7.9 H z, 1H), 2.83 (t, *J* = 7.7 Hz, 2H), 2.56 (t, 2H). ^13^C NMR (126 MHz, DMSO-*d*_6_) *δ* 1 70.5, 169.9, 169.4, 148.3, 147.9, 139.9, 139.8, 132.5, 132.4, 132.3, 132.0, 131.6, 130. 7, 130.3, 129.3, 129.3, 128.9, 128.8, 128.8, 125.8, 123.6, 123.5, 120.9, 120.7, 118.2, 117.9, 117.6, 116.6, 38.1, 30.1. HRMS calcd for C_43_H_33_N_3_O_9_S_4_ [M-H]^-^, 862.1027; found, 862.0803.

###### 3’-(3-(3-carboxy-4-((5-(4-fluorophenyl)thiophene)-2-sulfonamido)phenyl)propanamido)-4-((5-(4-fluorophenyl)thiophene)-2-sulfonamido)-[1,1’-biphenyl]-3-carboxylic acid (Y-2)

Yield 54%, white solid. ^1^H NMR (500 MHz, DMSO-*d*_6_) *δ* 10.08 (s, 1H), 8.14 (s, 1H), 7.81 (s, 2H), 7.72 – 7.66 (m, 5H), 7.62 (d, *J* = 8.6 Hz, 1H), 7.57 (d, *J* = 3.9 Hz, 1 H), 7.56 – 7.52 (m, 2H), 7.49 (d, *J* = 8.5 Hz, 1H), 7.44 – 7.42 (m, 2H), 7.39 (d, *J* = 3.9 Hz, 1H), 7.31 (t, *J* = 7.9 Hz, 1H), 7.27 – 7.22 (m, 5H), 2.86 (t, *J* = 7.6 Hz, 2H), 2.59 (t, *J* = 7.7 Hz, 2H). ^13^C NMR (126 MHz, DMSO-*d*_6_) *δ* 172.0, 170.4, 163. 5, 161.5, 161.4, 139.8, 139.3, 133.8, 133.5, 131.1, 130.9, 129.3, 128.8, 128.8, 128.6, 128.6, 128.2, 128.2, 128.1, 124.0, 123.9, 120.8, 118.8, 118.7, 117.9, 116.6, 116.4, 116. 3, 116.2, 116.2, 37.8, 29.9. HRMS calcd for C_43_H_31_F_2_N_3_O_9_S_4_ [M-H]^-^, 898.0838; found, 898.0850.

###### 3’-(3-(3-carboxy-4-((5-(4-nitrophenyl)thiophene)-2-sulfonamido)phenyl)propanamido)-4-((5-(4-nitrophenyl)thiophene)-2-sulfonamido)-[1,1’-biphenyl]-3-carboxylic acid (Y-3)

Yiel d 60%, white solid. ^1^H NMR (500 MHz, DMSO-*d*_6_) *δ* 9.96 (s, 1H), 8.23 – 8.20 (m, 4H), 8.15 (d, *J* = 2.4 Hz, 1H), 7.91 (dd, *J* = 15.7, 8.8 Hz, 4H), 7.82 – 7.79 (m, 2 H), 7.68 (d, *J* = 4.0 Hz, 1H), 7.65 – 7.62 (m, 2H), 7.60 – 7.57 (m, 2H), 7.54 (d, *J* = 4.0 Hz, 1H), 7.50 (d, *J* = 8.5 Hz, 1H), 7.45 (d, *J* = 8.4 Hz, 1H), 7.34 – 7.28 (m, 2H), 7.24 (d, *J* = 7.8 Hz, 1H), 2.84 (t, *J* = 7.7 Hz, 2H), 2.57 (t, *J* = 7.7 Hz, 2H). ^13^C NMR (126 MHz, DMSO-*d*_6_) *δ* 170.4, 169.6, 169.2, 147.0, 146.9, 145.5, 144.9, 1 39.8, 139.7, 138.6, 138.4, 133.2, 132.4, 131.8, 130.8, 130.7, 129.3, 128.8, 126.8, 126. 7, 126.6, 124.5, 120.7, 120.3, 119.6, 118.6, 118.5, 117.7, 116.6, 38.0, 30.0. HRMS calcd for C_43_H_31_N_5_O_13_S_4_ [M-H]^-^, 952.0728; found, 952.0474.

###### 3’-(3-(3-carboxy-4-((5-(4-hydroxyphenyl)thiophene)-2-sulfonamido)phenyl)propanamido) -4-((5-(4-hydroxyphenyl)thiophene)-2-sulfonamido)-[1,1’-biphenyl]-3-carboxylic acid (Y-4)

Yield 48%, white solid. ^1^H NMR (500 MHz, DMSO-*d*_6_) *δ* 10.00 (s, 1H), 9.92 (s, 2 H), 8.15 (s, 1H), 7.88 – 7.81 (m, 3H), 7.71 (d, *J* = 8.7 Hz, 1H), 7.65 (d, *J* = 4.0 Hz, 1H), 7.57 – 7.52 (m, 4H), 7.48 (dd, *J* = 16.1, 8.2 Hz, 5H), 7.36 – 7.29 (m, 3 H), 7.25 (d, *J* = 3.8 Hz, 1H), 6.80 (d, *J* = 6.6 Hz, 4H), 2.88 (t, *J* = 7.5 Hz, 2H), 2. 60 (t, *J* = 7.7 Hz, 2H). ^13^C NMR (126 MHz, DMSO-*d*_6_) *δ* 170.4, 163.1, 158.7, 158. 6, 139.8, 139.5, 133.6, 130.9, 129.3, 127.4, 127.4, 123.1, 122.9, 121.8, 121.7, 120.8, 118.5, 117.9, 116.7, 116.1, 116.1, 37.9, 30.0. HRMS calcd for C_43_H_33_N_3_O_11_S_4_ [M-H]^-^, 894.0925; found, 894.0858.

###### 3’-(3-(3-carboxy-4-((5-(4-methoxyphenyl)thiophene)-2-sulfonamido)phenyl)propanamido)-4-((5-(4-methoxyphenyl)thiophene)-2-sulfonamido)-[1,1’-biphenyl]-3-carboxylic acid (Y-5)

Yield 50%, white solid. ^1^H NMR (500 MHz, DMSO-*d*_6_) *δ* 9.96 (s, 1H), 8.17 (d, *J* = 2.4 Hz, 1H), 7.82 (s, 2H), 7.62 (dd, *J* = 8.6, 2.4 Hz, 1H), 7.58 – 7.55 (m, 3H), 7.55 – 7.52 (m, 2H), 7.49 (d, *J* = 7.1 Hz, 1H), 7.47 (d, *J* = 3.9 Hz, 1H), 7.45 – 7. 41 (m, 2H), 7.32 – 7.27 (m, 3H), 7.25 – 7.23 (m, 2H), 6.98 – 6.94 (m, 4H), 3.76 (d, *J* = 1.5 Hz, 6H), 2.83 (t, *J* = 7.7 Hz, 2H), 2.56 (t, 2H). ^13^C NMR (126 MHz, D MSO-*d*_6_) *δ* 170.6, 170.2, 169.7, 160.0, 160.0, 148.9, 148.4, 140.1, 139.9, 132.7, 132.5, 132.0, 131.9, 130.9, 130.4, 129.4, 129.0, 127.4, 127.4, 125.2, 125.0, 122.4, 122.4, 12 0.9, 118.3, 118.1, 117.8, 116.8, 114.8, 55.5, 55.5, 38.3, 30.3. HRMS calcd for C_45_H_37_ N_3_O_11_S_4_ [M-H]^-^, 922.1238; found, 922.1220.

###### 3’-(3-(3-carboxy-4-((5-(3-fluorophenyl)thiophene)-2-sulfonamido)phenyl)propanamido)-4-((5-(3-fluorophenyl)thiophene)-2-sulfonamido)-[1,1’-biphenyl]-3-carboxylic acid (Y-6)

Y ield 58%, white solid. ^1^H NMR (500 MHz, DMSO-*d*_6_) *δ* 9.96 (s, 1H), 8.16 (d, *J* = 2. 4 Hz, 1H), 7.81 (s, 2H), 7.62 (dd, *J* = 8.6, 2.4 Hz, 1H), 7.57 (d, *J* = 8.5 Hz, 2H), 7.54 – 7.53 (m, 1H), 7.52 – 7.50 (m, 3H), 7.50 – 7.48 (m, 1H), 7.48 (s, 2H), 7.47 (s, 1H), 7.45 (dd, *J* = 3.1, 1.6 Hz, 1H), 7.44 – 7.43 (m, 2H), 7.42 (s, 1H), 7.32 – 7. 27 (m, 2H), 7.24 (d, *J* = 7.9 Hz, 1H), 7.21 – 7.16 (m, 2H), 2.83 (t, *J* = 7.7 Hz, 2 H), 2.56 (t, 2H). ^13^C NMR (126 MHz, DMSO-*d*_6_) *δ* 170.6, 170.0, 169.5, 163.7, 161. 8, 140.0, 139.9, 134.9, 134.8, 134.7, 134.7, 132.9, 132.2, 131.7, 131.6, 131.6, 131.5, 131.5, 130.9, 130.6, 129.4, 129.0, 124.9, 124.8, 122.2, 122.1, 120.9, 118.4, 118.2, 117. 7, 116.7, 115.9, 115.7, 115.7, 115.6, 112.8, 112.8, 112.6, 112.6, 38.2, 30.2. HRMS calcd for C_43_H_31_F_2_N_3_O_9_S_4_ [M-H]^-^, 898.0838; found, 898.0650.

###### 3’-(3-(3-carboxy-4-((5-(3-chlorophenyl)thiophene)-2-sulfonamido)phenyl)propanamido)-4-((5-(3-chlorophenyl)thiophene)-2-sulfonamido)-[1,1’-biphenyl]-3-carboxylic acid (Y-7)

Yield 61%, white solid. ^1^H NMR (500 MHz, DMSO-*d*_6_) *δ* 9.96 (s, 1H), 8.16 (d, *J* = 2.4 Hz, 1H), 7.80 (d, *J* = 5.0 Hz, 2H), 7.72 (d, *J* = 13.7 Hz, 2H), 7.60 – 7.59 (m, 1H), 7.58 (d, *J* = 2.3 Hz, 1H), 7.56 – 7.54 (m, 2H), 7.52 (d, *J* = 3.9 Hz, 1H), 7.5 0 – 7.48 (m, 2H), 7.47 (d, *J* = 3.9 Hz, 1H), 7.45 – 7.41 (m, 2H), 7.41 – 7.40 (m, 2H), 7.40 – 7.38 (m, 2H), 7.29 (t, *J* = 7.9 Hz, 1H), 7.24 – 7.19 (m, 2H), 2.81 (t, 2 H), 2.55 (t, 2H). ^13^C NMR (126 MHz, DMSO-*D*_6_) *δ* 170.5, 169.9, 169.4, 145.6, 145. 5, 144.9, 144.2, 143.7, 142.6, 140.0, 139.8, 134.5, 134.5, 134.0, 133.6, 132.7, 132.6, 132.1, 131.6, 131.5, 131.3, 131.2, 131.2, 131.1, 131.1, 130.6, 130.2, 129.9, 129.2, 12 8.8, 128.8, 128.7, 128.4, 128.4, 125.3, 125.2, 124.7, 124.6, 124.5, 124.5, 120.9, 120.6, 118.1, 117.6, 117.5, 116.5, 38.2, 30.2. HRMS calcd for C_43_H_31_Cl_2_N_3_O_9_S_4_ [M-H]-, 93 0.0247; found, 930.0244.

###### 3’-(3-(3-carboxy-4-((5-(3-nitrophenyl)thiophene)-2-sulfonamido)phenyl)propanamido)-4 -((5-(3-nitrophenyl)thiophene)-2-sulfonamido)-[1,1’-biphenyl]-3-carboxylic acid (Y-8)

Yiel d 54%, white solid. ^1^H NMR (500 MHz, DMSO-*d*_6_) *δ* 9.97 (s, 1H), 8.39 (t, *J* = 2.0 Hz, 1H), 8.36 (t, *J* = 2.1 Hz, 1H), 8.18 – 8.14 (m, 3H), 8.10 – 8.07 (m, 1H), 8.06 – 8.03 (m, 1H), 7.81 – 7.78 (m, 2H), 7.70 (d, *J* = 8.3 Hz, 1H), 7.66 (dd, *J* = 6.1, 2.2 Hz, 2H), 7.63 – 7.60 (m, 2H), 7.57 (d, *J* = 8.6 Hz, 1H), 7.54 (d, *J* = 3.9 Hz, 1 H), 7.51 – 7.48 (m, 2H), 7.43 (d, *J* = 8.4 Hz, 1H), 7.31 – 7.27 (m, 2H), 7.23 (dt, *J* = 7.8, 1.4 Hz, 1H), 2.82 (t, 2H), 2.56 (t, 2H). ^13^C NMR (126 MHz, DMSO-*d*_6_) *δ* 1 70.5, 169.6, 169.2, 148.4, 148.4, 144.7, 139.8, 139.8, 134.0, 133.9, 132.8, 132.1, 131. 9, 131.5, 130.9, 130.7, 130.5, 129.3, 128.7, 125.7, 125.7, 123.2, 123.1, 120.7, 120.5, 120.1, 120.0, 118.4, 118.2, 117.6, 116.5, 38.1, 30.1. HRMS calcd for C_43_H_31_N_5_O_13_S_4_ [M-H]^-^, 952.0728; found, 952.0712.

###### 3’-(3-(3-carboxy-4-((5-(3-hydroxyphenyl)thiophene)-2-sulfonamido)phenyl)propanamido)-4-((5-(3-hydroxyphenyl)thiophene)-2-sulfonamido)-[1,1’-biphenyl]-3-carboxylic acid (Y-9)

Yield 54%, white solid. ^1^H NMR (500 MHz, DMSO-*d*_6_) *δ* 9.97 (s, 1H), 9.71 (d, *J* = 10.7 Hz, 2H), 8.18 (d, *J* = 2.4 Hz, 1H), 7.82 (d, *J* = 6.9 Hz, 2H), 7.60 (dd, *J* = 8.6, 2.3 Hz, 1H), 7.56 (d, *J* = 8.6 Hz, 1H), 7.49 (d, *J* = 8.1 Hz, 1H), 7.46 (d, *J* = 3.9 Hz, 1H), 7.42 – 7.39 (m, 2H), 7.34 (d, *J* = 3.9 Hz, 1H), 7.32 – 7.28 (m, 2H), 7.25 – 7.21 (m, 2H), 7.19 (dd, *J* = 7.9, 3.9 Hz, 2H), 7.06 – 7.02 (m, 2H), 6.99 – 6.97 (m, 2H), 6.76 (d, *J* = 8.2 Hz, 2H), 3.45 (s, 0H), 2.81 (t, 2H), 2.56 (t, 2H). ^13^C NMR (126 MHz, DMSO-*d*_6_) *δ* 170.5, 170.0, 169.5, 158.0, 158.0, 148.0, 147.9, 142.4, 140.0, 139.8, 133.6, 133.5, 133.0, 132.1, 131.5, 131.4, 130.7, 130.5, 130.4, 130.2, 12 9.3, 128.9, 123.3, 123.3, 121.2, 121.1, 120.7, 118.1, 117.6, 117.5, 116.6, 116.6, 116.0, 116.0, 112.3, 40.1, 40.0, 39.9, 39.7, 39.4, 39.2, 39.0, 38.2, 30.2. HRMS calcd for C _43_H_33_N_3_O_11_S_4_ [M-H]-, 894.0925; found, 894.0878.

###### 3’-(3-(3-carboxy-4-((5-(3-methoxyphenyl)thiophene)-2-sulfonamido)phenyl)propanamido)-4-((5-(3-methoxyphenyl)thiophene)-2-sulfonamido)-[1,1’-biphenyl]-3-carboxylic acid (Y-10)

Yield 60%, white solid. ^1^H NMR (500 MHz, DMSO-*d*_6_) *δ* 10.05 (s, 1H), 8.15 (s, 1H), 7.82 (d, *J* = 8.1 Hz, 2H), 7.74 (d, *J* = 8.6 Hz, 1H), 7.64 (d, *J* = 8.6 Hz, 1H), 7.61 (d, *J* = 3.9 Hz, 1H), 7.58 – 7.53 (m, 2H), 7.51 (d, *J* = 8.5 Hz, 2H), 7.46 (d, *J* = 3.9 Hz, 2H), 7.31 (td, *J* = 7.7, 4.8 Hz, 3H), 7.26 (d, *J* = 7.8 Hz, 1H), 7.18 (d d, *J* = 8.4, 6.1 Hz, 4H), 6.94 (dd, *J* = 8.9, 2.2 Hz, 2H), 3.78 (d, *J* = 2.1 Hz, 6H), 2.87 (t, *J* = 7.6 Hz, 2H), 2.59 (t, *J* = 7.7 Hz, 2H). ^13^C NMR (126 MHz, DMSO-*D*_6_) *δ* 170.6, 170.1, 169.7, 160.7, 160.0, 158.8, 139.9, 139.9, 133.8, 133.6, 133.3, 132.5, 132.3, 132.2, 131.7, 131.6, 130.6, 130.0, 129.0, 128.9, 128.7, 126.5, 124.2, 124.1, 119. 2, 118.4, 118.4, 115.9, 115.0, 114.9, 111.3, 111.2, 108.0, 104.8, 101.3, 55.4, 55.4, 38. 2, 30.2. HRMS calcd for C_45_H_37_N_3_O_11_S_4_ [M-H]-, 922.1238; found, 922.1200.

###### 3’-(3-(3-carboxy-4-((5-(3,4-difluorophenyl)thiophene)-2-sulfonamido)phenyl)propanamido)-4-((5-(3,4-difluorophenyl)thiophene)-2-sulfonamido)-[1,1’-biphenyl]-3-carboxylic acid (Y-11)

Yield 52%, white solid. ^1^H NMR (500 MHz, DMSO-*d*_6_) *δ* 9.97 (s, 1H), 8.16 (d, *J* = 2.4 Hz, 1H), 7.82 (d, *J* = 8.4 Hz, 3H), 7.78 (dd, *J* = 9.3, 2.2 Hz, 1H), 7.62 (dd, *J* = 8.6, 2.4 Hz, 1H), 7.56 (d, *J* = 8.6 Hz, 1H), 7.51 (t, *J* = 4.2 Hz, 2H), 7.4 7 (t, *J* = 3.5 Hz, 4H), 7.46 – 7.44 (m, 2H), 7.44 – 7.42 (m, 2H), 7.32 – 7.28 (m, 2 H), 7.24 (d, *J* = 8.0 Hz, 1H), 2.83 (t, 2H), 2.57 (t, *J* = 7.7 Hz, 2H). ^13^C NMR (126 MHz, DMSO-*d*_6_) *δ* 170.5, 169.8, 169.4, 150.7, 150.5, 148.8, 148.6, 145.9, 145.4, 14 4.2, 143.4, 139.8, 133.9, 132.8, 132.1, 131.7, 131.6, 130.7, 130.5, 130.1, 130.0, 129.3, 128.8, 124.7, 122.9, 120.7, 120.6, 120.0, 118.5, 118.3, 118.1, 117.6, 116.6, 115.2, 11 5.0, 38.1, 30.1. HRMS calcd for C_43_H_29_F_4_N_3_O_9_S_4_ [M-H]^-^, 934.0650; found, 934.0408. HPLC purity: 99.03% (cf. **Figure S6**).

###### 3’-(3-(3-carboxy-4-((5-(3,4-dichlorophenyl)thiophene)-2-sulfonamido)phenyl)propanamido)-4-((5-(3,4-dichlorophenyl)thiophene)-2-sulfonamido)-[1,1’-biphenyl]-3-carboxylic acid (Y-12)

Yield 55%, white solid. ^1^H NMR (500 MHz, DMSO-*d*_6_) *δ* 9.96 (s, 1H), 8.16 (d, *J* = 2.4 Hz, 1H), 7.96 – 7.85 (m, 2H), 7.80 (d, *J* = 2.3 Hz, 2H), 7.74 – 7.66 (m, 1H), 7.63 (s, 1H), 7.62 – 7.61 (m, 1H), 7.60 – 7.57 (m, 2H), 7.55 (dd, *J* = 6.4, 2.4 Hz, 2H), 7.51 – 7.48 (m, 3H), 7.44 (d, *J* = 3.8 Hz, 1H), 7.40 (d, *J* = 8.4 Hz, 1H), 7.29 (t, *J* = 7.9 Hz, 1H), 7.25 – 7.21 (m, 2H), 2.81 (t, *J* = 7.7 Hz, 2H), 2.56 (t, 2H). ^13^C NMR (126 MHz, DMSO-*d*_6_) *δ* 170.5, 169.7, 169.3, 144.4, 139.9, 139.8, 133.1, 133.0, 132.4, 132.1, 132.0, 131.3, 131.1, 131.0, 130.6, 130.4, 129.3, 128.8, 12 7.3, 125.9, 125.2, 120.7, 120.6, 118.2, 117.9, 117.5, 116.6, 38.1, 30.1. HRMS calcd f or C_43_H_29_Cl_4_N_3_O_9_S_4_ [M-H]^-^, 997.9468; found, 997.9420.

###### 3’-(3-(3-carboxy-4-((5-(3,4-dimethylphenyl)thiophene)-2-sulfonamido)phenyl)propanamido)-4-((5-(3,4-dimethylphenyl)thiophene)-2-sulfonamido)-[1,1’-biphenyl]-3-carboxylic acid (Y-13)

Yield 59%, white solid. ^1^H NMR (500 MHz, DMSO-*d*_6_) *δ* 9.96 (s, 1H), 8.17 (d, *J* = 2.3 Hz, 1H), 7.81 (d, *J* = 2.3 Hz, 2H), 7.61 (dd, *J* = 8.6, 2.3 Hz, 1H), 7.56 (d, *J* = 8.6 Hz, 1H), 7.50 – 7.47 (m, 2H), 7.44 – 7.40 (m, 3H), 7.38 (d, *J* = 2.0 Hz, 1H), 7.34 (dd, *J* = 7.2, 2.9 Hz, 2H), 7.32 – 7.26 (m, 4H), 7.25 – 7.22 (m, 1H), 7.15 (t, *J* = 8.0 Hz, 2H), 2.82 (t, *J* = 7.6 Hz, 2H), 2.56 (t, *J* = 7.7 Hz, 2H), 2.23 – 2.19 (m, 12H). ^13^C NMR (126 MHz, DMSO-*d*_6_) *δ* 170.5, 170.1, 169.6, 148.9, 148. 5, 139.9, 139.8, 137.5, 137.4, 137.3, 137.3, 132.5, 132.2, 131.8, 130.8, 130.3, 130.3, 130.0, 129.8, 129.3, 128.9, 126.7, 123.2, 122.8, 122.8, 121.0, 120.8, 120.6, 118.1, 117. 8, 117.6, 116.7, 38.1, 30.1, 19.3, 19.3, 19.2, 19.2. HRMS calcd for C_47_H_41_N_3_O_9_S_4_ [M-H]^-^, 918.1653; found, 918.1421.

###### 3’-(3-(3-carboxy-4-((5-(2,4-difluorophenyl)thiophene)-2-sulfonamido)phenyl)propanamido)-4-((5-(2,4-difluorophenyl)thiophene)-2-sulfonamido)-[1,1’-biphenyl]-3-carboxylic acid (Y-14)

Yield 52%, white solid. ^1^H NMR (500 MHz, DMSO-*d*_6_) *δ* 10.16 (s, 1H), 8.14 (d, *J* = 2.3 Hz, 1H), 7.88 – 7.81 (m, 4H), 7.71 (dd, *J* = 8.6, 2.3 Hz, 1H), 7.62 (d, *J* = 8.7 Hz, 2H), 7.58 – 7.54 (m, 2H), 7.52 – 7.48 (m, 2H), 7.46 (d, *J* = 4.0 Hz, 1H), 7.45 – 7.38 (m, 3H), 7.31 (t, *J* = 7.8 Hz, 1H), 7.25 (d, *J* = 7.7 Hz, 1H), 7.19 – 7.14 (m, 2H), 2.86 (t, *J* = 7.6 Hz, 2H), 2.60 (t, *J* = 7.7 Hz, 2H). ^13^C NMR (126 MHz, DMSO-*d*_6_) *δ* 170.4, 169.8, 169.4, 163.2, 161.2, 161.1, 159.5, 159.4, 157.5, 15 7.4, 139.8, 139.6, 133.3, 132.2, 131.9, 131.2, 130.9, 130.7, 130.3, 130.2, 129.3, 128.8, 126.2, 120.8, 120.3, 119.4, 118.5, 117.7, 116.9, 116.6, 112.8, 112.6, 105.2, 105.0, 10 4.8, 37.9, 30.0. HRMS calcd for C_43_H_29_F_4_N_3_O_9_S_4_ [M-H]^-^, 934.0650; found, 934.0686.

###### 3’-(3-(3-carboxy-4-((5-(3,5-difluorophenyl)thiophene)-2-sulfonamido)phenyl)propanamido)-4-((5-(3,5-difluorophenyl)thiophene)-2-sulfonamido)-[1,1’-biphenyl]-3-carboxylic acid (Y-15)

Yield 60%, white solid. ^1^H NMR (500 MHz, DMSO-*d*_6_) *δ* 10.10 (s, 1H), 8.60 (s, 1H), 8.15 (d, *J* = 5.2 Hz, 1H), 7.82 (s, 2H), 7.71 (dd, *J* = 8.6, 2.3 Hz, 1H), 7. 64 – 7.57 (m, 5H), 7.55 (d, *J* = 7.9 Hz, 1H), 7.50 – 7.46 (m, 2H), 7.46 – 7.41 (m, 5H), 7.31 (t, *J* = 7.8 Hz, 1H), 7.27 – 7.22 (m, 3H), 2.86 (t, *J* = 7.6 Hz, 2H), 2.59 (t, *J* = 7.6 Hz, 2H). ^13^C NMR (126 MHz, DMSO-*d*_6_) *δ* 170.4, 163.8, 163.7, 163.0, 1 61.9, 161.8, 149.2, 146.6, 145.8, 142.5, 140.7, 139.8, 139.3, 139.0, 136.8, 135.6, 135. 6, 135.5, 135.3, 135.2, 133.9, 133.3, 132.8, 132.4, 131.2, 130.9, 129.3, 128.8, 126.0, 125.9, 124.2, 120.8, 119.5, 118.8, 118.8, 118.5, 117.9, 116.6, 109.4, 109.3, 109.3, 109. 2, 109.2, 109.1, 104.3, 104.1, 37.8, 29.9. HRMS calcd for C_43_H_29_F_4_N_3_O_9_S_4_ [M-H]^-^, 93 4.0650; found, 934.0652.

###### 3’-(3-(3-carboxy-4-((5-(3,4,5-trifluorophenyl)thiophene)-2-sulfonamido)phenyl)propanamido)-4-((5-(3,4,5-trifluorophenyl)thiophene)-2-sulfonamido)-[1,1’-biphenyl]-3-carboxylic a cid (Y-16)

Yield 60%, white solid. ^1^H NMR (500 MHz, DMSO-*d*_6_) *δ* 9.97 (s, 1H), 8. 15 (d, *J* = 2.4 Hz, 1H), 7.82 – 7.79 (m, 2H), 7.71 – 7.65 (m, 4H), 7.60 (dd, *J* = 8. 7, 2.4 Hz, 1H), 7.55 – 7.53 (m, 2H), 7.52 – 7.48 (m, 3H), 7.47 (d, *J* = 3.9 Hz, 1 H), 7.40 (d, *J* = 8.4 Hz, 1H), 7.30 (t, *J* = 7.9 Hz, 1H), 7.27 – 7.22 (m, 2H), 2.82 (t, *J* = 7.7 Hz, 2H), 2.58 – 2.54 (m, 2H). ^13^C NMR (126 MHz, DMSO-*d*_6_) *δ* 170.5, 169.6, 169.2, 151.7, 151.6, 151.6, 149.7, 149.7, 149.6, 149.6, 144.8, 144.4, 144.0, 13 9.9, 139.8, 137.7, 132.7, 131.7, 131.4, 131.3, 130.6, 130.5, 129.3, 128.7, 125.6, 120.7, 120.5, 120.0, 118.3, 118.0, 117.6, 116.5, 110.8, 110.8, 110.8, 110.8, 110.7, 110.6, 38. 1, 30.1. HRMS calcd for C_43_H_27_F_6_N_3_O_9_S_4_ [M-H]^-^, 970.0462; found, 970.0418.

###### 3’-(3-(3-carboxy-4-((5-(2,3,4-trifluorophenyl)thiophene)-2-sulfonamido)phenyl)propanamido)-4-((5-(2,3,4-trifluorophenyl)thiophene)-2-sulfonamido)-[1,1’-biphenyl]-3-carboxylic a cid (Y-17)

Yield 55%, white solid. ^1^H NMR (500 MHz, DMSO-*d*_6_) *δ* 10.17 (s, 1H), 8.13 (d, *J* = 2.4 Hz, 1H), 7.85 – 7.81 (m, 2H), 7.73 – 7.68 (m, 5H), 7.61 – 7.57 (m, 4H), 7.56 – 7.53 (m, 2H), 7.47 (d, *J* = 8.4 Hz, 1H), 7.42 (dd, *J* = 8.5, 2.1 Hz, 1H), 7.31 (t, *J* = 7.9 Hz, 1H), 7.24 (d, *J* = 7.7 Hz, 1H), 2.86 (t, *J* = 7.6 Hz, 2H), 2.60 (t, *J* = 7.7 Hz, 2H). ^13^C NMR (126 MHz, DMSO-*d*_6_) *δ* 170.4, 163.0, 151.6, 15 1.6, 149.7, 149.7, 149.6, 145.9, 145.1, 139.9, 139.3, 137.9, 135.6, 133.9, 133.3, 132.8, 132.5, 131.2, 130.9, 129.3, 129.1, 128.8, 126.0, 125.9, 120.8, 119.5, 118.8, 118.5, 11 7.9, 116.7, 111.1, 111.0, 111.0, 110.9, 110.9, 110.8, 38.2, 29.9. HRMS calcd for C_43_H _27_F_6_N_3_O_9_S_4_ [M-H]^-^, 970.0462; found, 970.0207.

### 4.3. Biological evaluation

#### 4.3.1. Broth microdilution assay

The broth microdilution assay recommended by the Clinical and Laboratory Standards Institute was followed to determine MIC. ^33^ The compound was dissolved in DMSO to make a stock solution. The bacterial inoculum of *A. baumannii* (ATCC 19606) was incubated overnight at 37 °C and transferred to new Cation Adjusted Mueller Hinton broth (CAMHB) until bacterial growth reached the logarithmic phase. The bacteria were then diluted to 10^5^ ∼10^6^ colony forming units/mL in CAMHB. Then, 200 μL of the bacteria suspension was added to the first well of the 96-well plate, while 100 μL of suspension was added to the other 11 wells. The stock solution was added to the first well, and 2-fold serial dilution was performed. Accordingly, compound solutions at concentrations ranging from 128 to 0.06 μg/mL were made. The 96-well plate was incubated at 37 °C for 18∼24 h. Lastly, the concentration at which bacterial growth was completely inhibited was determined as the MIC value of the compound. MIC measurements were done in triplicate. Meropenem was used as the positive control.

#### 4.3.2. Construction of AbMsbA and ATPase activity assay

##### 4.3.2.1. Overexpression and purification of AbMsbA

AbMsbA with an N-terminal His-tag was cloned into the pET-28a vector and expressed in *E. coli* C43 (DE3) cells. Cultures were grown in Terrific Broth at 37°C until OD_600_ reached about 1.0, induced with 1 mM IPTG, and incubated overnight at 18°C. Cells were harvested, lysed by high-pressure homogenization in lysis buffer (50 mM Tris-HCl pH 8.0, 300 mM NaCl, 10% glycerol, 1 mM TCEP), and subjected to low-speed centrifugation to remove debris. Membranes were isolated by ultracentrifugation (150,000 × g, 1 h, 4°C), resuspended, and solubilized in 1% DDM with gentle stirring at 4°C. The solubilized material was loaded onto TALON cobalt affinity resin, washed extensively with Buffer A (including 0.1% DDM and 10 mM imidazole), and eluted with Buffer B containing 250 mM imidazole. The eluate was concentrated using a 100 kDa MWCO device and further purified by size-exclusion chromatography on a Superdex 200 column equilibrated with SEC buffer (25 mM Tris-HCl pH 8.0, 150 mM NaCl, 5% glycerol, 0.5 mM TCEP, 0.05% DDM). Peak fractions were pooled and concentrated to 10 mg/mL for subsequent nanodisc reconstitution.

##### 4.3.2.2. Nanodisc reconstitution

Purified MsbA was reconstituted into nanodiscs by incubating with membrane scaffold protein MSP1D1 and POPG lipids at a molar ratio of 1:2:120 in nanodisc buffer (25 mM Tris-HCl pH 8.0, 150 mM NaCl) for 1 hour at 4°C. Detergent removal was performed by adding 0.6 g/mL Bio-Beads SM-2, followed by overnight incubation at 4°C. After bead removal, the mixture was further purified by size-exclusion chromatography using the same buffer. The resulting MsbA-containing nanodiscs were concentrated and used for ATPase activity measurements.

##### 4.3.2.3. ATPase activity measurement

ATP hydrolysis by AbMsbA was quantified using a molybdate–ascorbic acid-based colorimetric assay. Purified protein was pre-incubated with test compounds (0, 1 and 10 µM) in a 96-well plate at room temperature for 10 min in a total volume of 5 µL. The reaction was initiated by adding 20 µL of pre-warmed reaction buffer (20 mM HEPES pH 7.5, 100 mM NaCl, 2 mM ATP, 4 mM MgCl₂, 2 mM DTT) and incubated at 37°C for 20 min. The reaction was terminated by the addition of 25 µL of 12% (w/v) SDS. Subsequently, 50 µL of freshly prepared Solution A (2% ammonium molybdate and 12% ascorbic acid, mixed 1:1) was added and incubated at room temperature for 5 min. Finally, 75 µL of color development solution (2% sodium citrate, 2% sodium meta-arsenite, 2% acetic acid) was added, followed by a 20-min incubation. Absorbance was measured at 750 nm using a microplate reader.

#### 4.3.3. Time-kill curves

Overnight-cultured *A. baumannii* (ATCC 19606) was diluted with CAMHB medium (5 mL) containing an inoculum of 1 × 10^6^ CFU/mL per strain. Distilled water or compounds were added to yield concentrations of 0 ×, 1 ×, 4 × and 8 ×MIC. At the time point of 0, 2, 4, 8, 18, and 24 h, cultures incubated at 37°C with 200 rpm were serially diluted (10-fold) in sterile saline. 10 μL of each serial dilution was placed onto MH agar plates in duplicate and incubated at 37°C for 16 h. The viable colonies were counted and presented as logarithms of log_10_ (CFU/mL). The same procedure was repeated twice.

#### 4.3.4. Spontaneous resistance assay

*A. baumannii* (ATCC19606) was incubated in CAMHB medium overnight at 37°C. The bacteria were collected by centrifugation at 3,000 rpm, and the supernatant was removed and the concentration of the bacteria was adjusted to 1 × 10^10^ CFU/mL with sterile physiological saline. The bacterial solution (100 µL) was placed onto MH agar plates containing compounds at the concentration of 16 × MIC, incubated at 37°C for 48 h, and the number of colonies on the plates was recorded. The spontaneous resistance frequency is the ratio of the number of colonies on the plate to the actual inoculum CFU of the plate.

#### 4.3.5. TEM analysis

Under 200 rpm at 37℃, *A. baumannii* (ATCC19606, 1×10^7^ CFU/mL) was cultured in CAMHB medium with the compound at the concentration of 0.25 μg/mL, 0.5 μg/mL and 1 μg/mL for 14 h. Cells were pelleted, washed with PBS, fixed with 1.5 mL of fixative (2.5% glutaraldehyde), post-fixed in 1% reduced osmium tetroxide (Ted Pella Inc.) for 2 h. The samples were dehydrated through an ascending series of ethanol (30%, 50%, 70%, 80%, 95%, 100%), then washed with propylene oxide and finally embedded in 812 Kit (SPI-PON). Ultrathin sections (60∼80 nm) were cut with an Ultracut microtome (RMC). The sections were stained with a saturated solution of 2% uranyl acetate in alcohol. Staining was performed with 2.6% lead citrate solution protected by CO_2_. Then, the morphology of outer membrane was observed under HT7800 TEM (HITACHI), and images were recorded.

#### 4.3.6. Hemolysis test

An appropriate amount of decellularized rabbit blood was diluted with saline (1:10, v/v), centrifuged at 1,000∼1,500 rpm for 15 min, and washed for 2∼3 times with saline until the supernatant was clear. A 2% red blood cell suspension was then prepared in saline. For each assay, 600 μL of the red blood cell suspension was mixed with the compound solution (dissolved in DMSO) and diluted with saline to a final volume of 1.2 mL, yielding final concentrations of 0.625∼100 μg/mL (DMSO ≤1%). Samples were incubated at 37°C for 3 h, then centrifuged at 800 g for 5 min. Subsequently, 200 μL of the supernatant was transferred to a 96-well plate. Three parallel portions were prepared, and the absorbance was measured at 540 nm and 720 nm using an enzyme-linked immunosorbent assay reader (A = A_540nm_ -A_720nm_). The hemolysis rate was calculated as [(A_sample − A_negative)/(A_positive − A_negative)] × 100%. 1% Triton X-100 at the same concentration and saline served as the positive and negative control, respectively. The HC_50_ value was calculated using GraphPad Prism 5.

#### 4.3.7. Cytotoxicity assay

The cytotoxicity of compounds to HEK293 was evaluated by using the Cell Counting Kit-8 (CCK-8) assay (Beyotime, China). Briefly, the cells were seeded into 96-well flat-bottom plates containing the culture medium (10% fetal bovine serum) and serial concentrations of compounds, at a density of 1 × 10^4^ cells per well. The plates were incubated at 37°C in a humidified atmosphere containing 5% CO_2_ for 72 h. 10 µL of the CCK-8 reagent was directly added to each well. The plates were incubated for 30 min at 37°C. The optical density (OD) was measured at 450 nm using a ELx800 plate reader (BioTek). Cell viability was calculated and expressed as a percentage, with the untreated cells as reference. CC_50_ was determined using nonlinear regression with normalized dose-response fit implemented in GraphPad Prism 5.

The serum-free cytotoxicity assay was the same except for that the culture medium was FBS free and the time of co-incubation was 12 h.

#### 4.3.8. Animal studies

##### 4.3.8.1. General

Animals were obtained from SPF Biotechnology Co., Ltd. (Beijing, China). All animal experiments were approved by the Animal Care and Use Committee of the Institute of Materia Medica, Chinese Academy of Medical Sciences (Beijing, China). Animal care and experimental procedures were conducted in accordance with the Beijing Administration Rule of Laboratory Animals. Mice were housed at 22 °C and 50% humidity, with a cycle of 12 h light: 12 h dark.

##### 4.3.8.2. In vivo pharmacokinetics studies

Six Kunming mice (male: female= 1:1) of 5∼7 weeks were used for studying pharmacokinetics properties. **Y-11** and **Cpd 4** was respectively dissolved in 0.5% CMC-Na and intravenously injected to mice at 5 mg/kg. 0.2 mL of blood sample was collected via the retro-orbital sinus at 5 min, 15 min, 30 min, 1 h, 2 h, 4 h, 6 h, 8 h, 24 h, respectively. Each sample was treated with K_2_EDTA immediately after collection, and then centrifuged at 8,000 rpm at 8℃ for 6 min. The compound concentration of each sample was quantified by HPLC. Accordingly, the time-concentration curve was plotted, based on which the pharmacokinetic parameters were determined by non-compartmental analysis using Phoenix.

##### 4.3.8.3. Murine infection model

Six-week old ICR mice (22∼25 g) were randomly divided into 5 groups (n=5). Mice were intraperitoneally infected with 0.5 mL of saline containing 2.0 × 10^8^ CFU of *A. baumannii* (ATCC 19606). After 0.5 h, mice were injected intraperitoneally with saline (vehicle), meropenem (50 mg/kg), **Cpd 4** (200 mg/kg), **Y-11** (200 mg/kg), respectively. Mice were sacrificed at 8 h after infection. Spleen was aseptically excised, weighed, homogenized in 1 mL of saline, serially diluted, and plated on Mueller Hinton (MH) agar plates for CFU determination. Each spleen was counted twice, and the bacterial colony counts were represented by log_10_ CFU/g. Spleens were also collected from the infected but untreated mice, which was used for comparison.

## Supporting information

Supplemental NMR

## ASSOCIATED CONTENT

### Supporting Information

The Supporting Information is available free of charge via the Internet at http://pubs.acs.org.

- Detailed results and structural characterization spectra of compounds (Supporting_Information.docx)
- Molecular formula strings and activity data (compounds.csv)

## ABBREVIATIONS USED

LPS: lipopolysaccharide; *A. baumannii: Acinetobacter baumannii*; CRAB: carbapenem-resistant *A. baumanni*; ABC: ATP binding cassette; LOS: lipooligosaccharide; ATP: adenosine triphosphate; ADME/T: absorption, distribution, metabolism, excretion and toxicity.; PPB: plasma protein binding; AI: artificial intelligence; MRSA: methicillin-resistant *Staphylococcus aureus*; SA: synthetic accessibility; TEM: transmission electron microscopy; RL: reinforcement learning; MPO: multi-parameter optimization; DMF: N,N-Dimethylformamide; DCM: dichloromethane; TEA: triethylamin; TFA: trifluoroacetic acid; TCFH: N,N,N’,N’-tetramethylchloroformamidinium hexafluorophosphate; NMI: 1-methylimidazole; MIC: minimal inhibit concentration; NMR: nuclear magnetic resonance; MS: mouse serum; SAR: structure-activity relationship; HEK293: human embryonic kidney 293 Cells; i.p.: intraperitoneal injection; MD: molecular dynamics simulation; DS: Discovery Studio; TLC: thin-layer chromatography; HRMS: high-resolution mass spectrometry; HPLC: high-performance liquid chromatography; EtOAc: ethyl acetate; PE: petroleum ether

## AUTHOR INFORMATION

### Author contributions

L.W., B.J and Q.L. contributed equally to this work. L.W. designed and conducted the AI-based drug design and chemical synthesis. B.J. designed, conducted, and analyzed biological activities of compounds. Q. L conducted virtual screening and molecular dynamics simulations. J. Z. conducted the enzyme activity test. Y. Y. contributed to data collection and helped with computational modelling. H. J. provided technical support and resources of computational modeling. L.W. and J.X. wrote the manuscript. S.W., L.P., Y.L and J.X. supervised the studies. All authors have approved the final version of the manuscript.

### Notes

The authors declare that they have no known competing financial interests.

## ACKNOWLEDGMENTS

This work was supported by the CAMS Innovation Fund for Medical Sciences (Grant No.: 2021-I2M-1-069, awarded to J.X. and S.W.), the Fundamental Research Funds for Central Universities, Chinese Academy of Medical Sciences & Peking Union Medical College (Grant No.: 3332026197, awarded to L.W.) and the Guangdong Innovative and Entrepreneurial Research Team Program (Grant No.: 2023ZT10Y013, awarded to L.P.). We thank the assistance of Southern University of Science and Technology (SUSTech) Cryo-EM Center. Y.L. is an investigator of SUSTech Institute for Biological Electron Microscopy. Y.L. discloses support for the research of this work from the National Key R&D Program of China (Grant No.:2024YFA0919900), National Natural Science Foundation of China (Grant No.: 32322004, U22A20338), and the Guangdong Innovative and Entrepreneurial Research Team Program (Grant No.: 2023ZT10Y013).

