## Supplemental NMR for "Deep reinforcement learning-driven discovery of a MsbA-targeted small-molecule antibiotic for the treatment of *Acinetobacter baumannii* infection"

[1] *State Key Laboratory of Bioactive Substance and Function of Natural Medicines, Institute of Materia Medica, Chinese Academy of Medical Sciences and Peking Union Medical College, Beijing 100050, China;* [2] *Department of Clinical Laboratory, The Fifth Affiliated Hospital of Guangzhou Medical University, Guangzhou 510700, China;* [3] *Institute for Biological Electron Microscopy, Southern University of Science and Technology, Shenzhen, Guangdong, 518055, China;* [4] *Department of Chemical Biology, School of Life Sciences, Southern University of Science and Technology, Shenzhen, Guangdong, 518055, China;* [5] *State Key Laboratory of Natural and Biomimetic Drugs, School of Pharmaceutical Sciences, Peking University, Beijing 100191, China.*

[#] These authors contributed equally to this work.

Jie Xia & Song Wu, Institute of Materia Medica, Chinese Academy of Medical Sciences and Peking Union Medical College

Yanyan Li, Southern University of Science and Technology

Liang Peng, The Fifth Affiliated Hospital of Guangzhou Medical University

### Table of Contents

1. **Table S1.** Five ensemble models for anti-*A. baumannii* activity prediction.........S3

2. **Table S2.** The selected molecules by virtual screening. .......................................S4

3. **Table S3.** Antibacterial activity spectrum of **Cpd 4**, **In-1**, **Y-11**. ..........................S6

4. **Table S4.** *In vivo* pharmacokinetics parameters of **Y-11** ......................................S7

5. **Figure S1.** Cerastecins with diverse linkers .......................................................S8

6. **Figure S2.** Cerastecin C bound to MsbA .............................................................S9

7. **Figure S3.** Reinforcement learning curves of the Finalscore.............................S10

8. **Figure S4.** $^{1}$H-, $^{13}$C-NMR and HRMS spectra for the derivatives ....................S11

9. **Figure S5.** The RMSD plot from molecular dynamics simulation ...................S58

10. **Figure S6.** HPLC purity data of **Y-11** .............................................................S59

**Table S1.** Five ensemble models for anti-*A. baumannii* activity prediction, built with AutoMolDesigner.

| Model | fingerprints | TN | FP | FN | TP | Accuracy | AUROC | MCC | F1 score |
|---|---|---|---|---|---|---|---|---|---|
| M1 | Rdkit_2D_normal | 96 | 5 | 4 | 75 | 0.950 | 0.983 | 0.899 | 0.943 |
| M2 | Rdkit_2D | 96 | 5 | 4 | 75 | 0.950 | 0.982 | 0.899 | 0.943 |
| **M3** | **ECFP-4** | **97** | **4** | **4** | **75** | **0.956** | **0.979** | **0.910** | **0.949** |
| M4 | FCFP-6 | 96 | 5 | 4 | 75 | 0.950 | 0.971 | 0.899 | 0.943 |
| M5 | MACCS | 96 | 5 | 4 | 75 | 0.950 | 0.983 | 0.899 | 0.943 |

**Table S2.** The selected molecules, including their linkers, Chemgauss4 scores(FRED), probability of being active (AutoMolDesigner), and synthesis accessibility (SA).

| Compound ID | linker | FRED Chemgauss4 Score | Probability of being active against *A. baumanni* | SA Score |
|---|---|---|---|---|
| Cpd 4 | 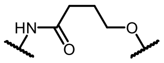 | -12.7748 | -- | -- |
| I-1 | 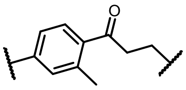 | -17.1104 | 0.52 | 3.29 |
| I-2 | 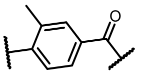 | -15.9363 | 0.54 | 3.18 |
| I-3 | 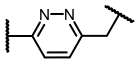 | -15.4808 | 0.52 | 3.25 |
| I-4 (**In-1**) | 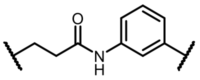 | -15.1477 | 0.56 | 3.23 |
| I-5 | 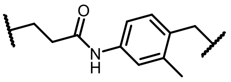 | -15.0982 | 0.64 | 3.29 |
| I-6 | 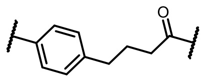 | -14.8665 | 0.54 | 3.22 |
| I-7 | 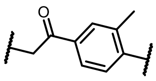 | -14.7365 | 0.51 | 3.22 |
| I-8 | 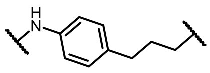 | -14.7329 | 0.57 | 3.17 |
| I-9 | 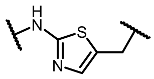 | -14.3693 | 0.60 | 3.26 |
| I-10 | 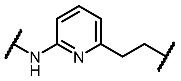 | -14.3505 | 0.50 | 3.28 |
| I-11 | 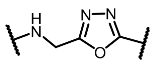 | -14.1810 | 0.61 | 3.24 |
| I-12 | 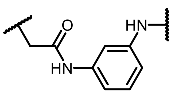 | -14.0513 | 0.62 | 3.24 |

| | | | | |
|---|---|---|---|---|
| I-13 | 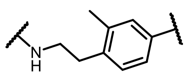 | -13.9949 | 0.58 | 3.26 |
| I-14 | 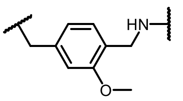 | -13.7020 | 0.60 | 3.29 |
| I-15 | 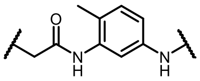 | -13.4388 | 0.62 | 3.28 |
| I-16 | 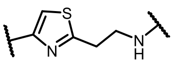 | -13.0069 | 0.51 | 3.27 |
| I-17 | 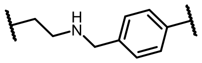 | -12.9087 | 0.55 | 3.17 |
| I-18 | 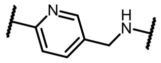 | -12.8635 | 0.58 | 3.20 |
| I-19 | 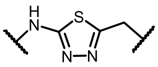 | -12.7836 | 0.52 | 3.26 |

**Table S3.** Antibacterial activity of **Cpd 4**, **In-1** and **Y-11**, with Meropenem as a positive control.

| Strain | Resistant bacteria | MIC (μg/mL) | | | |
|---|---|---|---|---|---|
| | | **Cpd 4** | **In-1** | **Y-11** | Meropenem |
| NCTC 13304 | Yes | 0.5 | 2 | 1 | 32 |
| CRAB24-1 | Yes | 0.5 | 2 | 1 | >32 |
| CRAB24-2 | Yes | 0.5 | 2 | 1 | >32 |
| CRAB24-3 | Yes | 1 | 2 | 1 | 32 |

**Table S4.** *In vivo* pharmacokinetics parameters of **Y-11**.

| Parameter | 5 mg/kg (intravenous injection) | |
|---|---|---|
| | **Y-11** | **Cpd 4** |
| $C_{max}$ (ng/mL) | 44,567 | 52,733 |
| $T_{max}$ (h) | 0.0833 | 0.0833 |
| $t_{1/2}$ (h) | 3.96 | 4.05 |
| $AUC_{0-t}$ (h*ng/m) | 14,582 | 21,878 |
| $AUC_{0-\infty}$ (h*ng/m) | 14,594 | 21,902 |
| $MRT_{0-t}$ (h) | 0.520 | 0.782 |
| $MRT_{0-\infty}$ (h) | 0.543 | 0.822 |

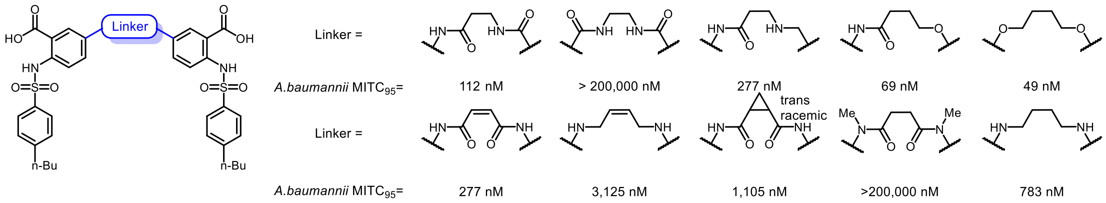


**Figure S1**. Cerastecins with diverse linkers and their anti-*A. baumannii* activity

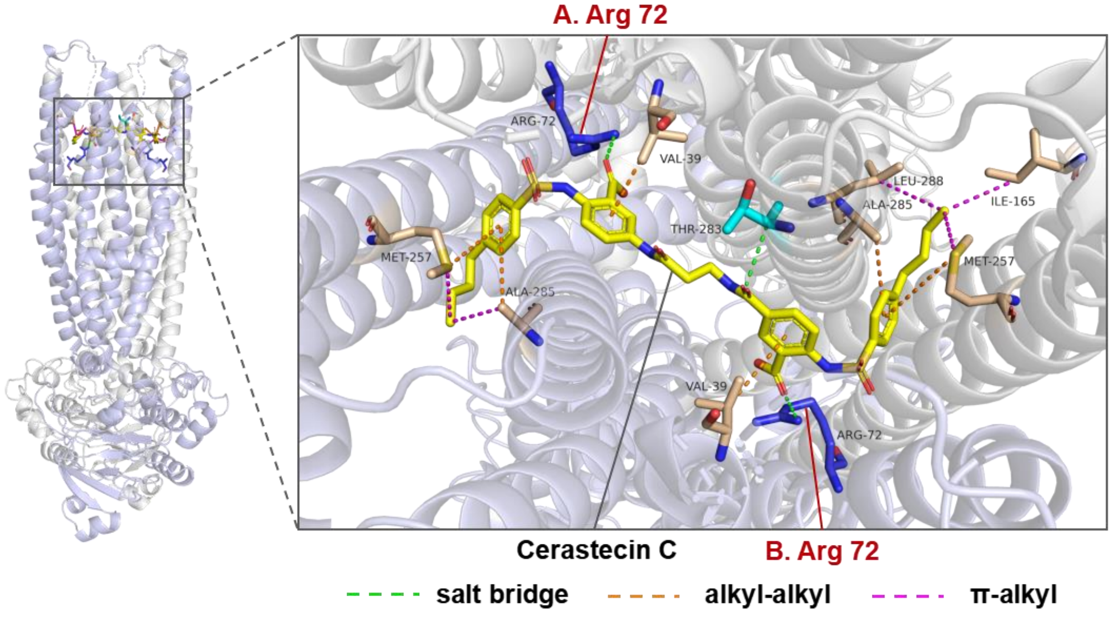


**Figure S2.** Cerastecin C bound to MsbA (PDB Entry: 8GK7). The salt bridge between each carboxyl group of cerastecins and Arg-72 of MsbA is highlighted.

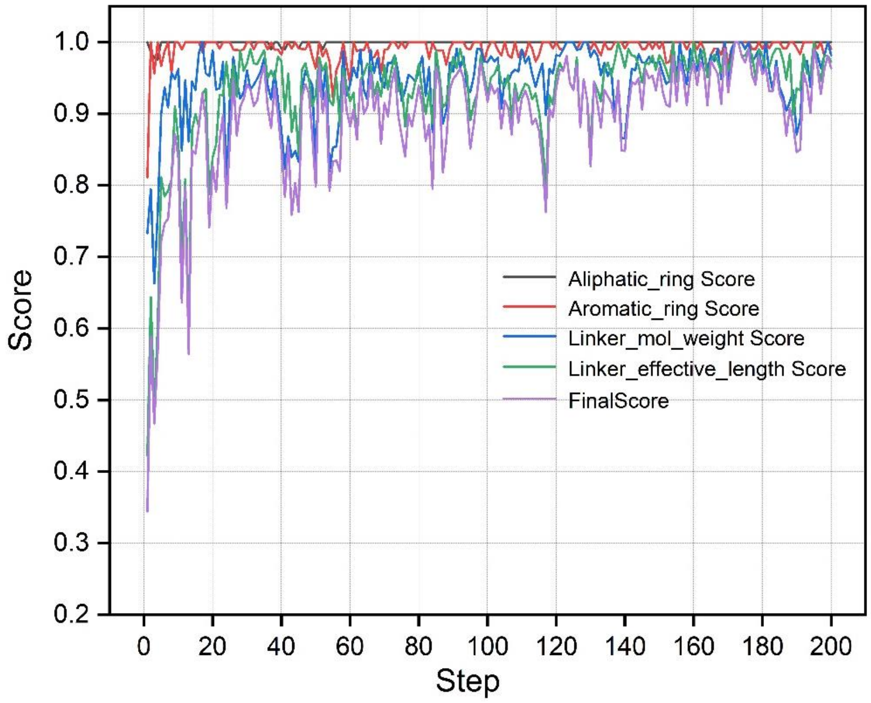


**Figure S3.** Learning curves of the Finalscore and each single component score during 200 Reinforcement Learning steps.

**Figure S4.** $^1$H-NMR, $^{13}$C-NMR and HRMS spectra of the intermediates **C1-C11** or derivatives **Y1-Y17**.

*2-(4-chlorophenyl)thiophene (**C1**).*

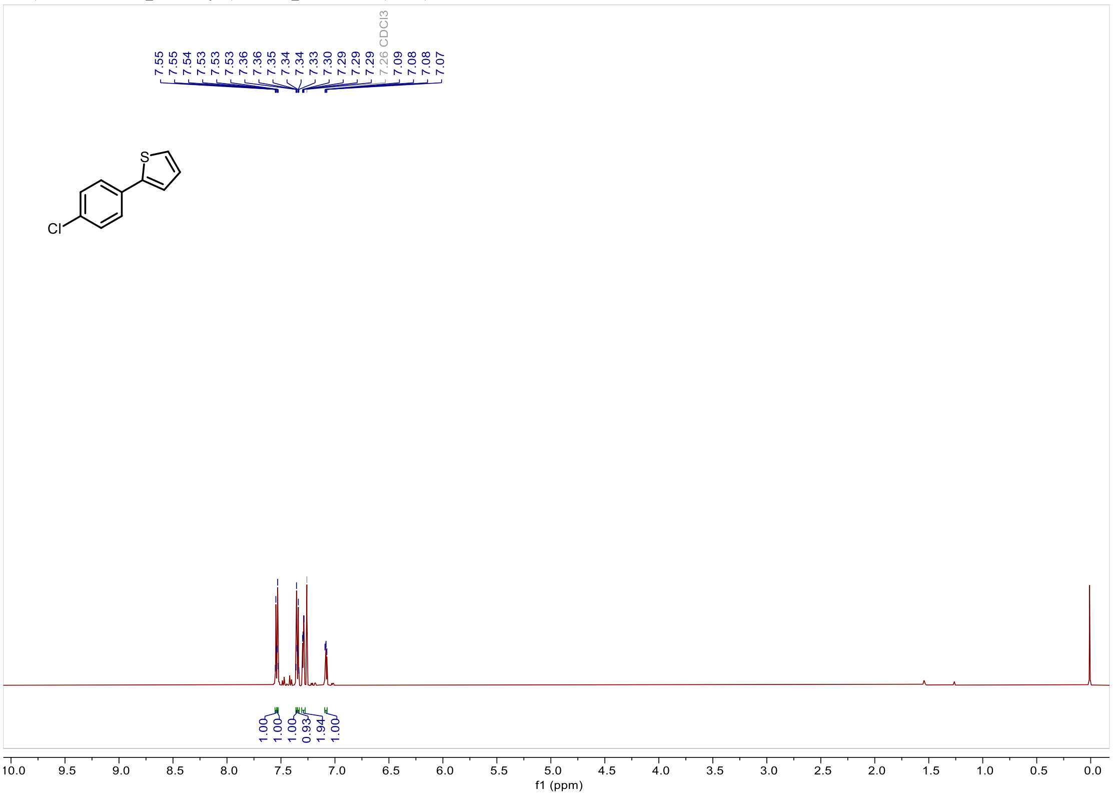

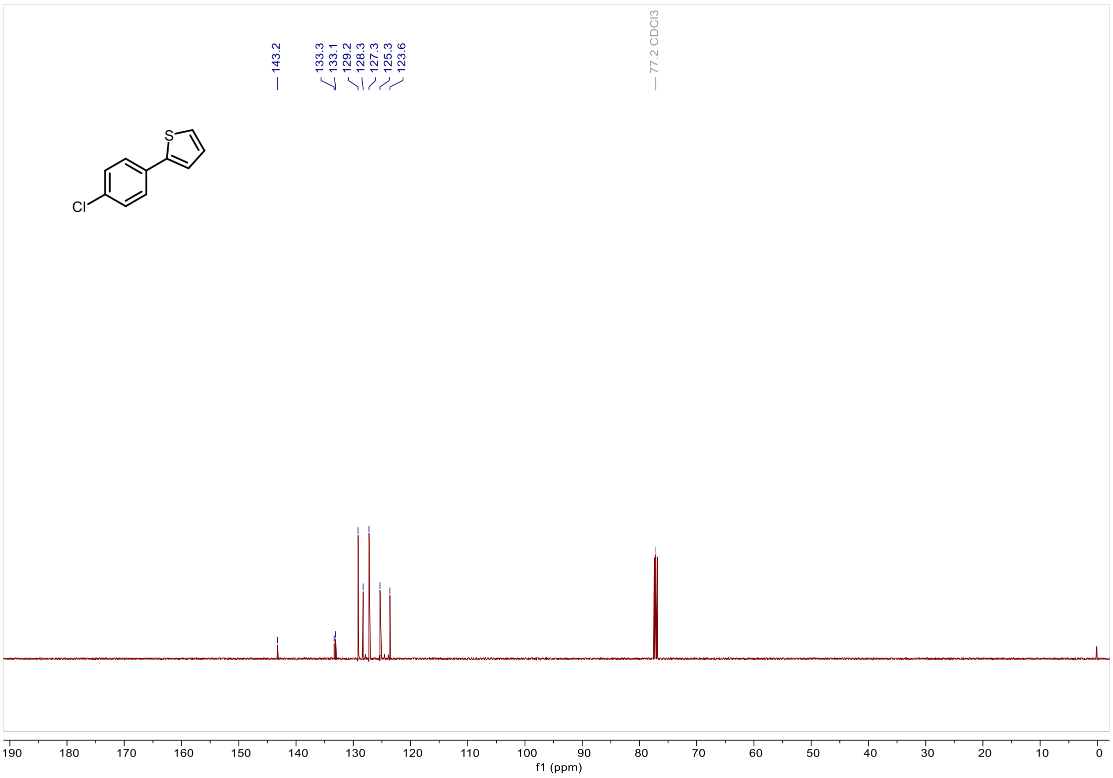

*5-(4-chlorophenyl)thiophene-2-sulfonic acid (**C2**).*

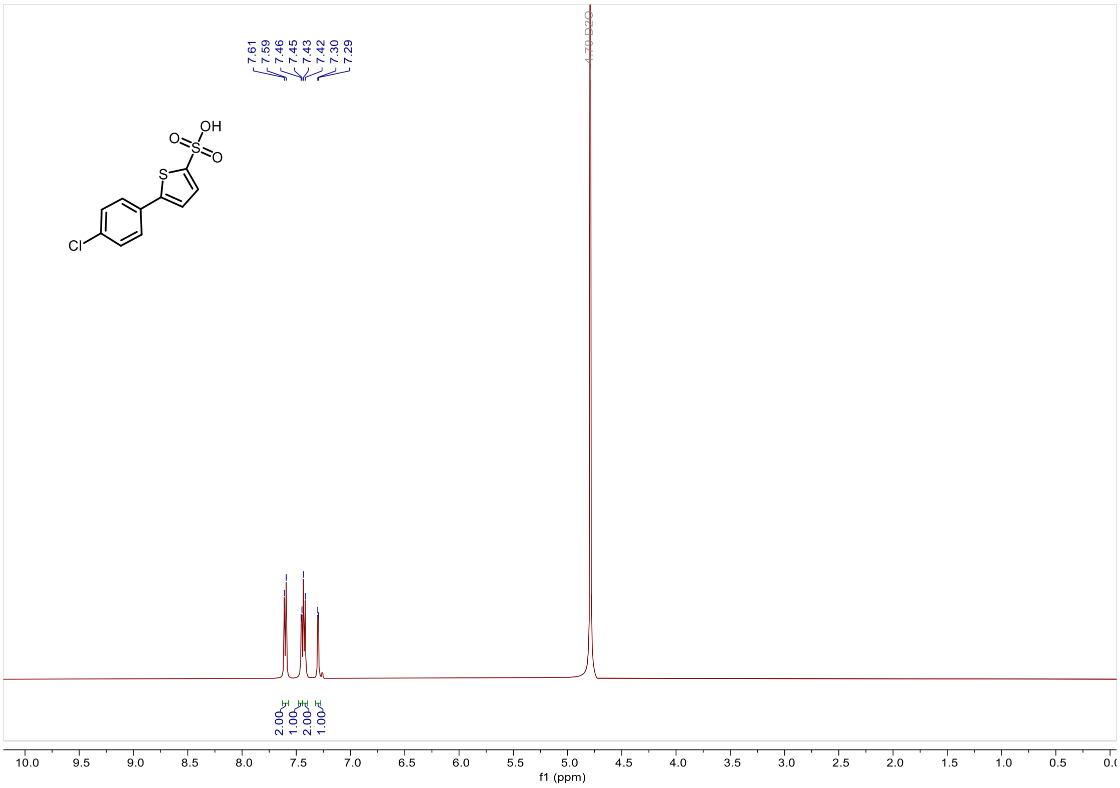


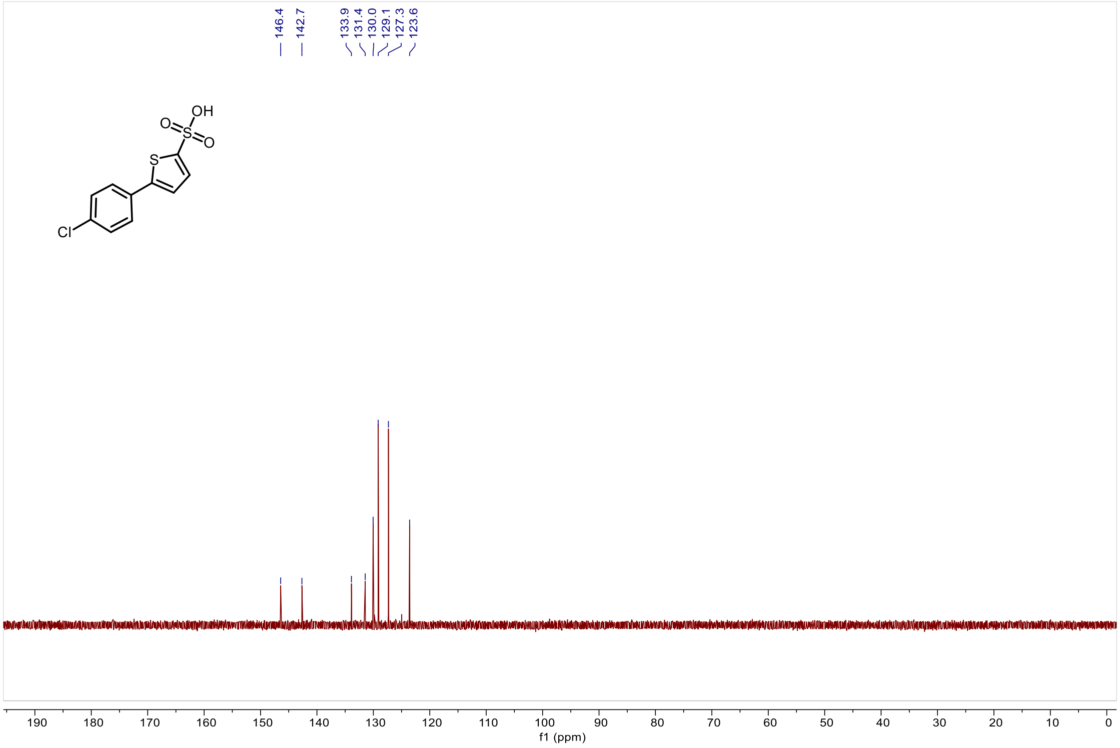

*5-(4-chlorophenyl)thiophene-2-sulfonyl chloride (**C3**).*

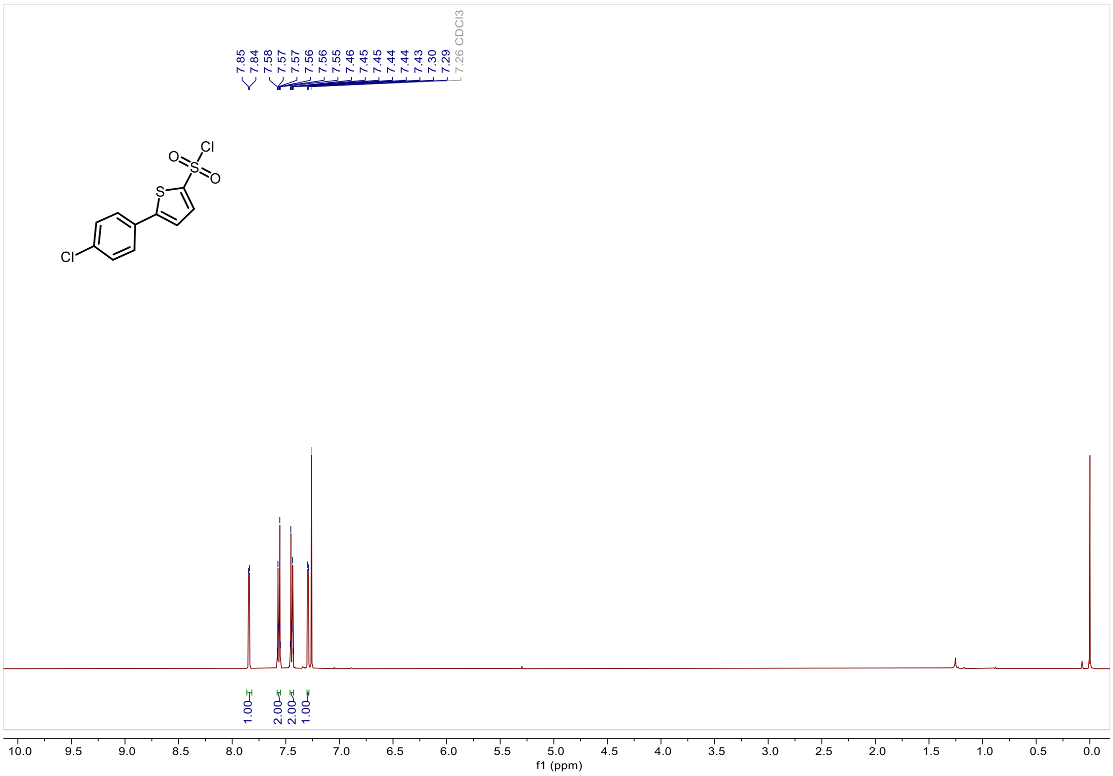


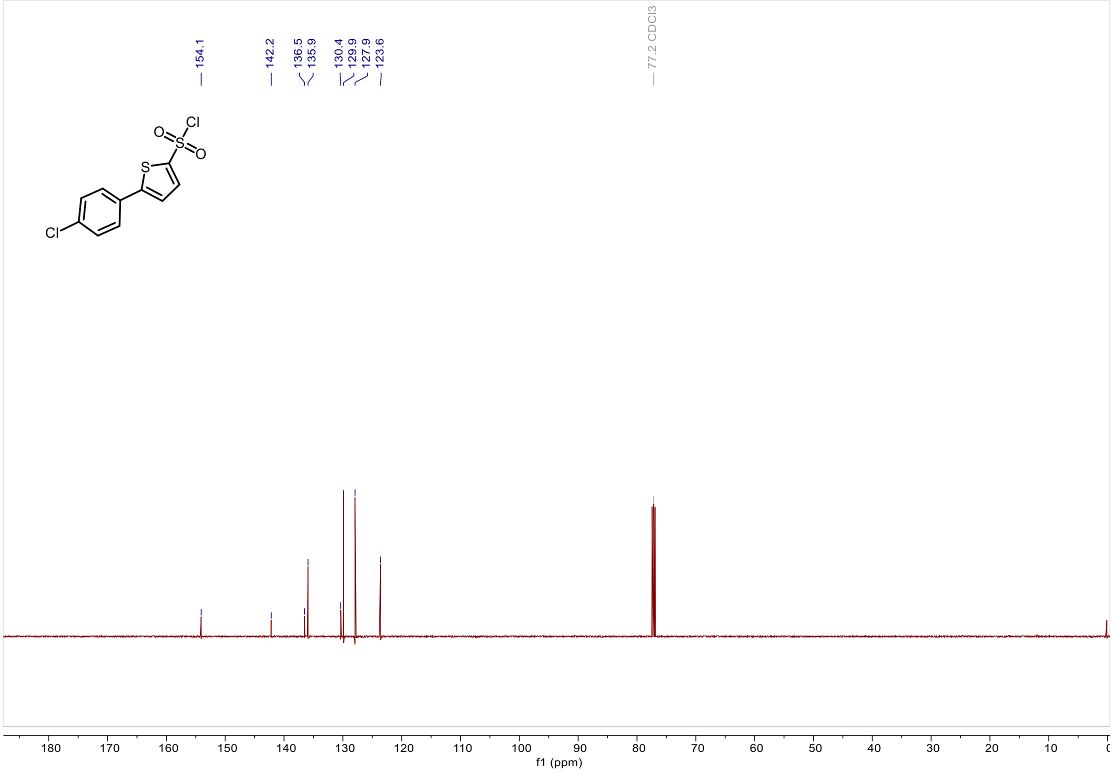

*2-((5-(4-chlorophenyl)thiophene)-2-sulfonamido)-5-iodobenzoate (**C4**).*

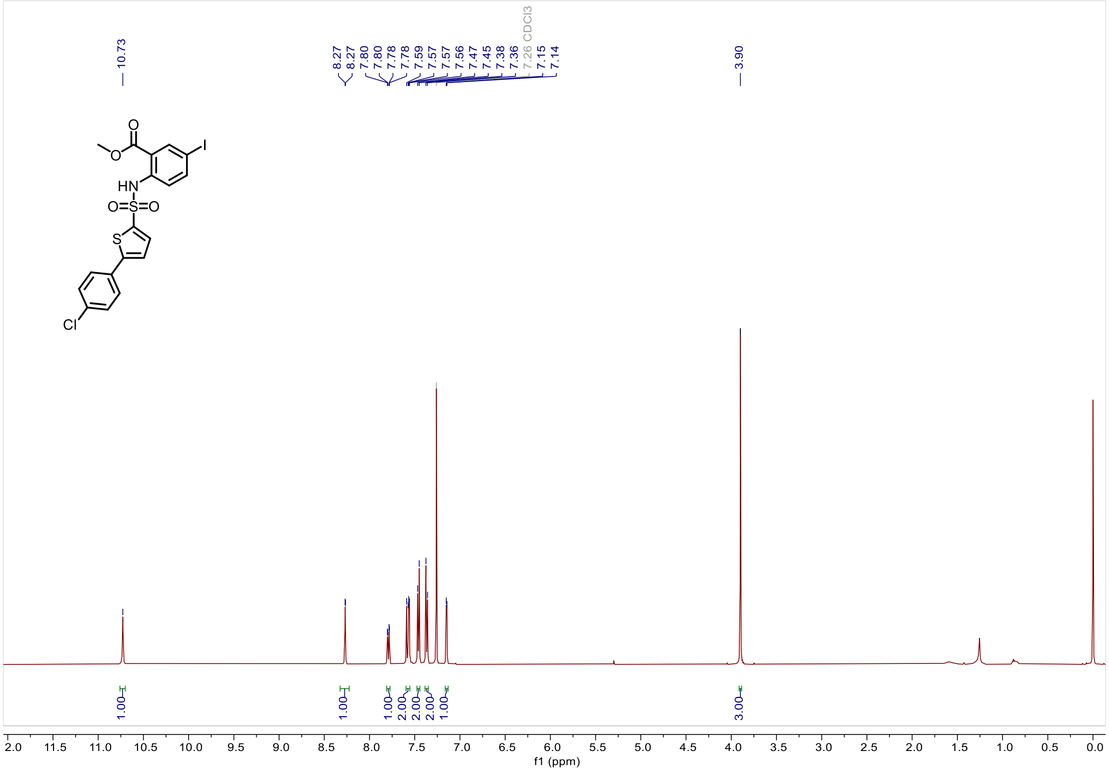


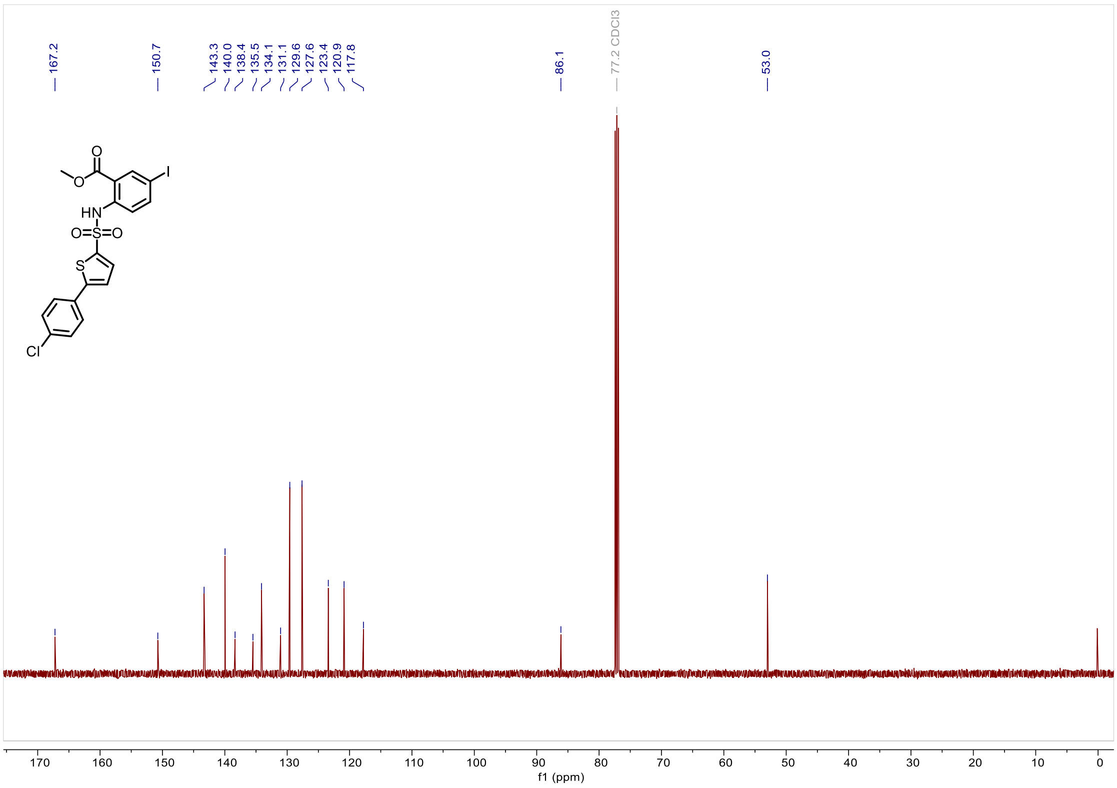

*3-(4-((5-(4-chlorophenyl)thiophene)-2-sulfonamido)-3-(methoxycarbonyl)phenyl)acrylic acid (**C5**).*

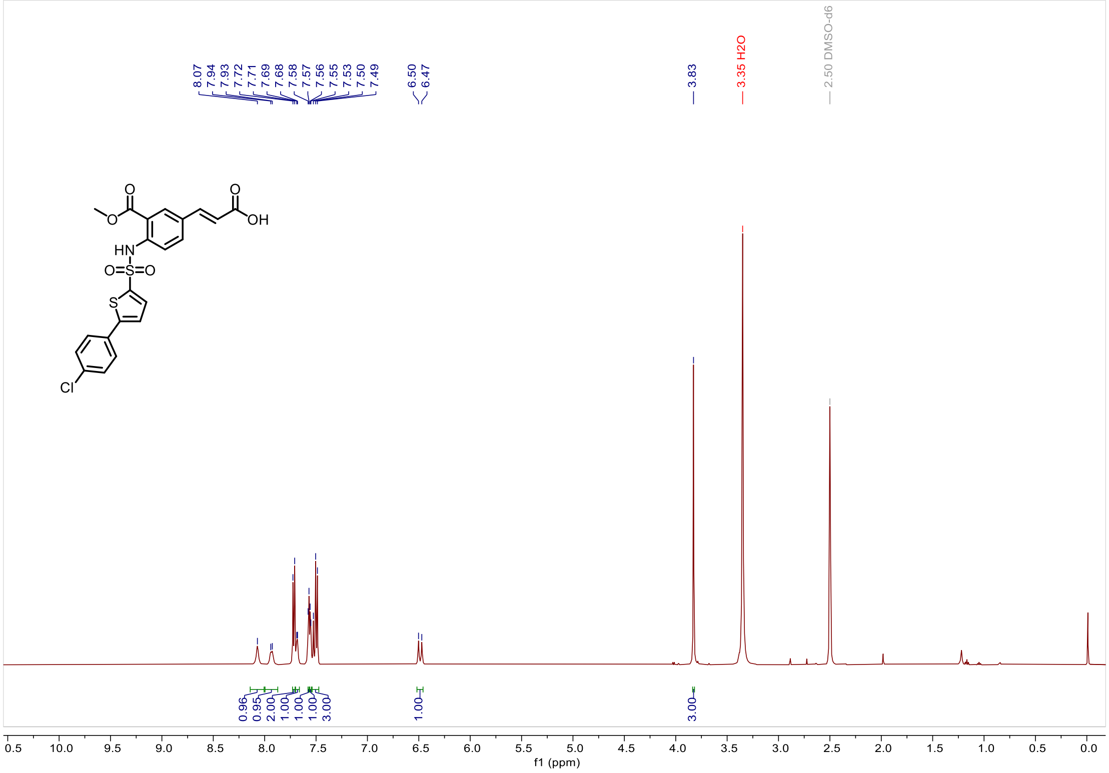


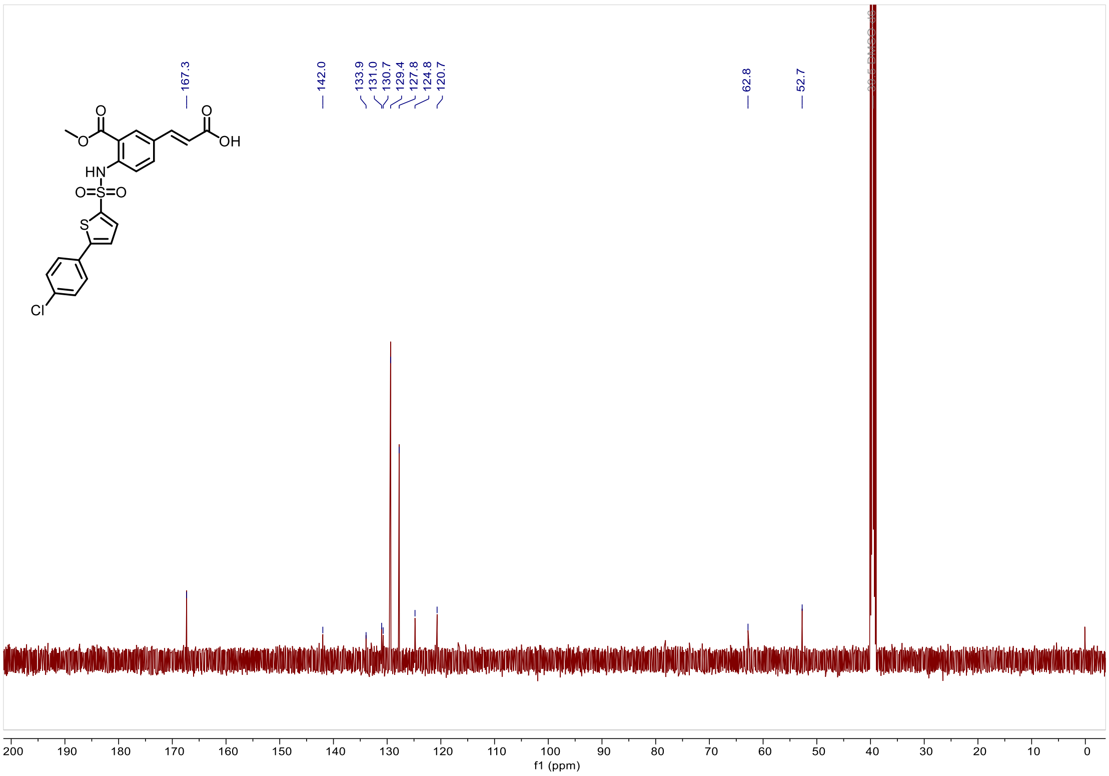

*3-(4-((5-(4-chlorophenyl)thiophene)-2-sulfonamido)-3-(methoxycarbonyl)phenyl)propanoic acid (**C6**).*

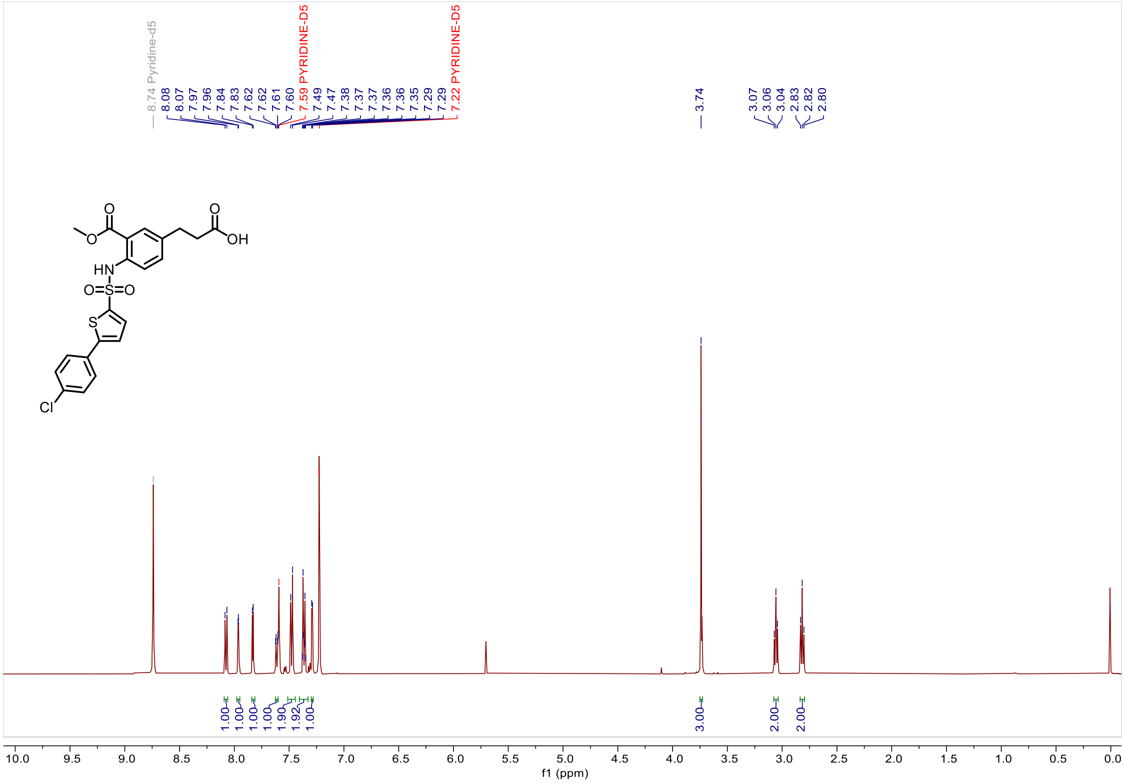

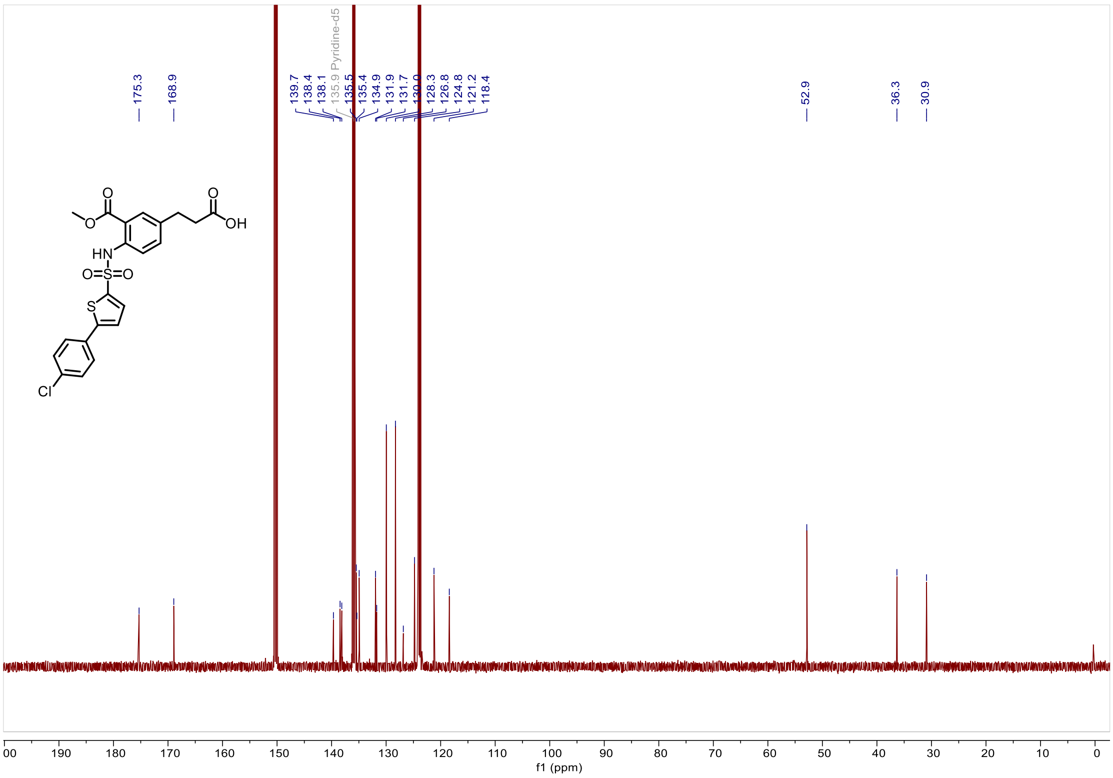

*methyl 3'-amino-4-((5-(4-chlorophenyl)thiophene)-2-sulfonamido)-[1,1'-biphenyl]-3-carboxylate (**C**7).*

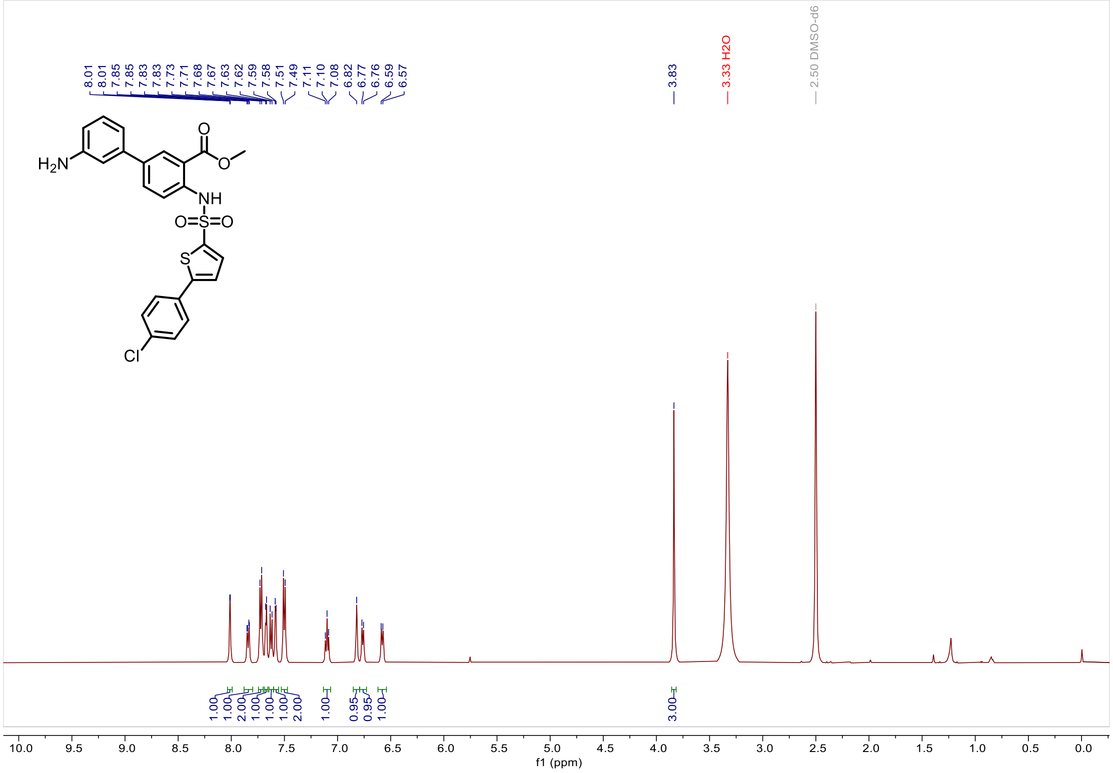


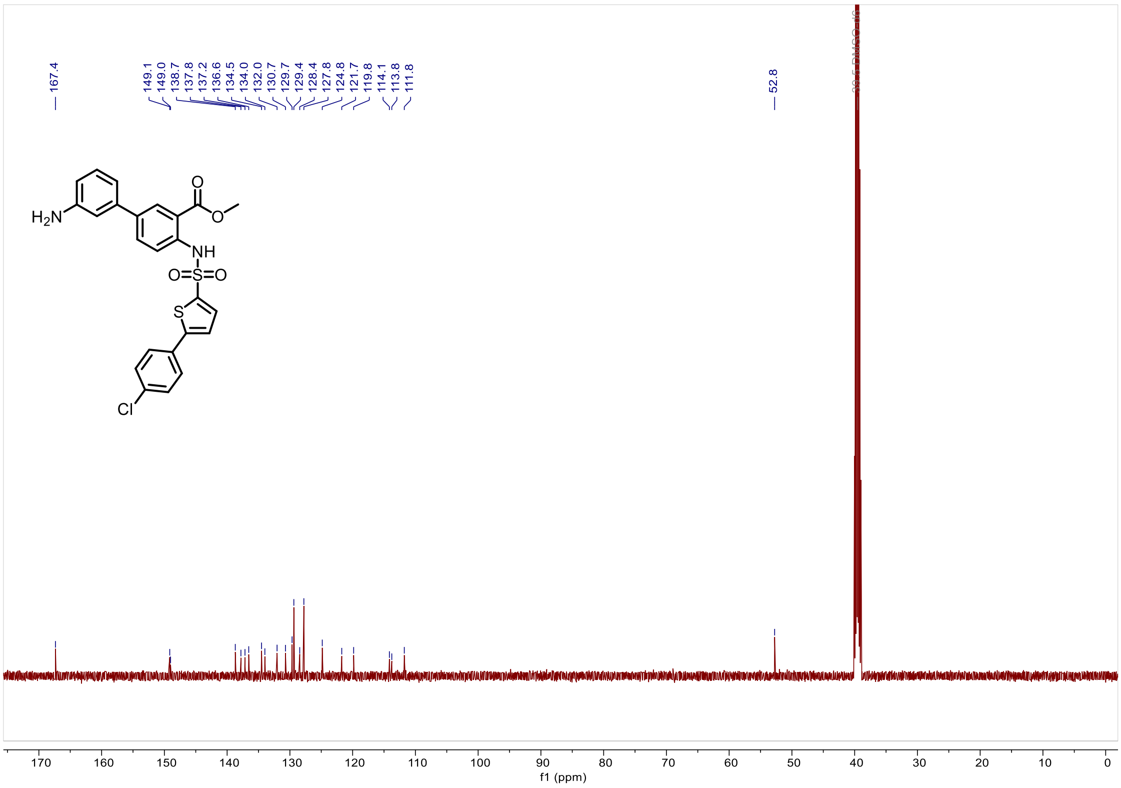

*3'-(3-(3-carboxy-4-((5-(4-chlorophenyl)thiophene)-2-sulfonamido)phenyl)propanamido)-4-((5-(4-chlorophenyl)thiophene)-2-sulfonamido)-[1,1'-biphenyl]-3-carboxylic acid (**In-1**).*

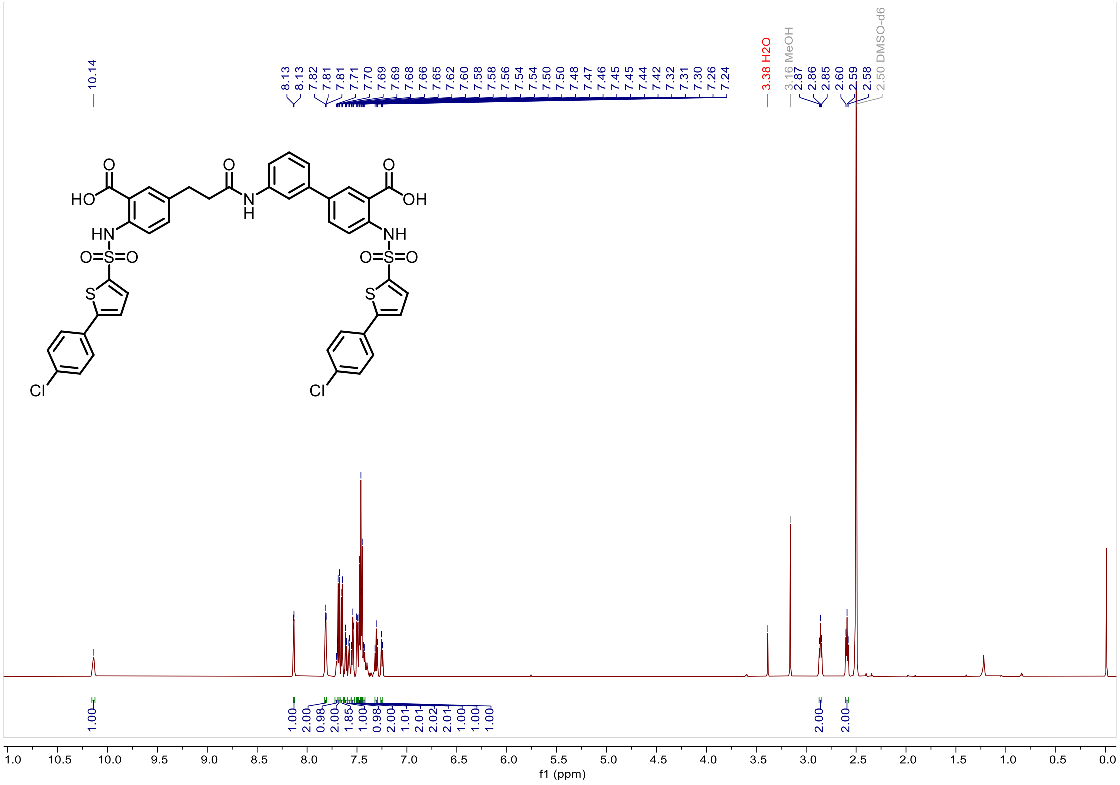

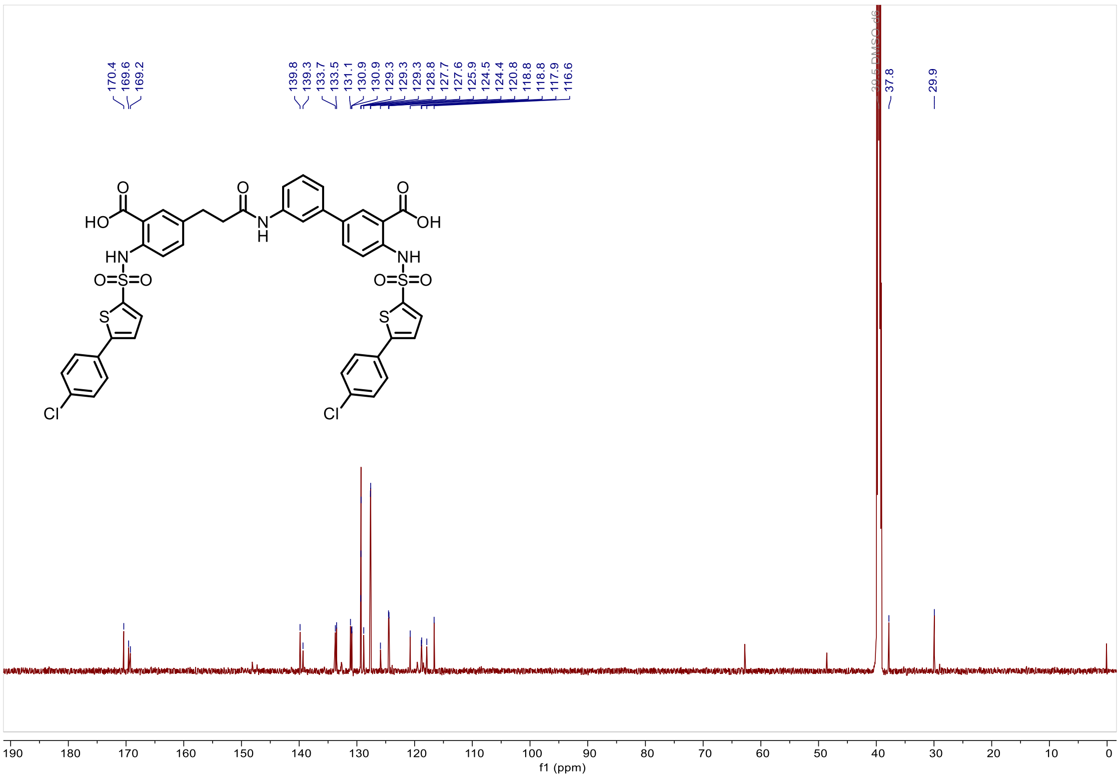

*3'-(3-(3-carboxy-4-((5-(4-chlorophenyl)thiophene)-2-sulfonamido)phenyl)propanamido)-4-((5-(4-chlorophenyl)thiophene)-2-sulfonamido)-[1,1'-biphenyl]-3-carboxylic acid (**In-1**).*

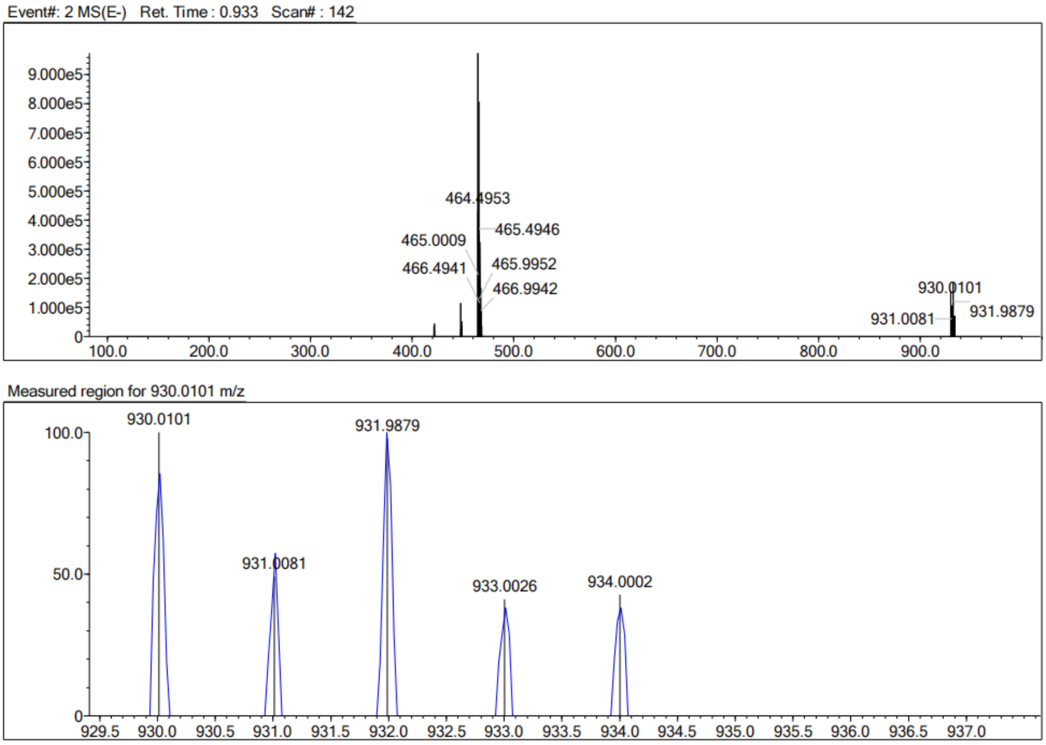


| Rank | Score | Formula (M) | Ion | Meas. m/z | Pred. m/z | Df. (mDa) | Df. (ppm) | Iso | DBE |
|---|---|---|---|---|---|---|---|---|---|
| 1 | 3.37 | C43 H31 N3 O9 S4 Cl2 | [M-H]- | 930.0101 | 930.0247 | -14.6 | -15.70 | 13.60 | 29.0 |

*Ethyl -2-amino-5-(3-(tert-butoxy)-3-oxoprop-1-en-1-yl)benzoate (**C8**).*

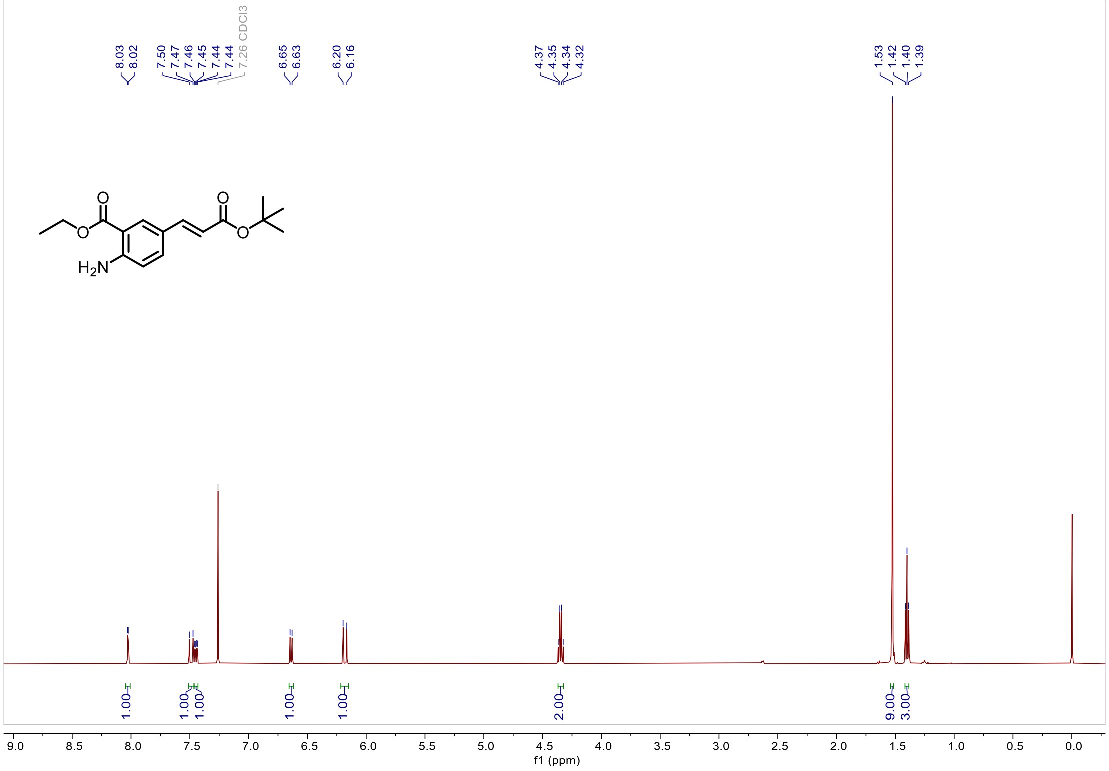

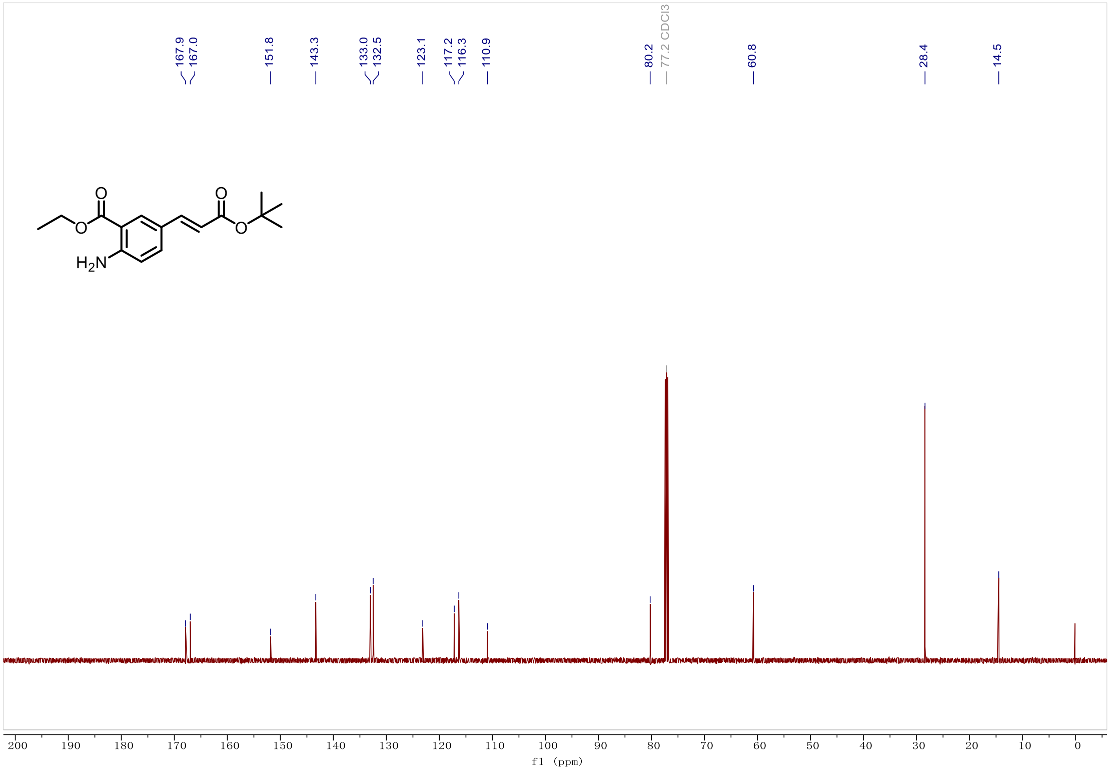

*2-amino-5-(3-(tert-butoxy)-3-oxopropyl)benzoate (**C9**)*

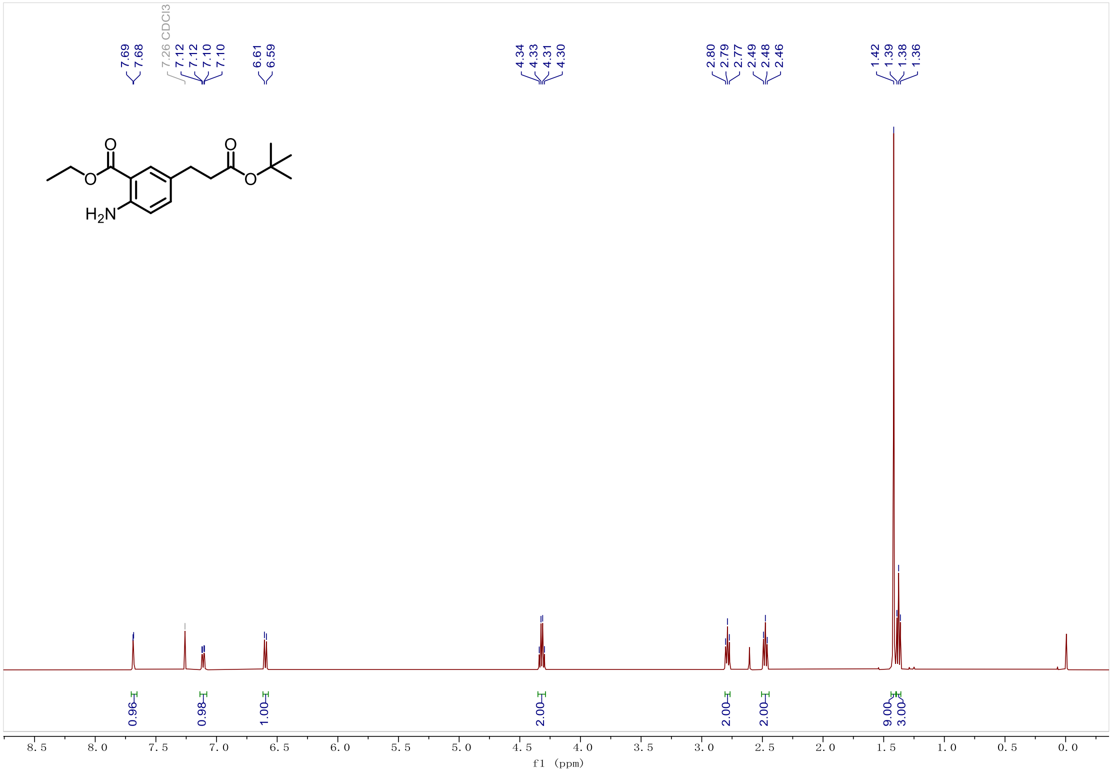


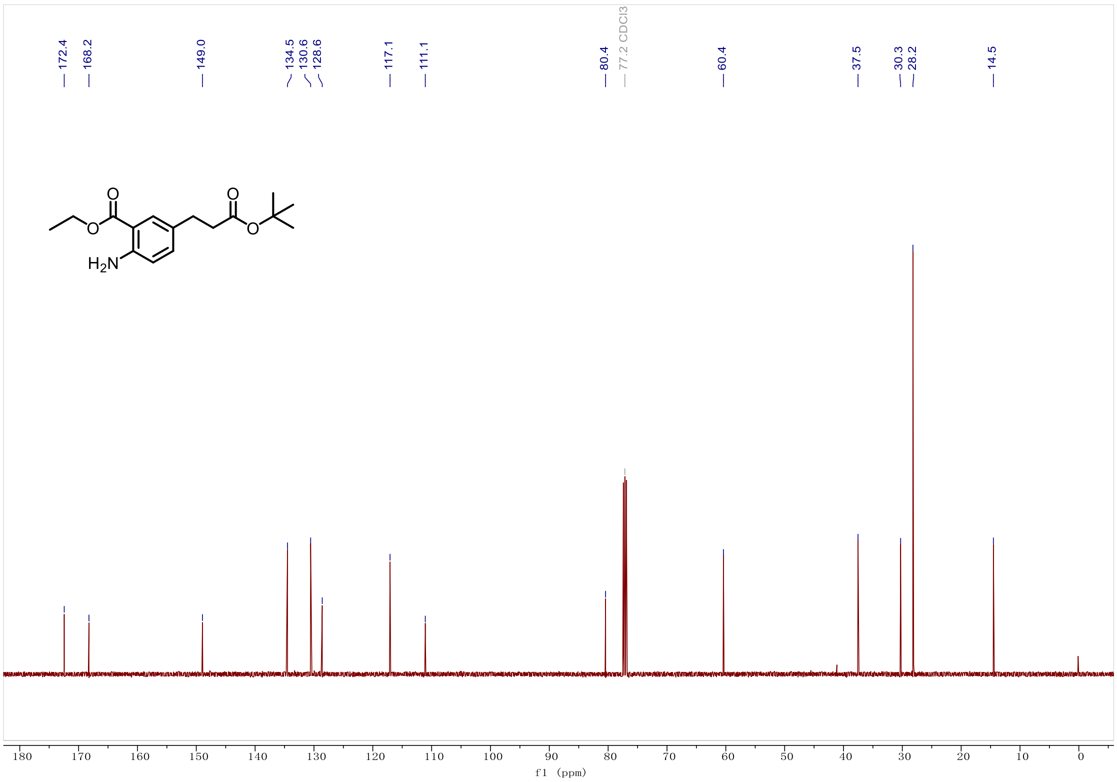

*Ethyl 4-amino-3'-((tert-butoxycarbonyl)amino)-[1,1'-biphenyl]-3-carboxylate(**C10**).*

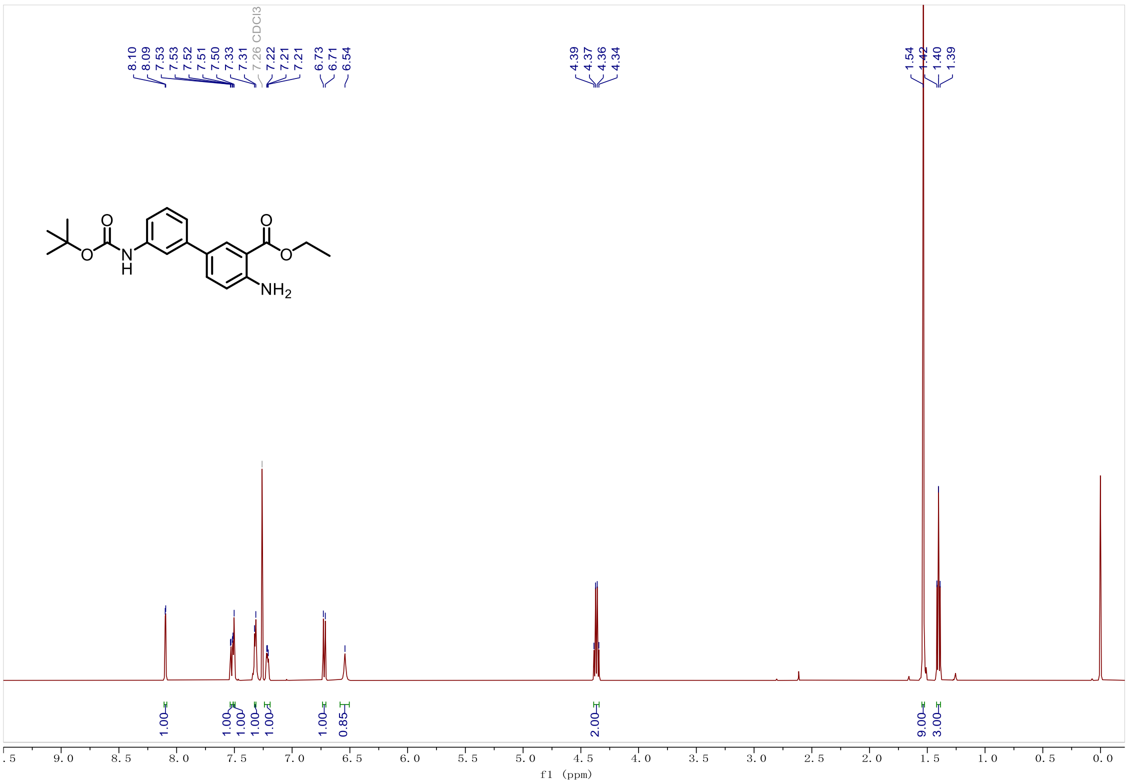


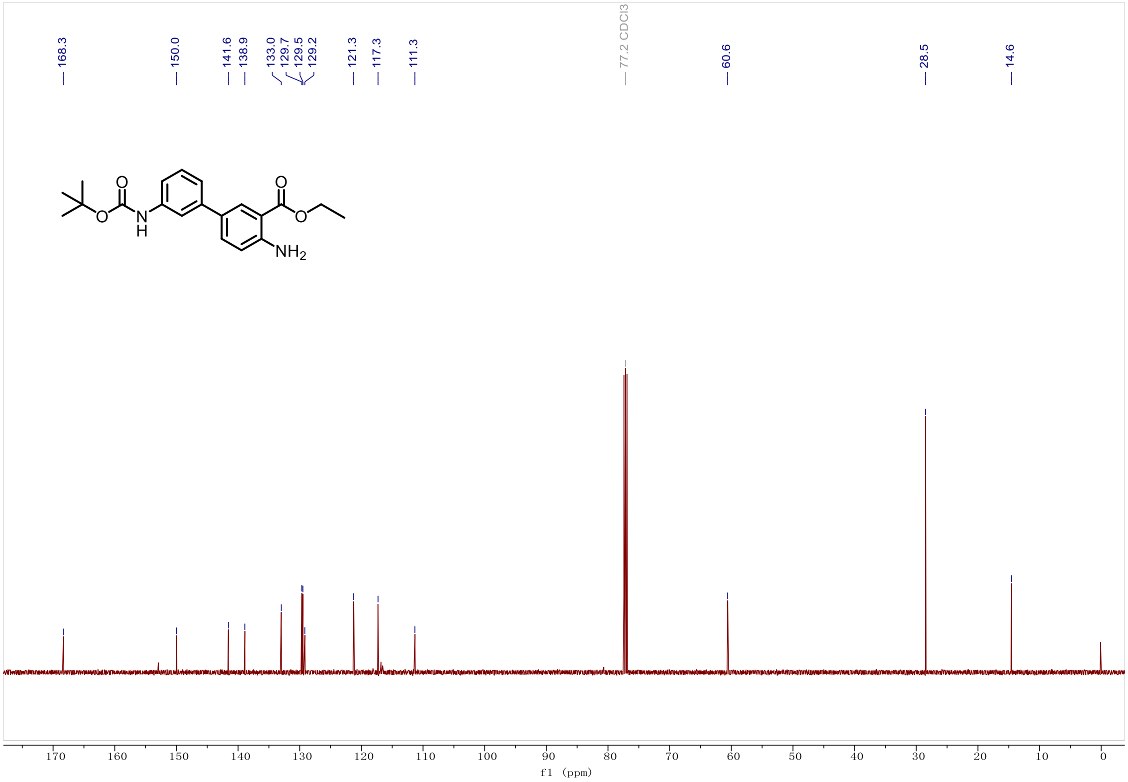

*Ethyl 4-((5-bromothiophene)-2-sulfonamido)-3'-(3-(4-((5-bromothiophene)-2-sulfonamido)-3-(ethoxycarbonyl)phenyl)propanamido)-[1,1'-biphenyl]-3-carboxylate (**C11**).*

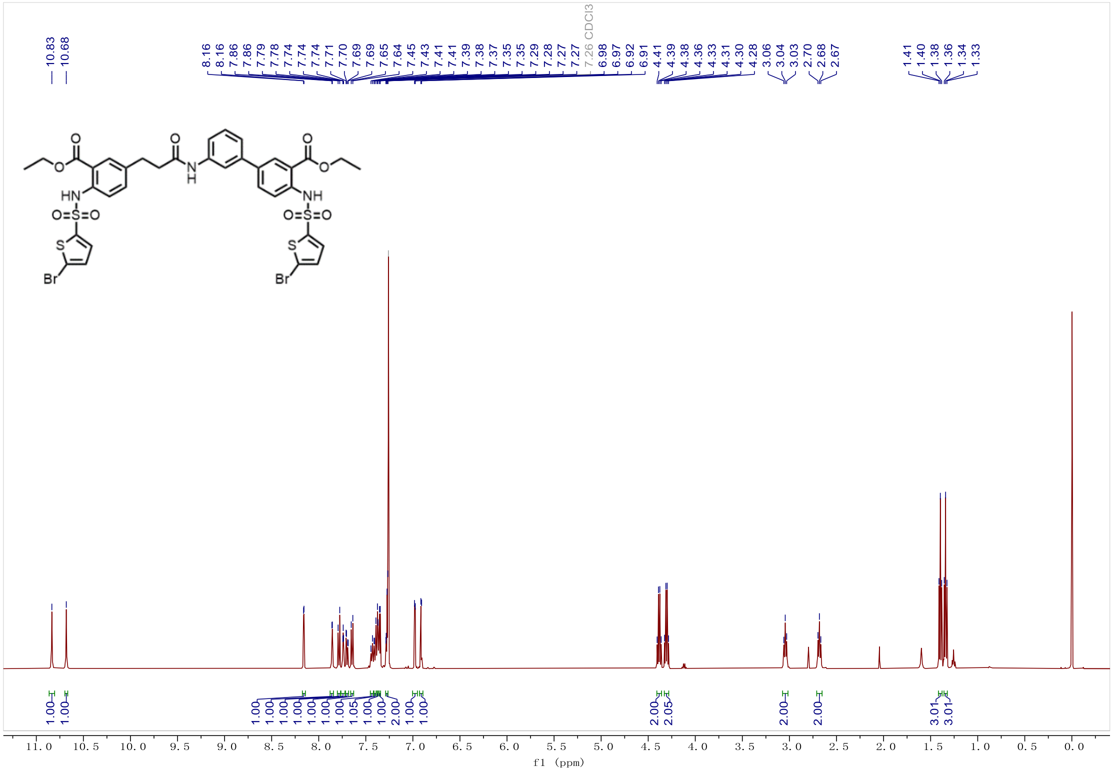

*3'-(3-(3-carboxy-4-((5-phenylthiophene)-2-sulfonamido)phenyl)propanamido)-4-((5-phenylthiophene)-2-sulfonamido)-[1,1'-biphenyl]-3-carboxylic acid (**Y-1**).*

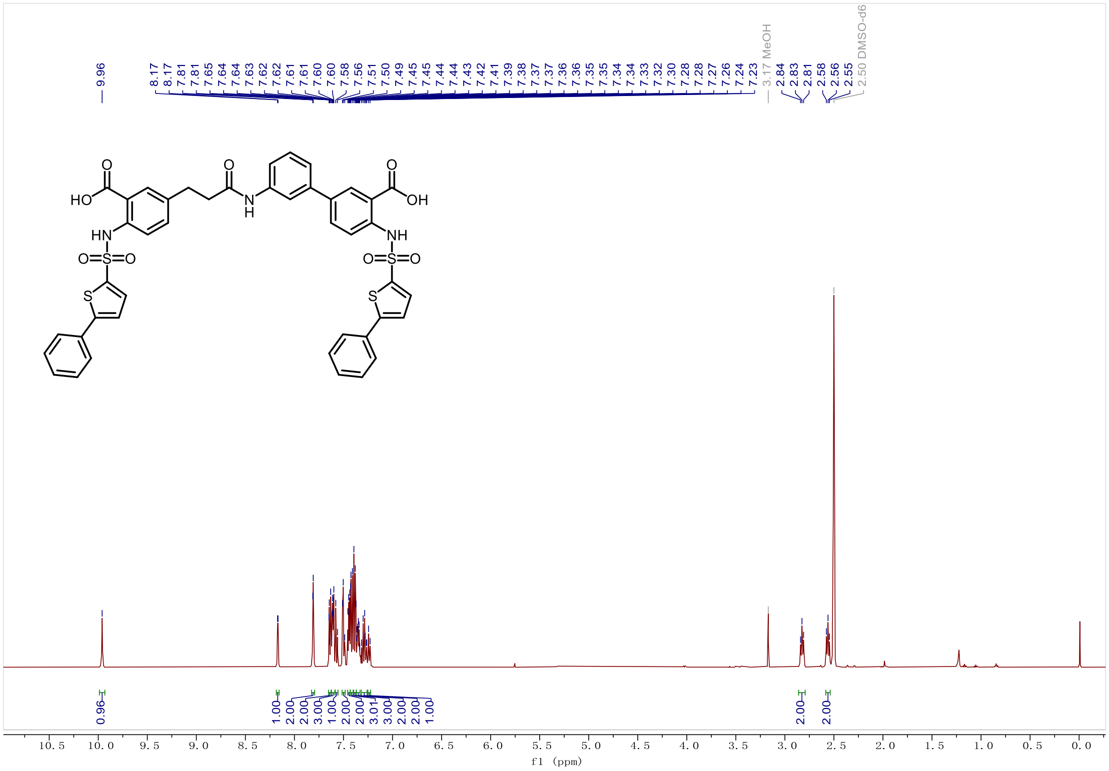

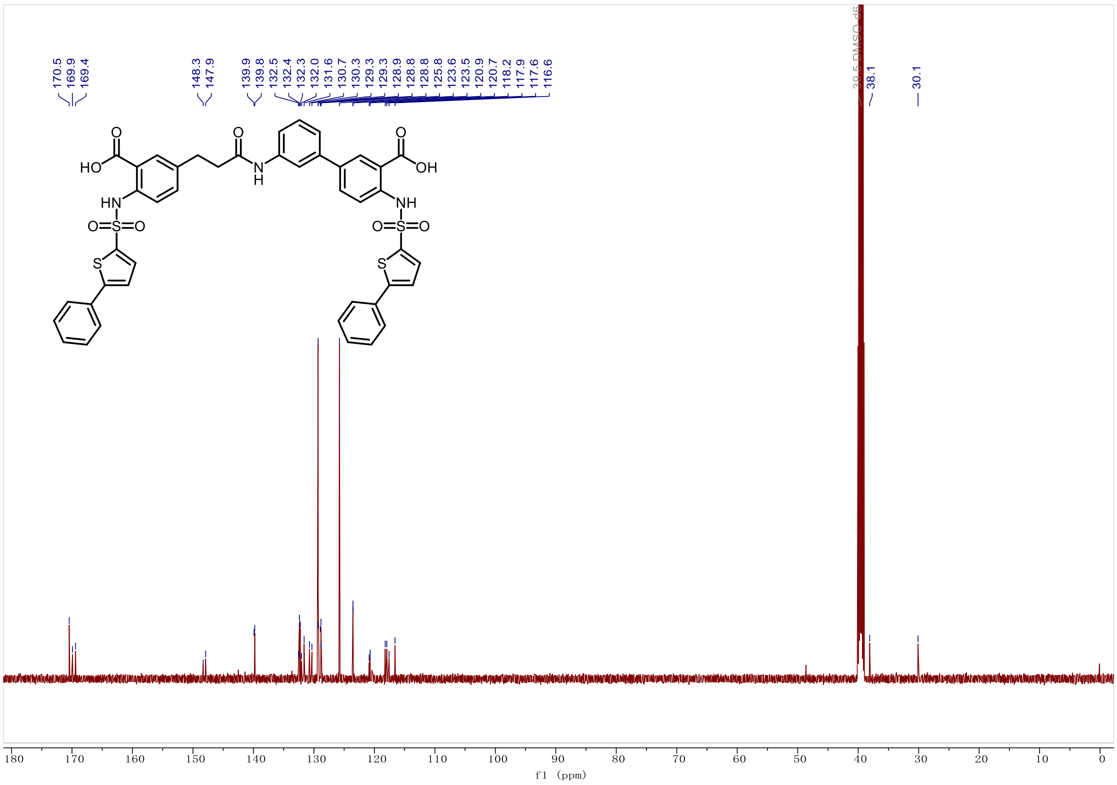

*3'-(3-(3-carboxy-4-((5-phenylthiophene)-2-sulfonamido)phenyl)propanamido)-4-((5-phenylthiophene)-2-sulfonamido)-[1,1'-biphenyl]-3-carboxylic acid (**Y-1**).*

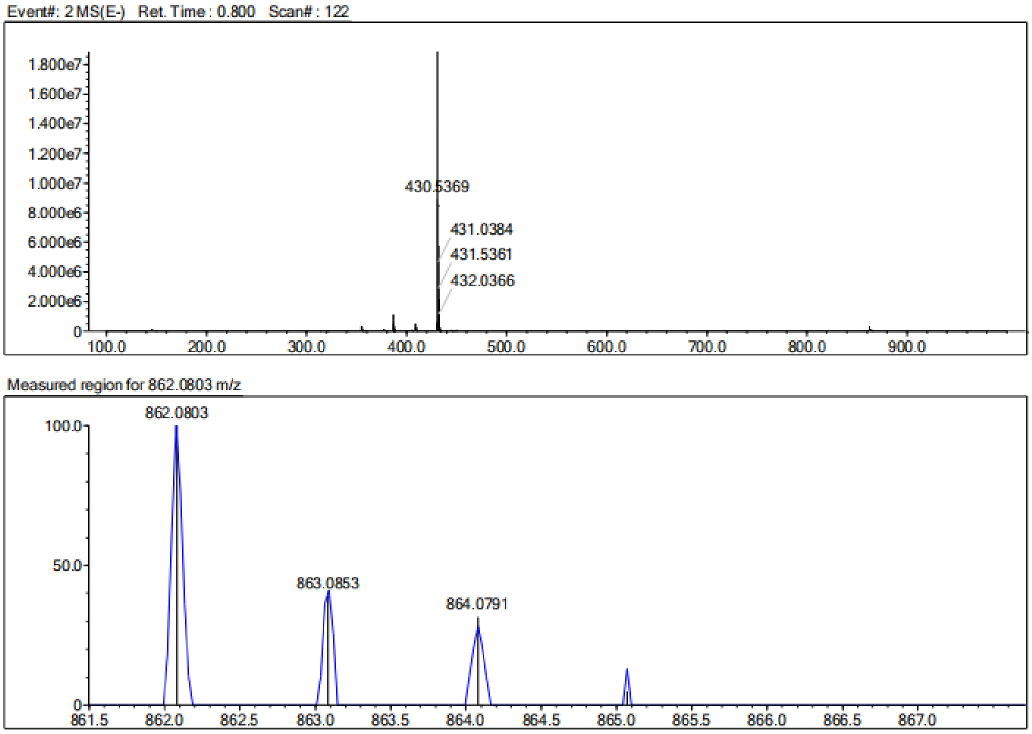


| Rank | Score | Formula (M) | Ion | Meas. m/z | Pred. m/z | Df. (mDa) | Df. (ppm) | Iso | DBE |
|---|---|---|---|---|---|---|---|---|---|
| 1 | 0.00 | C43 H33 N3 O9 S4 | [M-H]- | 862.0803 | 862.1027 | -22.4 | -25.98 | 13.23 | 29.0 |

*3'-(3-(3-carboxy-4-((5-(4-fluorophenyl)thiophene)-2-sulfonamido)phenyl)propanamido)-4-((5-(4-fluorophenyl)thiophene)-2-sulfonamido)-[1,1'-biphenyl]-3-carboxylic acid (**Y-2**).*

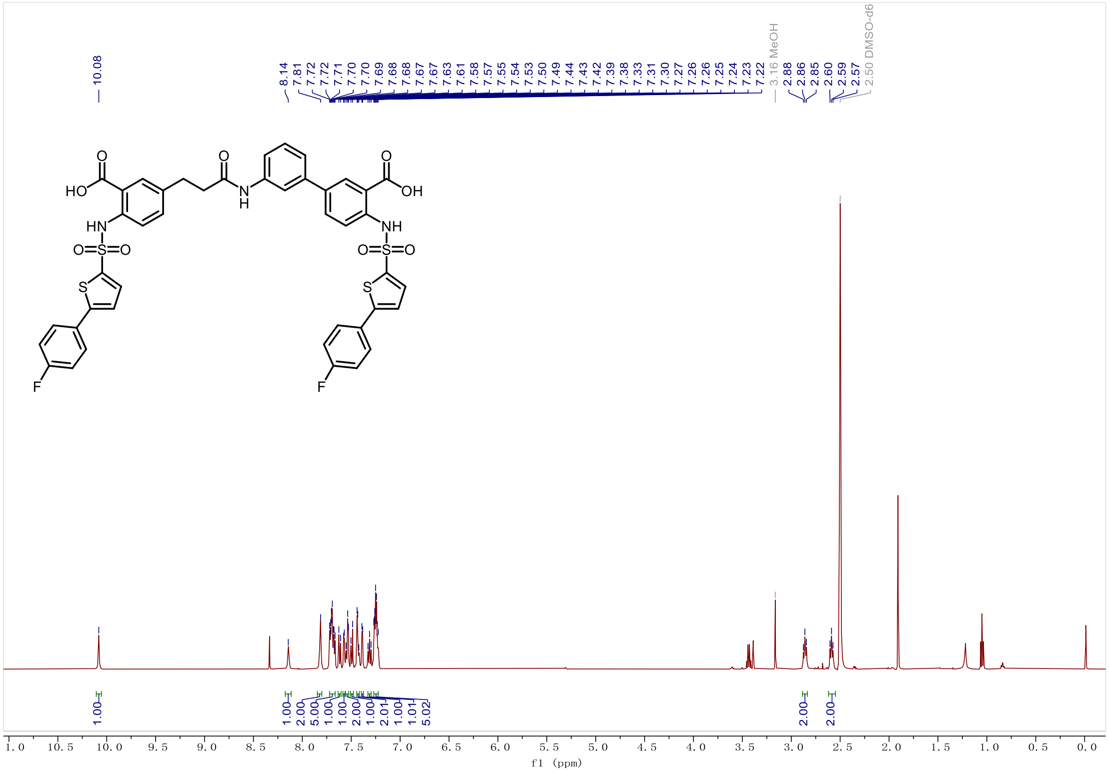

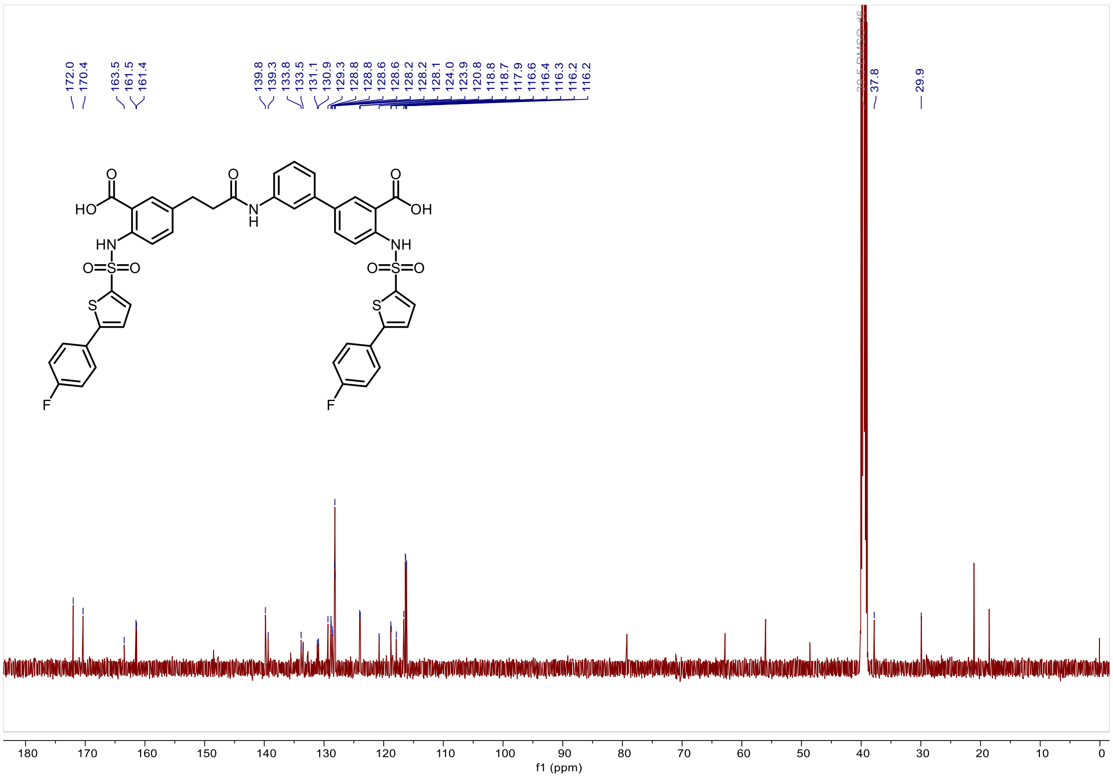

*3'-(3-(3-carboxy-4-((5-(4-fluorophenyl)thiophene)-2-sulfonamido)phenyl)propanamido)-4-((5-(4-fluorophenyl)thiophene)-2-sulfonamido)-[1,1'-biphenyl]-3-carboxylic acid (**Y-2**).*

Event#: 2 MS(E-) Ret. Time : 2.173 Scan# : 328

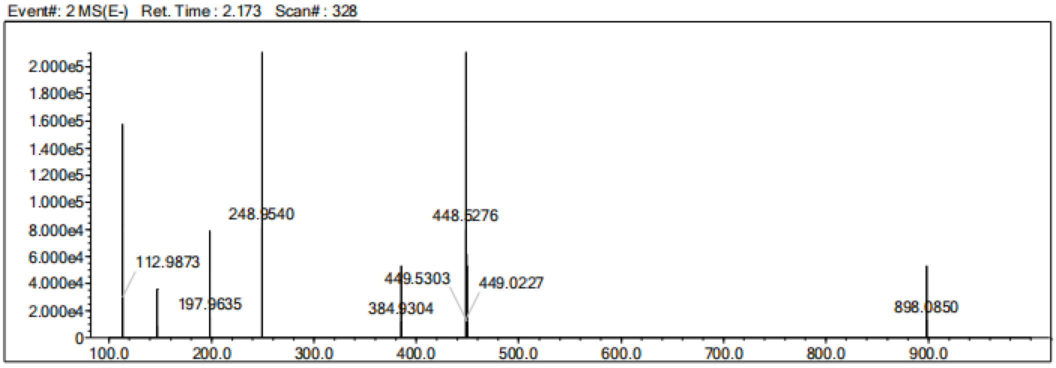


Measured region for 898.0850 m/z

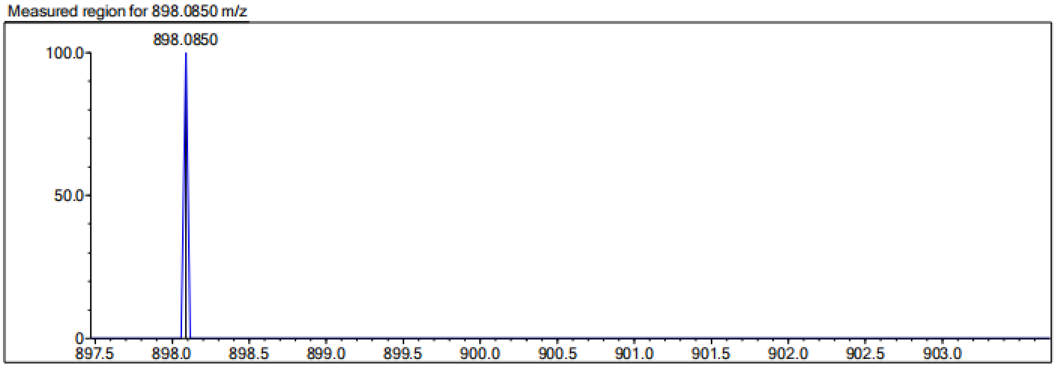


| Rank | Score | Formula (M) | Ion | Meas. m/z | Pred. m/z | Df. (mDa) | Df. (ppm) | Iso | DBE |
|---|---|---|---|---|---|---|---|---|---|
| 1 | 0.00 | C43 H31 N3 O9 F2 S4 | [M-H]- | 898.0850 | 898.0838 | 1.2 | 1.34 | 0.00 | 29.0 |

*3'-(3-(3-carboxy-4-((5-(4-nitrophenyl)thiophene)-2-sulfonamido)phenyl)propanamido)-4-((5-(4-nitrophenyl)thiophene)-2-sulfonamido)-[1,1'-biphenyl]-3-carboxylic acid (**Y-3**).*

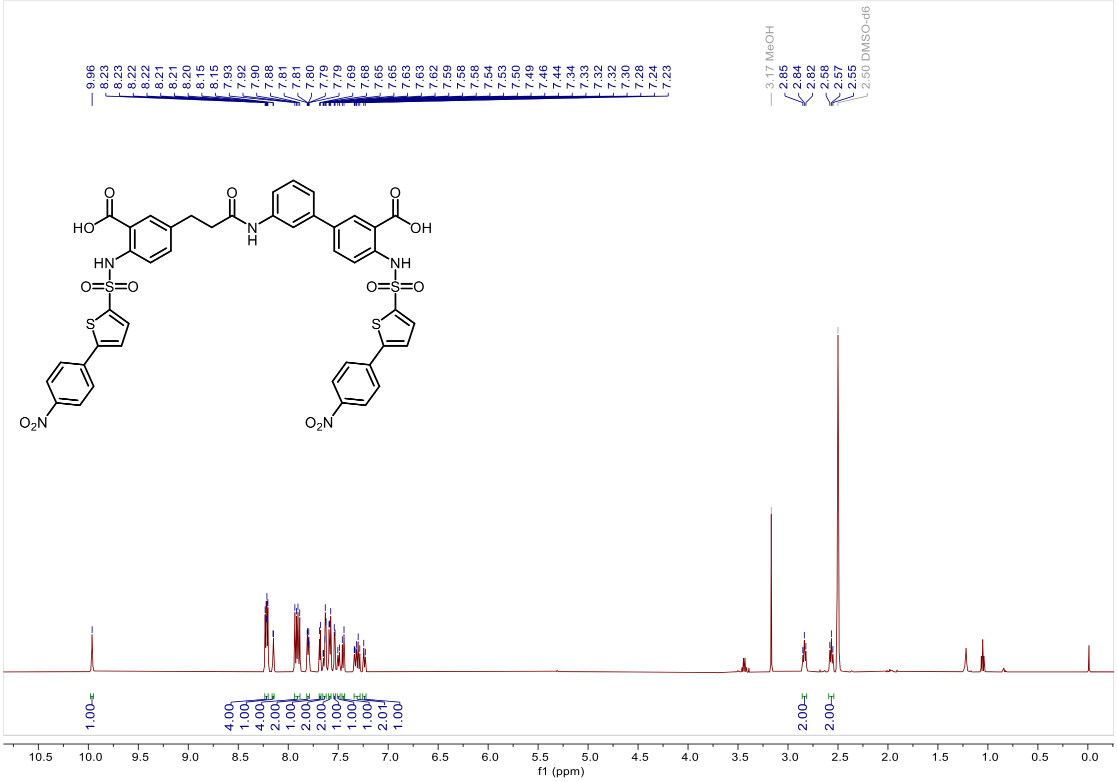


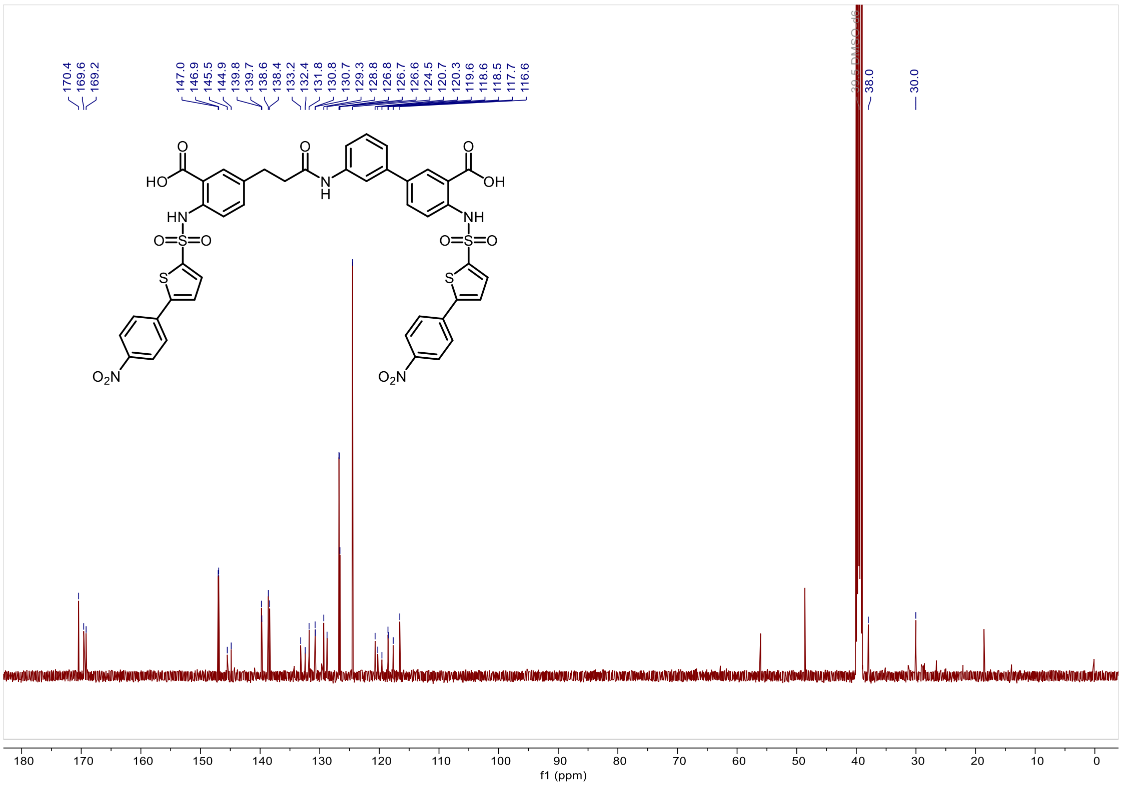

*3'-(3-(3-carboxy-4-((5-(4-nitrophenyl)thiophene)-2-sulfonamido)phenyl)propanamido)-4-((5-(4-nitrophenyl)thiophene)-2-sulfonamido)-[1,1'-biphenyl]-3-carboxylic acid (**Y-3**).*

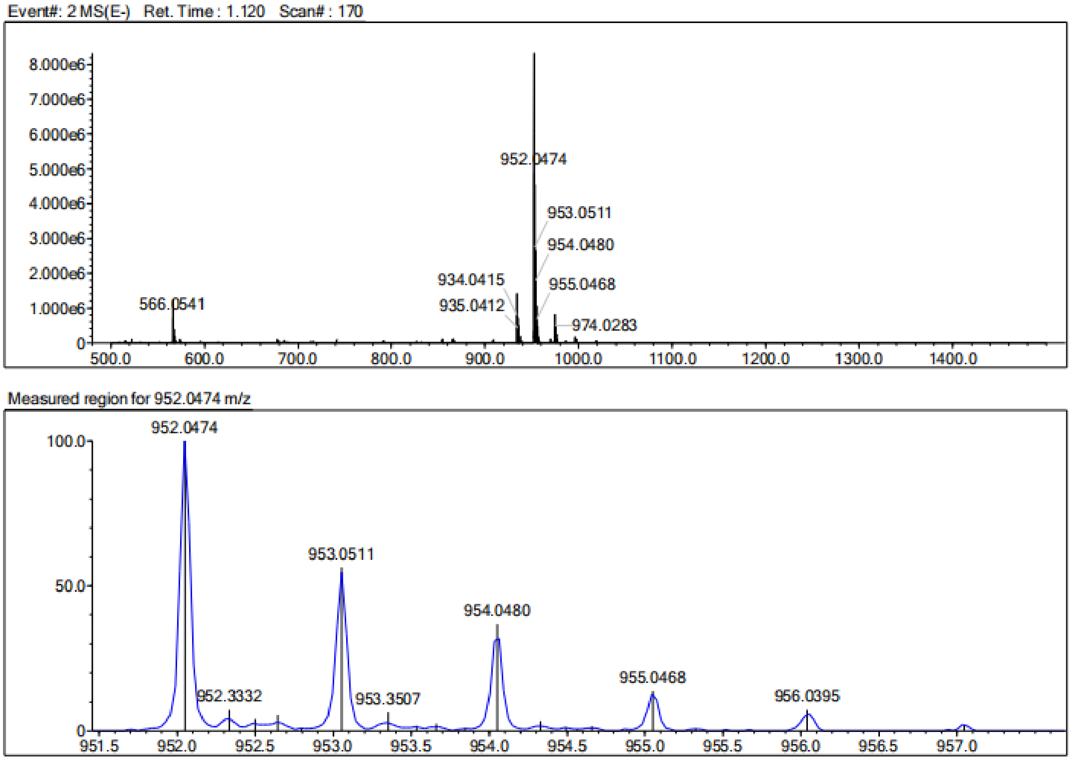


| Rank | Score | Formula (M) | Ion | Meas. m/z | Pred. m/z | Df. (mDa) | Df. (ppm) | Iso | DBE |
|---|---|---|---|---|---|---|---|---|---|
| 1 | 0.00 | C43 H31 N5 O13 S4 | [M-H]- | 952.0474 | 952.0728 | -25.4 | -26.68 | 33.11 | 31.0 |

*3'-(3-(3-carboxy-4-((5-(4-hydroxyphenyl)thiophene)-2-sulfonamido)phenyl)propanamido)-4-((5-(4-hydroxyphenyl)thiophene)-2-sulfonamido)-[1,1'-biphenyl]-3-carboxylic acid (**Y-4**).*

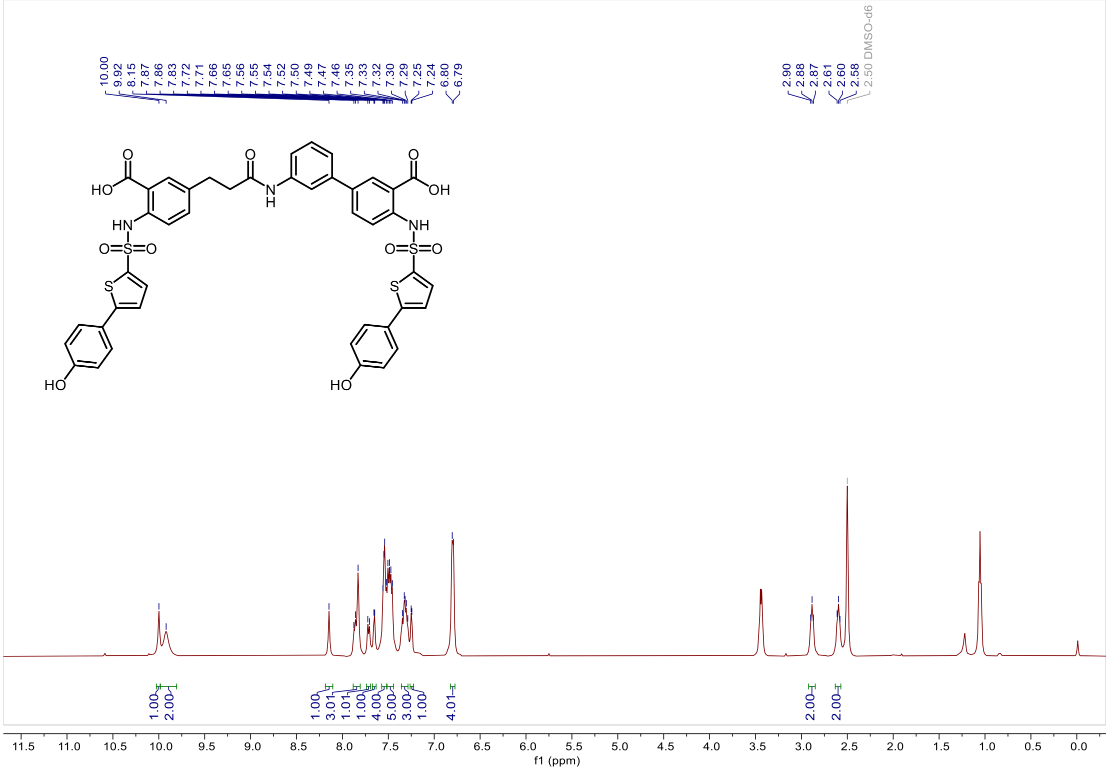


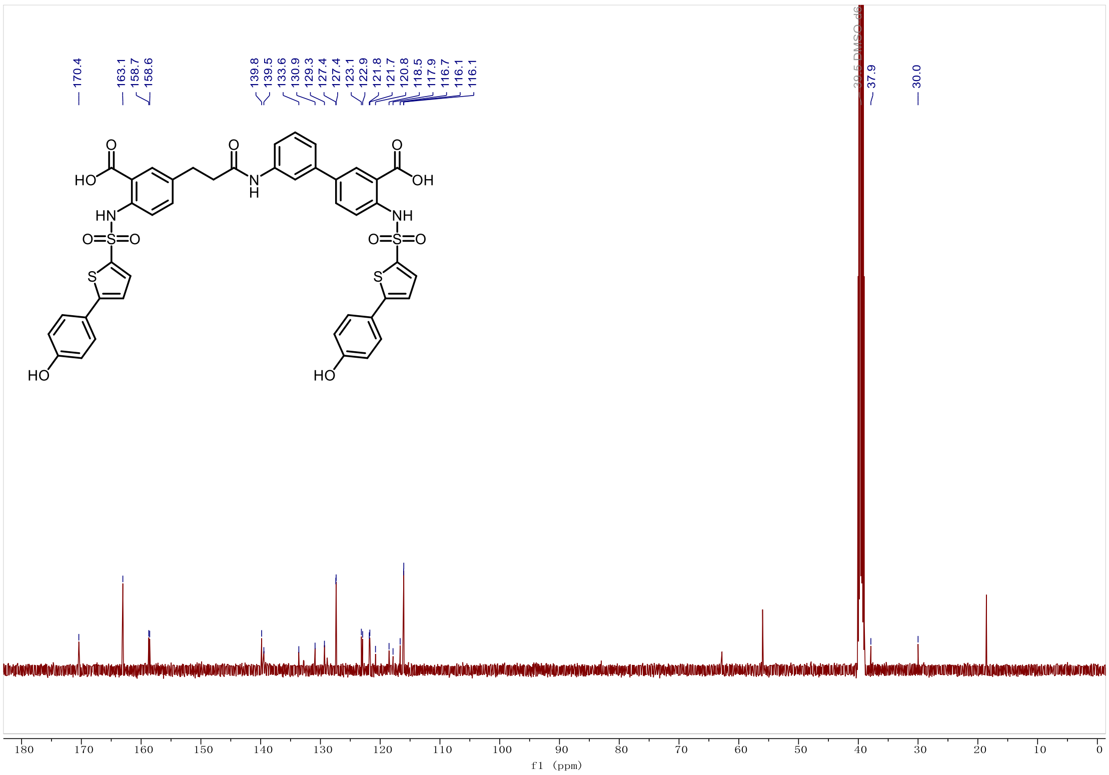

*3'-(3-(3-carboxy-4-((5-(4-hydroxyphenyl)thiophene)-2-sulfonamido)phenyl)propanamido)-4-((5-(4-hydroxyphenyl)thiophene)-2-sulfonamido)-[1,1'-biphenyl]-3-carboxylic acid (**Y-4**).*

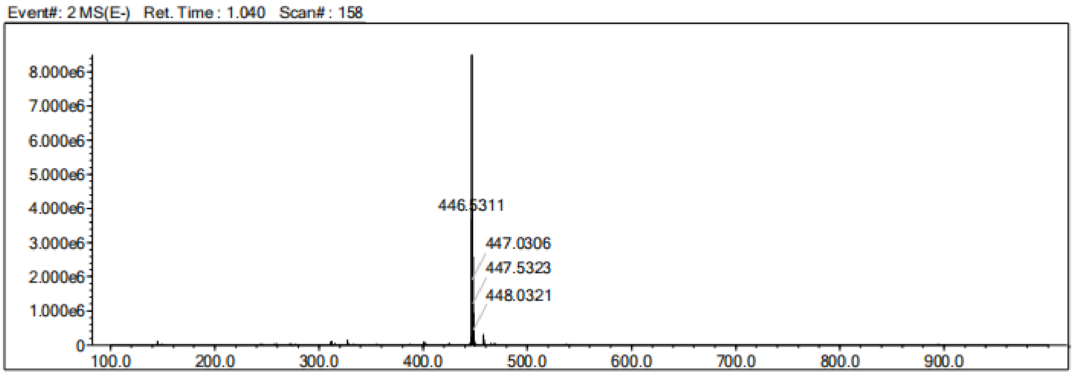


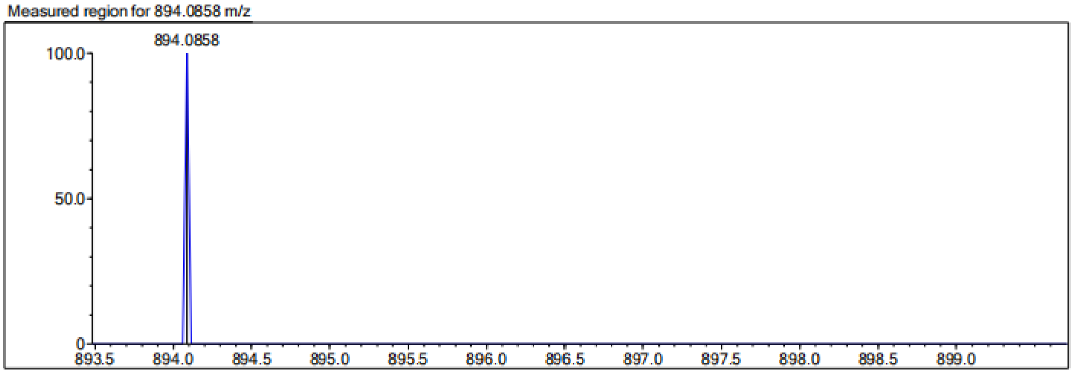


| Rank | Score | Formula (M) | Ion | Meas. m/z | Pred. m/z | Df. (mDa) | Df. (ppm) | Iso | DBE |
|---|---|---|---|---|---|---|---|---|---|
| 1 | 0.00 | C43 H33 N3 O11 S4 | [M-H]- | 894.0858 | 894.0925 | -6.7 | -7.49 | 0.00 | 29.0 |

*3'-(3-(3-carboxy-4-((5-(4-methoxyphenyl)thiophene)-2-sulfonamido)phenyl)propanamido)-4-((5-(4-methoxyphenyl)thiophene)-2-sulfonamido)-[1,1'-biphenyl]-3-carboxylic acid* (***Y-5***).

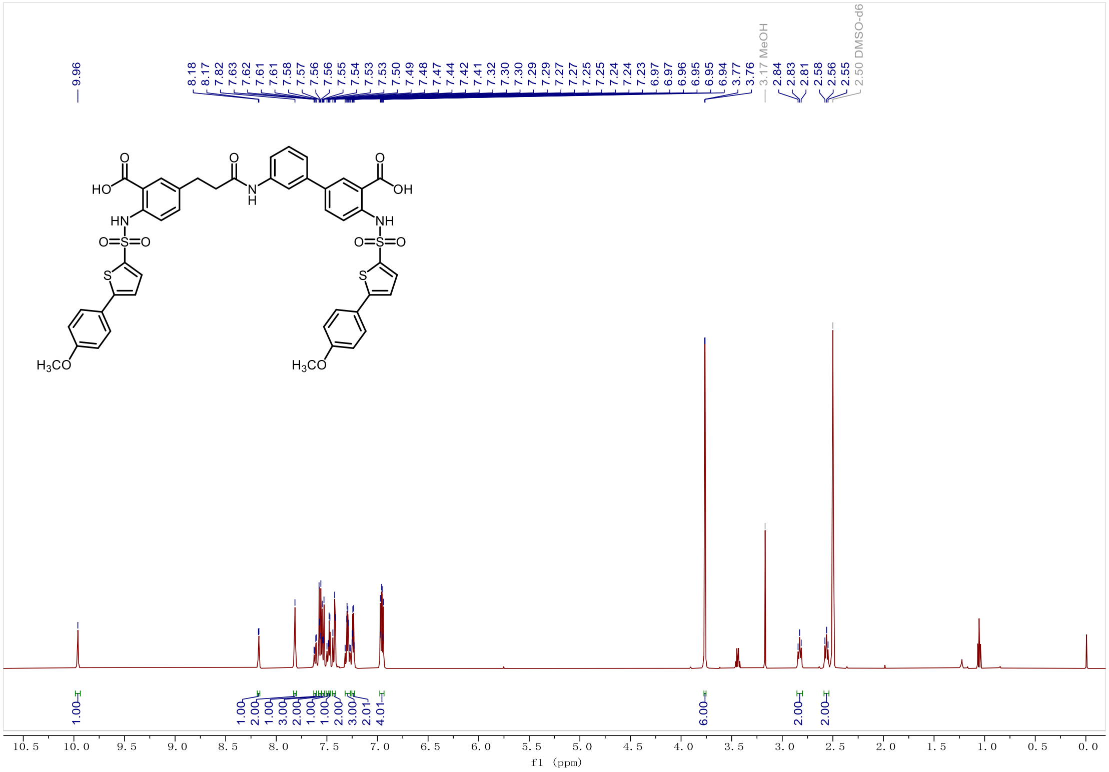

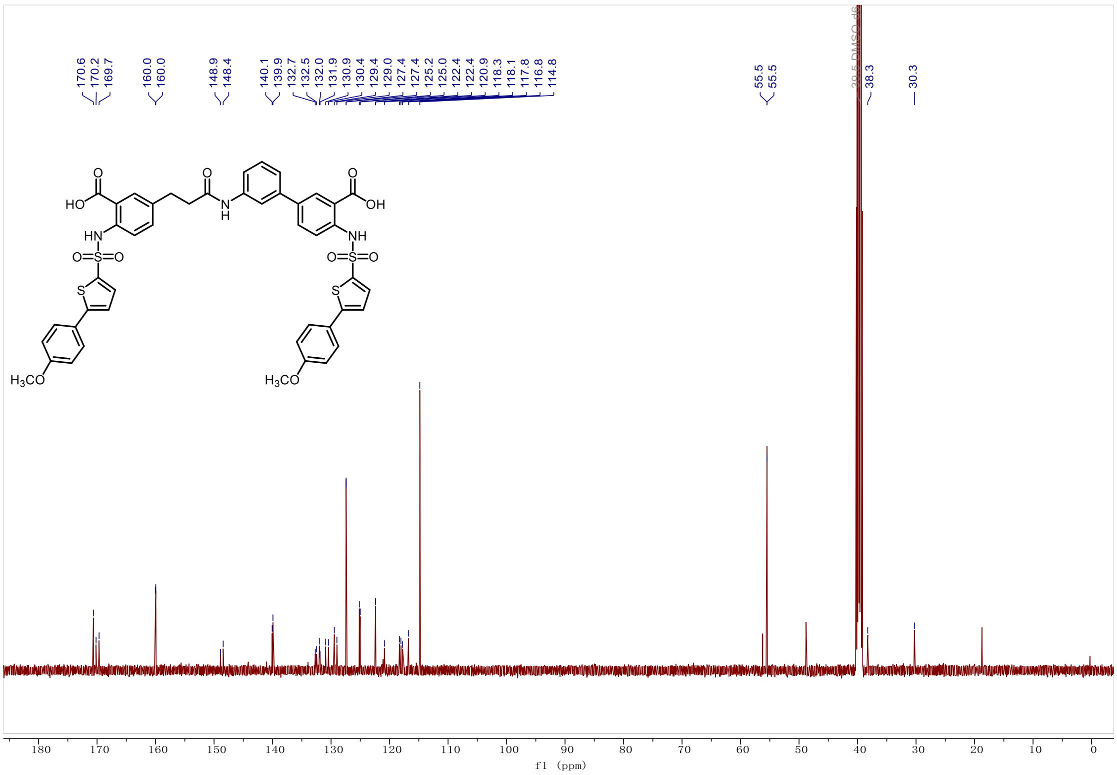

*3'-(3-(3-carboxy-4-((5-(4-methoxyphenyl)thiophene)-2-sulfonamido)phenyl)propanamido)-4-((5-(4-methoxyphenyl)thiophene)-2-sulfonamido)-[1,1'-biphenyl]-3-carboxylic acid (**Y-5**).*

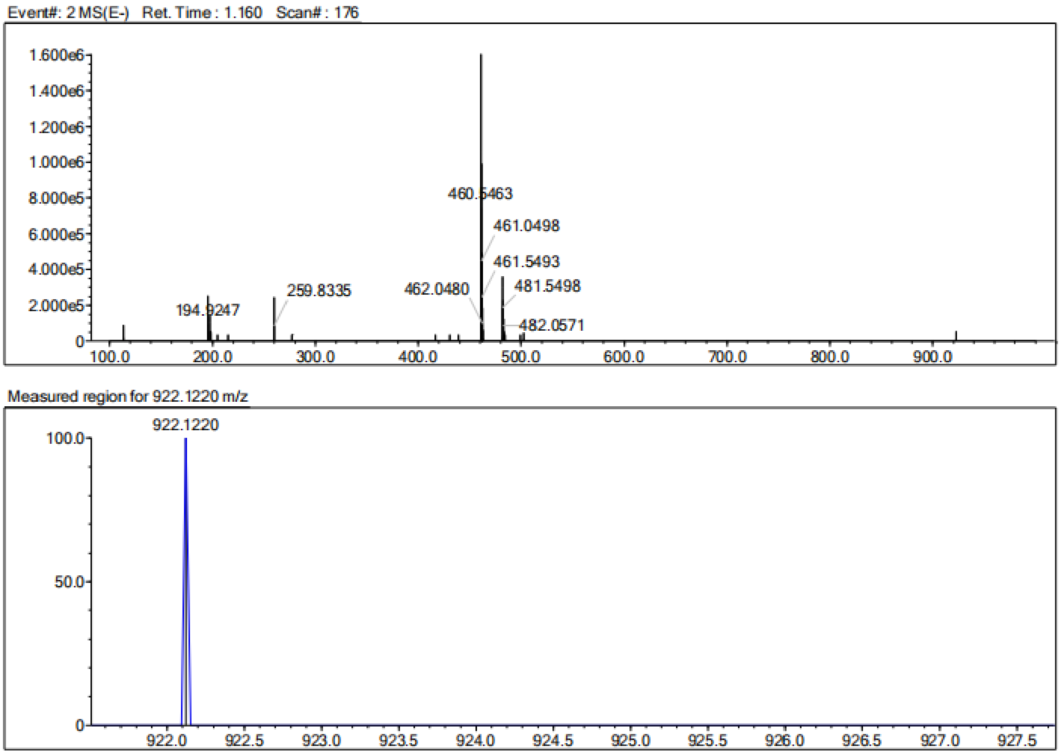


| Rank | Score | Formula (M) | Ion | Meas. m/z | Pred. m/z | Df. (mDa) | Df. (ppm) | Iso | DBE |
|---|---|---|---|---|---|---|---|---|---|
| 1 | 0.00 | C45 H37 N3 O11 S4 | [M-H]- | 922.1220 | 922.1238 | -1.8 | -1.95 | 0.00 | 29.0 |

*3'-(3-(3-carboxy-4-((5-(3-fluorophenyl)thiophene)-2-sulfonamido)phenyl)propanamido)-4-((5-(3-fluorophenyl)thiophene)-2-sulfonamido)-[1,1'-biphenyl]-3-carboxylic acid (**Y-6**).*

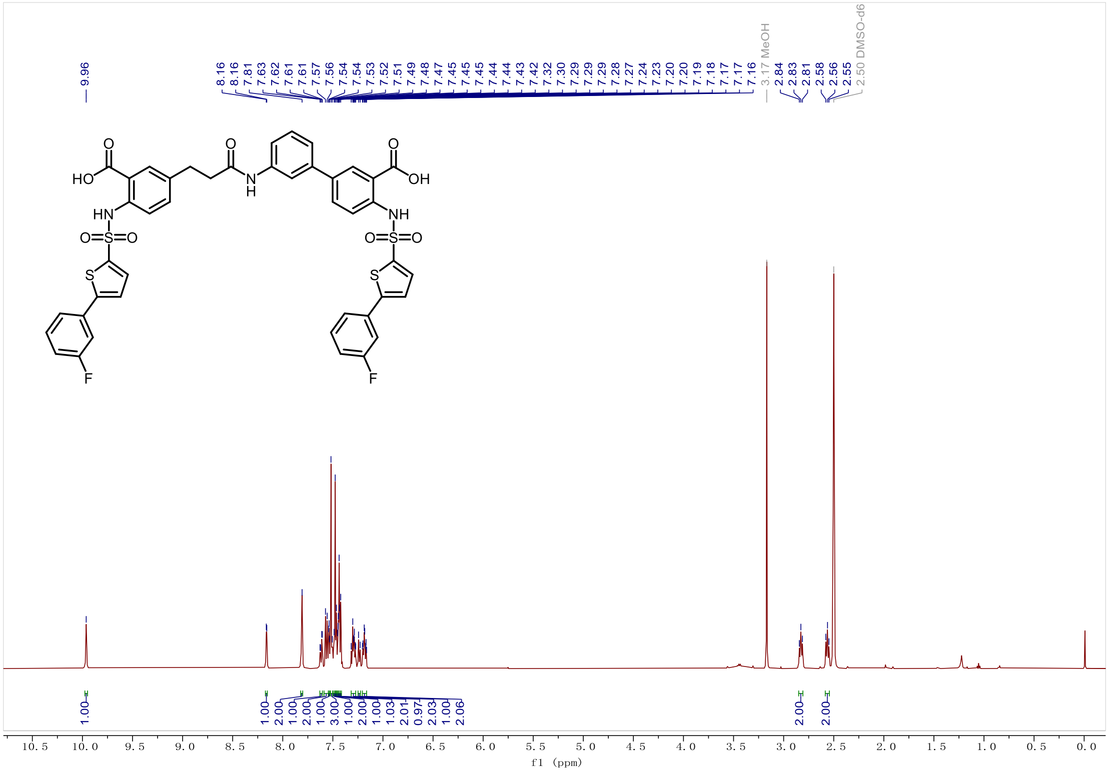

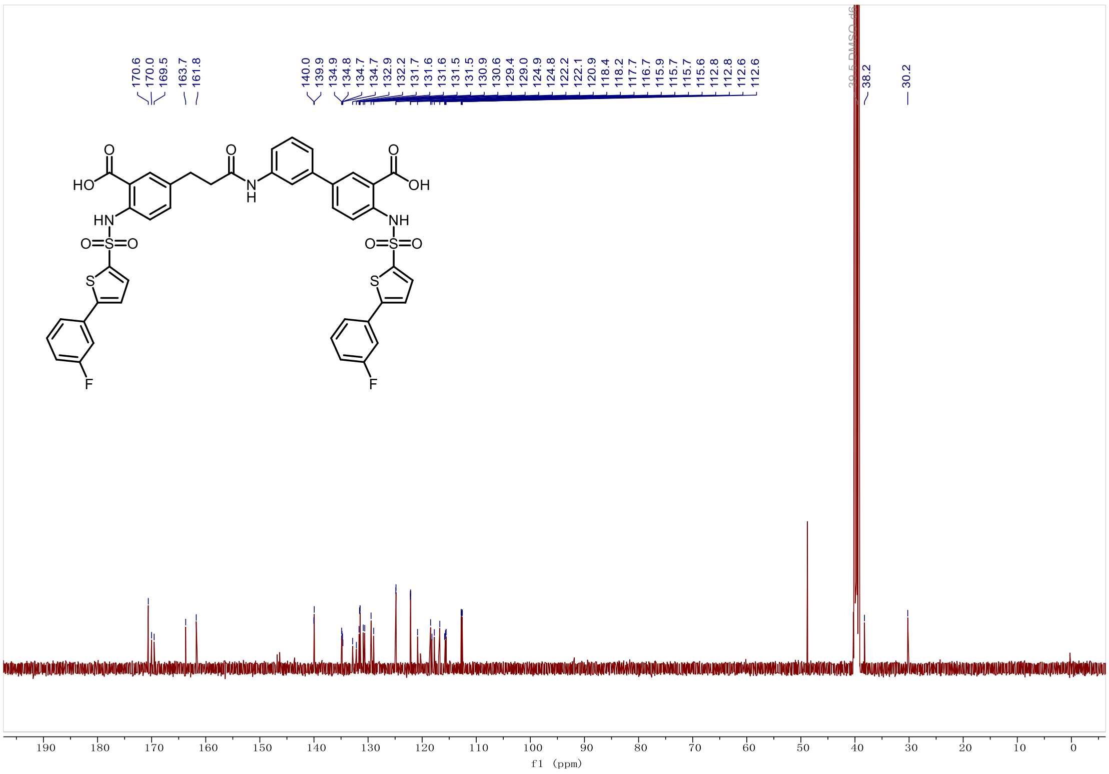

*3'-(3-(3-carboxy-4-((5-(3-fluorophenyl)thiophene)-2-sulfonamido)phenyl)propanamido)-4-((5-(3-fluorophenyl)thiophene)-2-sulfonamido)-[1,1'-biphenyl]-3-carboxylic acid (**Y-6**).*

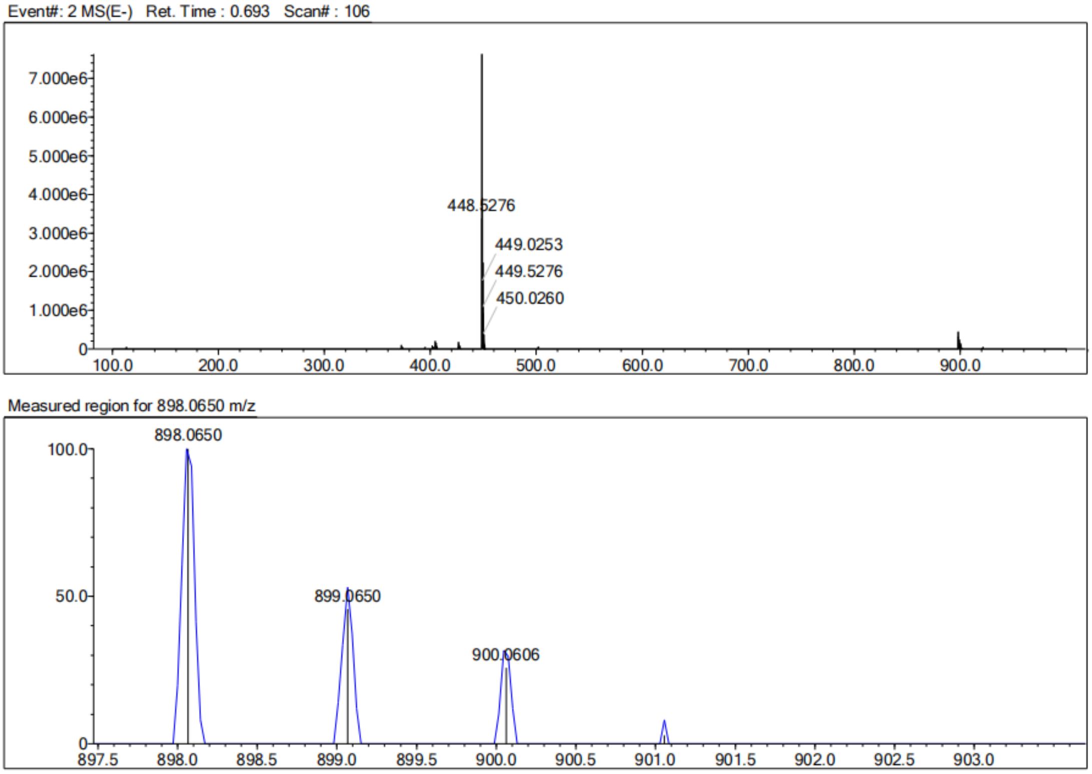


| Rank | Score | Formula (M) | Ion | Meas. m/z | Pred. m/z | Df. (mDa) | Df. (ppm) | Iso | DBE |
|---|---|---|---|---|---|---|---|---|---|
| 1 | 2.99 | C43 H31 N3 O9 F2 S4 | [M-H]- | 898.0650 | 898.0838 | -18.8 | -20.93 | 27.52 | 29.0 |

*3'-(3-(3-carboxy-4-((5-(3-chlorophenyl)thiophene)-2-sulfonamido)phenyl)propanamido)-4-((5-(3-chlorophenyl)thiophene)-2-sulfonamido)-[1,1'-biphenyl]-3-carboxylic acid (**Y-7**).*

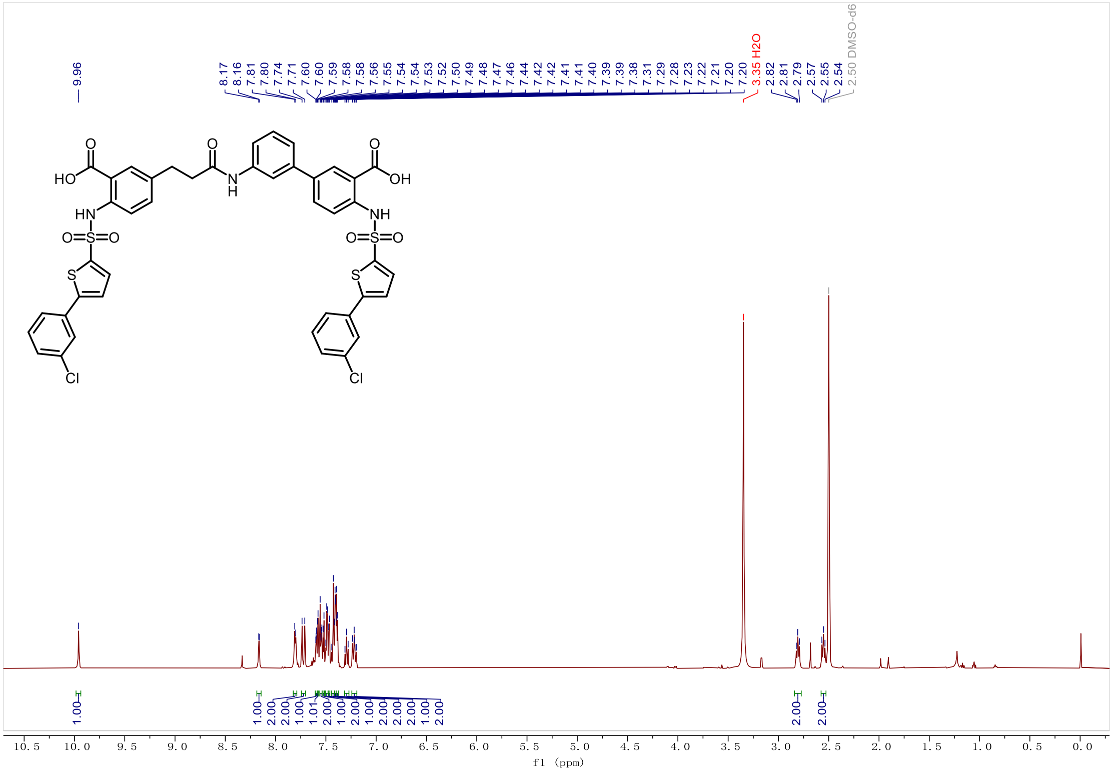


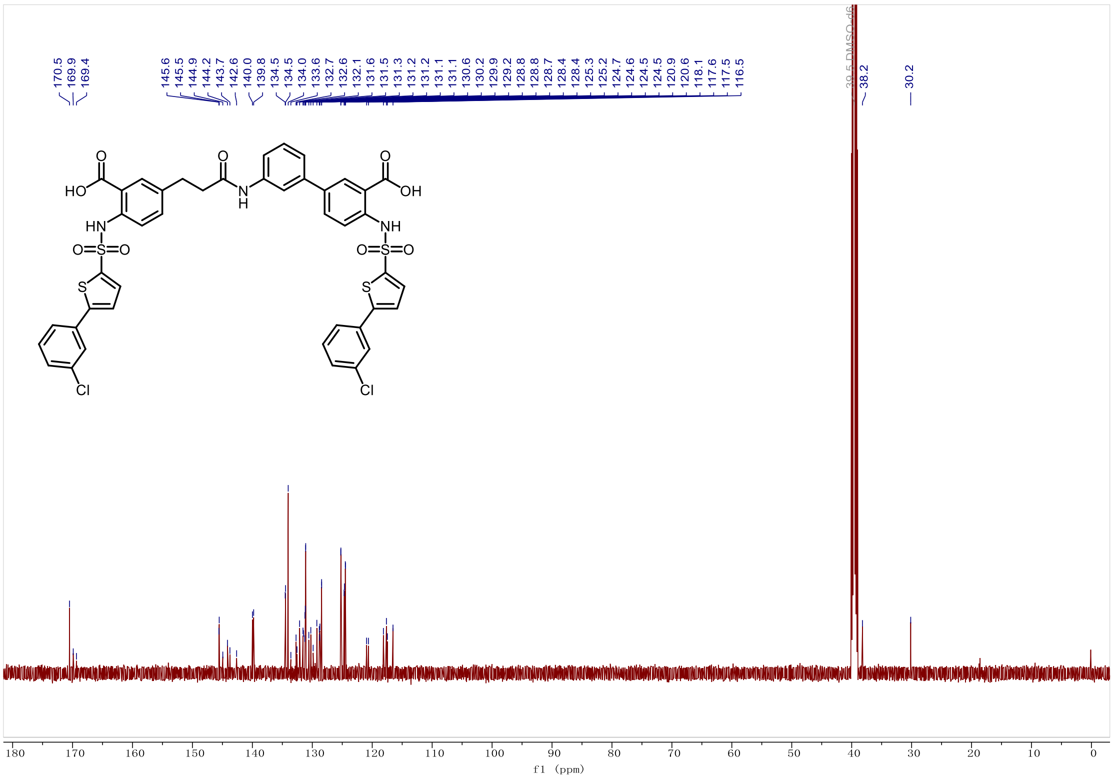

*3'-(3-(3-carboxy-4-((5-(3-chlorophenyl)thiophene)-2-sulfonamido)phenyl)propanamido)-4-((5-(3-chlorophenyl)thiophene)-2-sulfonamido)-[1,1'-biphenyl]-3-carboxylic acid (**Y-7**).*

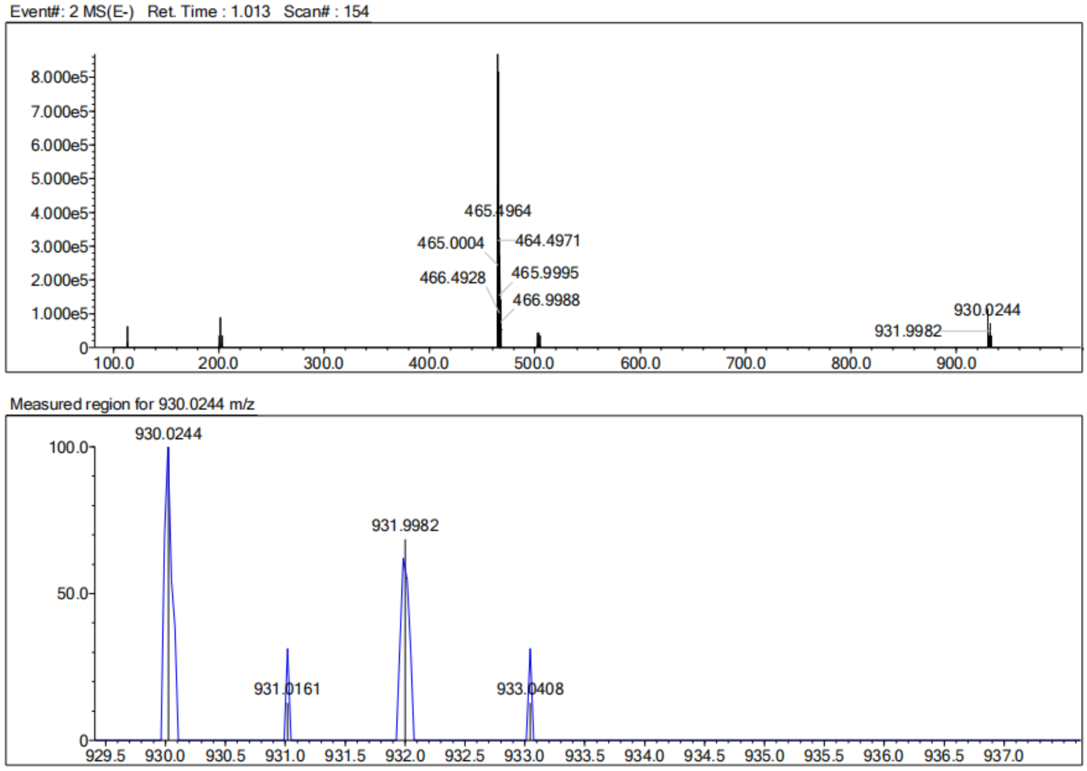


| Rank | Score | Formula (M) | Ion | Meas. m/z | Pred. m/z | Df. (mDa) | Df. (ppm) | Iso | DBE |
|---|---|---|---|---|---|---|---|---|---|
| 1 | 0.00 | C43 H31 N3 O9 S4 Cl2 | [M-H]- | 930.0244 | 930.0247 | -0.3 | -0.32 | 0.00 | 29.0 |

*3'-(3-(3-carboxy-4-((5-(3-nitrophenyl)thiophene)-2-sulfonamido)phenyl)propanamido)-4-((5-(3-nitrophenyl)thiophene)-2-sulfonamido)-[1,1'-biphenyl]-3-carboxylic acid **(Y-8)***

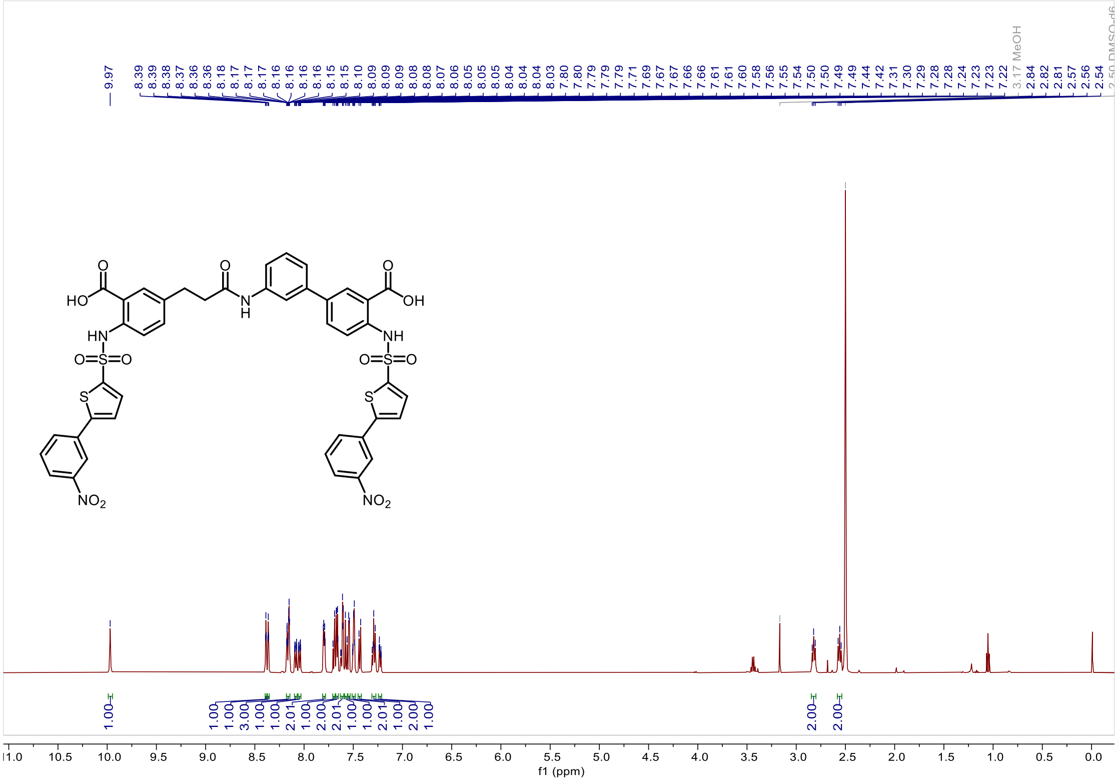


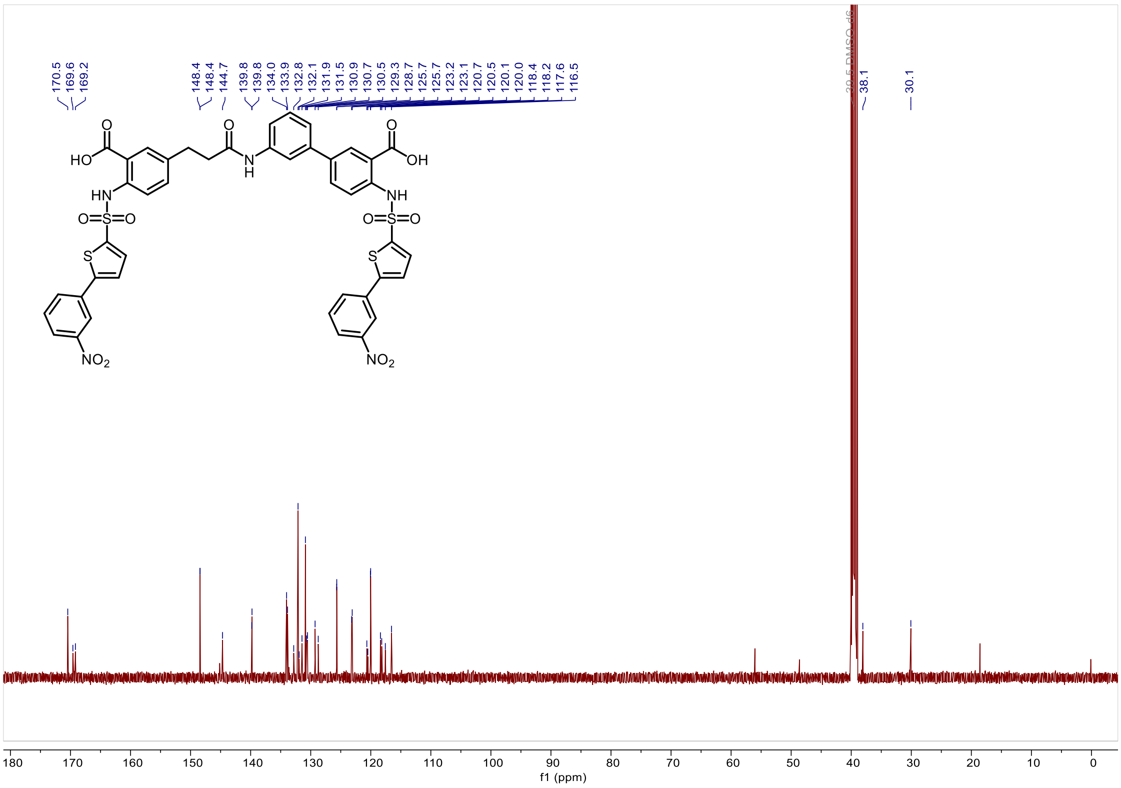

*3'-(3-(3-carboxy-4-((5-(3-nitrophenyl)thiophene)-2-sulfonamido)phenyl)propanamido)-4-((5-(3-nitrophenyl)thiophene)-2-sulfonamido)-[1,1'-biphenyl]-3-carboxylic acid (**Y-8**)*

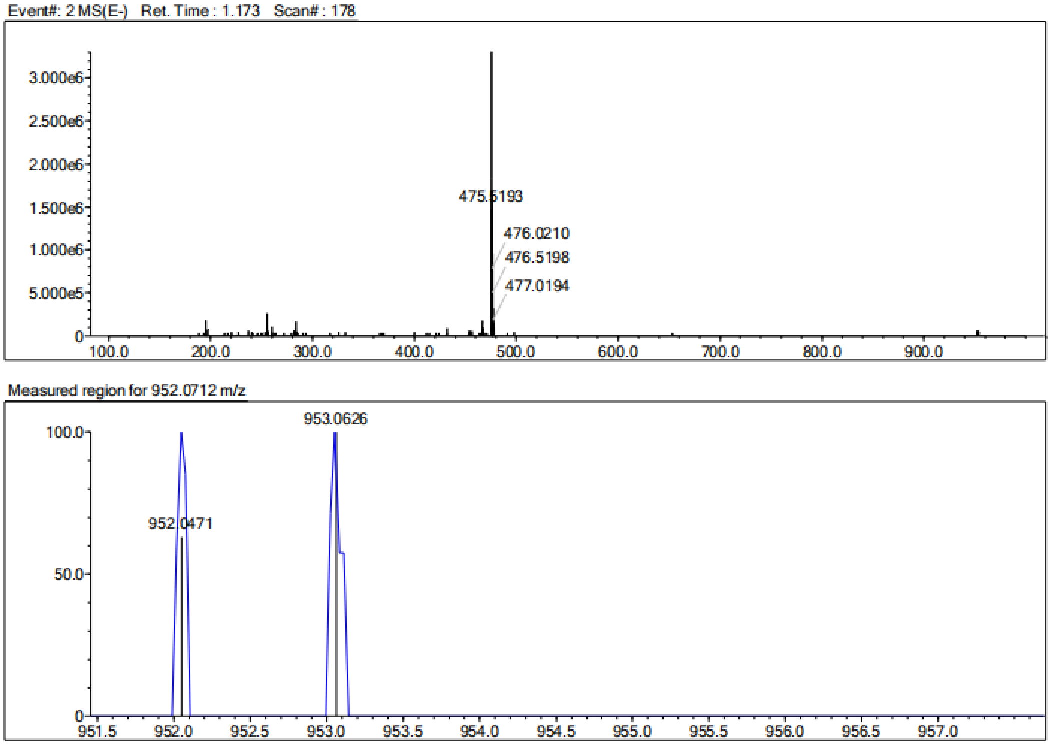


| Rank | Score | Formula (M) | Ion | Meas. m/z | Pred. m/z | Df. (mDa) | Df. (ppm) | Iso | DBE |
|---|---|---|---|---|---|---|---|---|---|
| 1 | 0.00 | C43 H31 N5 O13 S4 | [M-H]- | 952.0712 | 952.0728 | -1.6 | -1.68 | 0.00 | 31.0 |

*3'-(3-(3-carboxy-4-((5-(3-hydroxyphenyl)thiophene)-2-sulfonamido)phenyl)propanamido)-4-((5-(3-hydroxyphenyl)thiophene)-2-sulfonamido)-[1,1'-biphenyl]-3-carboxylic acid (**Y-9**).*

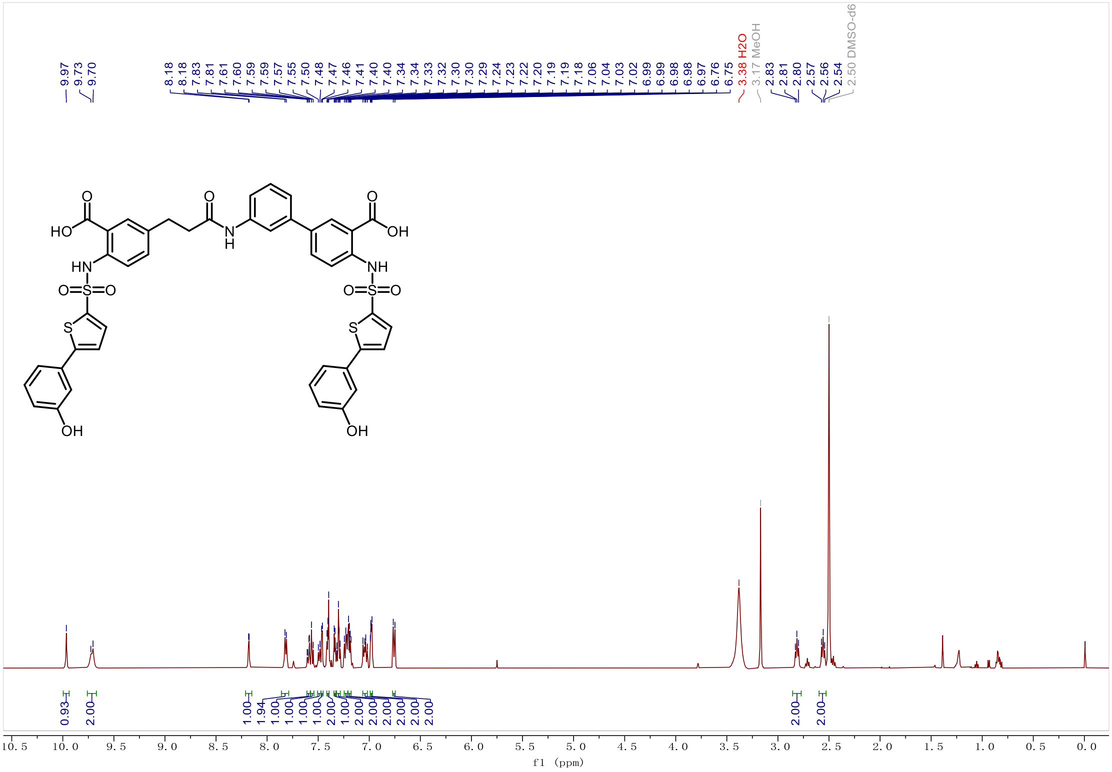

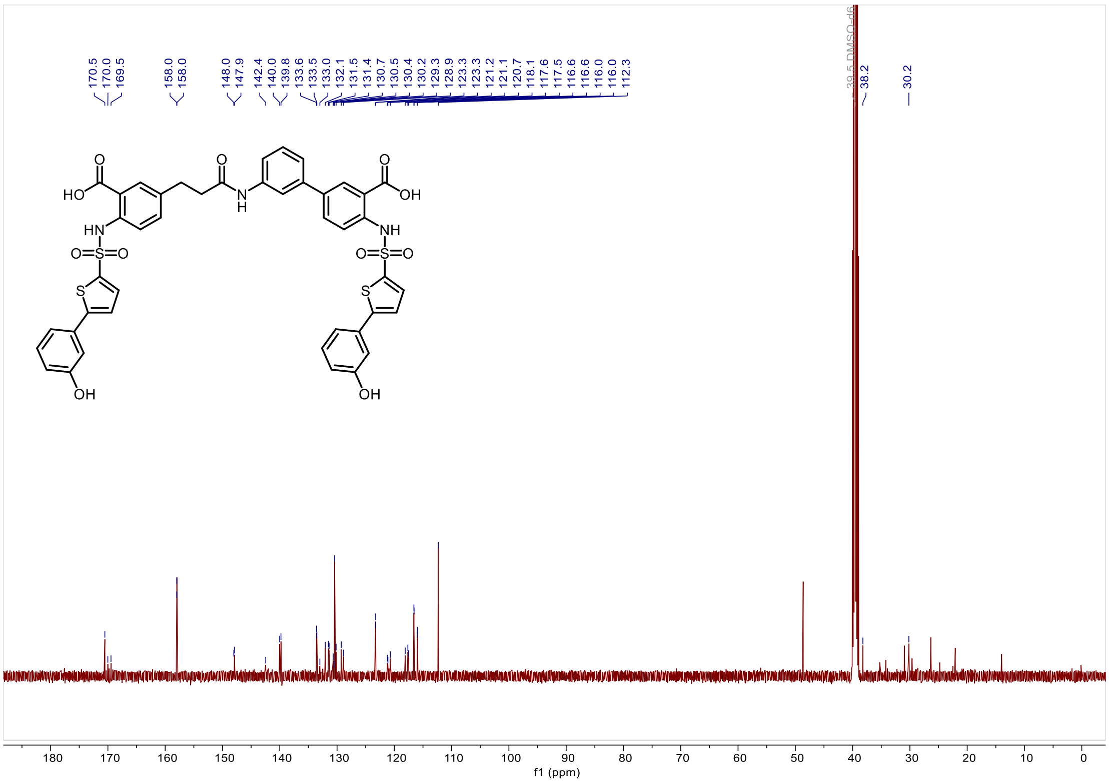

*3'-(3-(3-carboxy-4-((5-(3-hydroxyphenyl)thiophene)-2-sulfonamido)phenyl)propanamido)-4-((5-(3-hydroxyphenyl)thiophene)-2-sulfonamido)-[1,1'-biphenyl]-3-carboxylic acid (**Y-9**).*

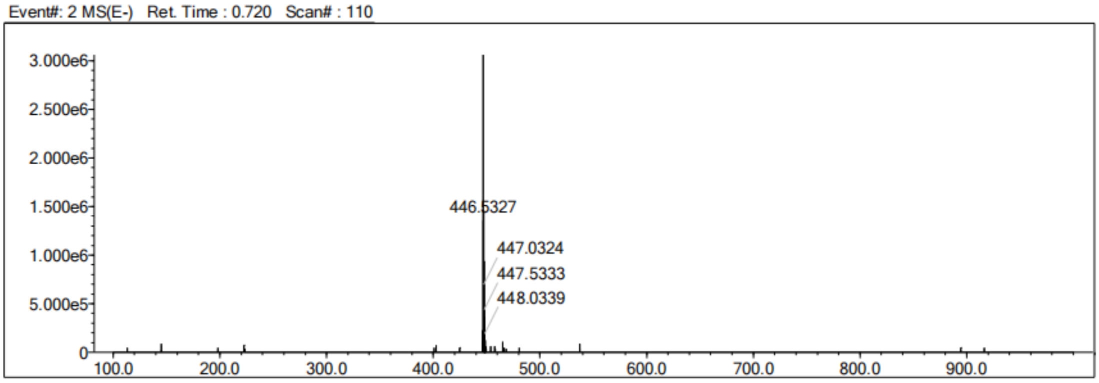


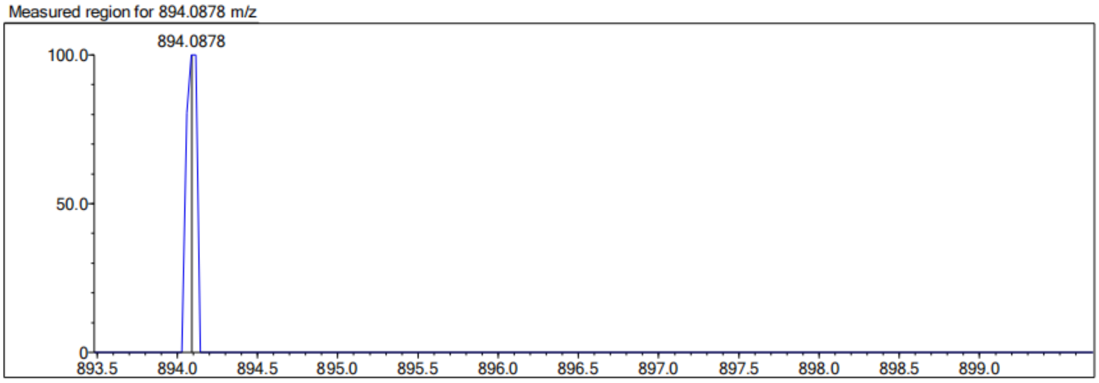


| Rank | Score | Formula (M) | Ion | Meas. m/z | Pred. m/z | Df. (mDa) | Df. (ppm) | Iso | DBE |
|---|---|---|---|---|---|---|---|---|---|
| 1 | 0.00 | C43 H33 N3 O11 S4 | [M-H]- | 894.0878 | 894.0925 | -4.7 | -5.26 | 0.00 | 29.0 |

*3'-(3-(3-carboxy-4-((5-(3-methoxyphenyl)thiophene)-2-sulfonamido)phenyl)propanamido)-4-((5-(3-methoxyphenyl)thiophene)-2-sulfonamido)-[1,1'-biphenyl]-3-carboxylic acid (**Y-10**).*

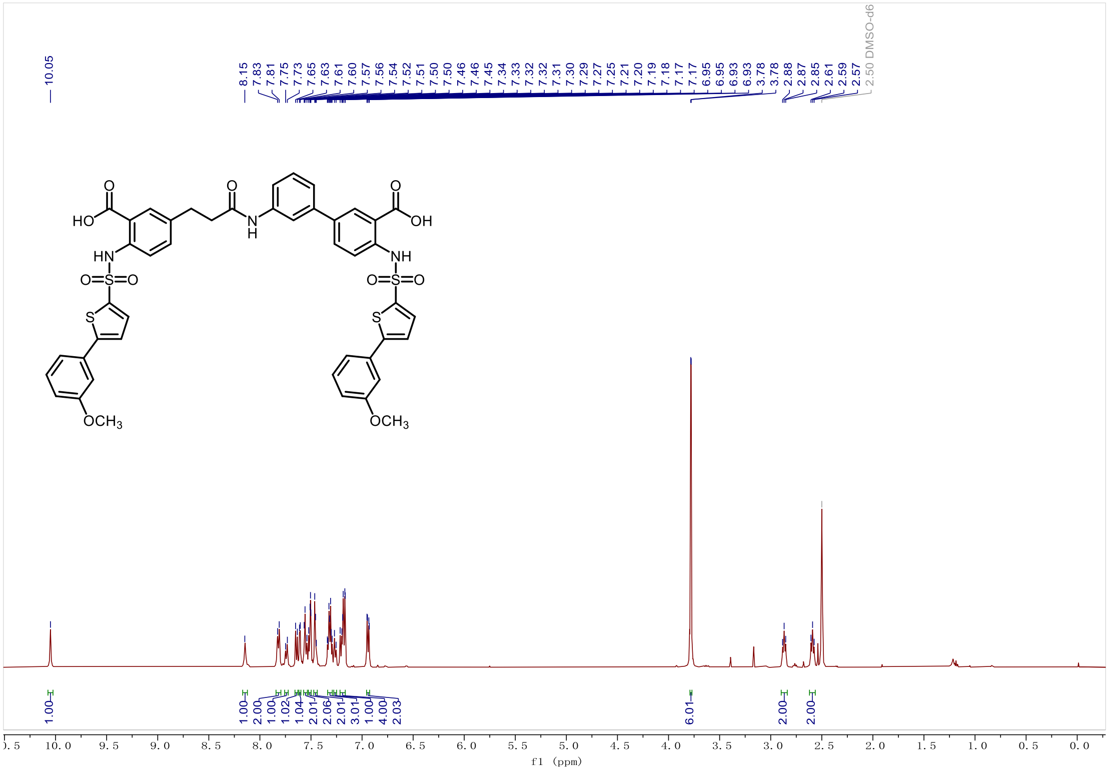

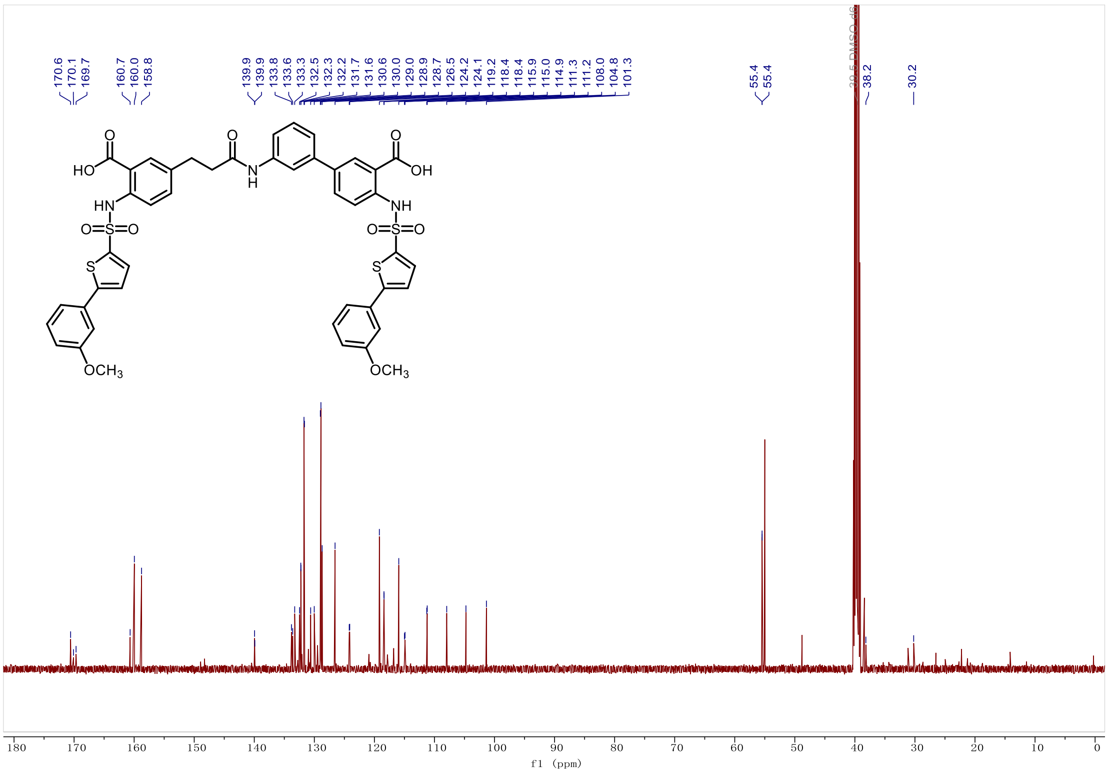

*3'-(3-(3-carboxy-4-((5-(3-methoxyphenyl)thiophene)-2-sulfonamido)phenyl)propanamido)-4-((5-(3-methoxyphenyl)thiophene)-2-sulfonamido)-[1,1'-biphenyl]-3-carboxylic acid (**Y-10**).*

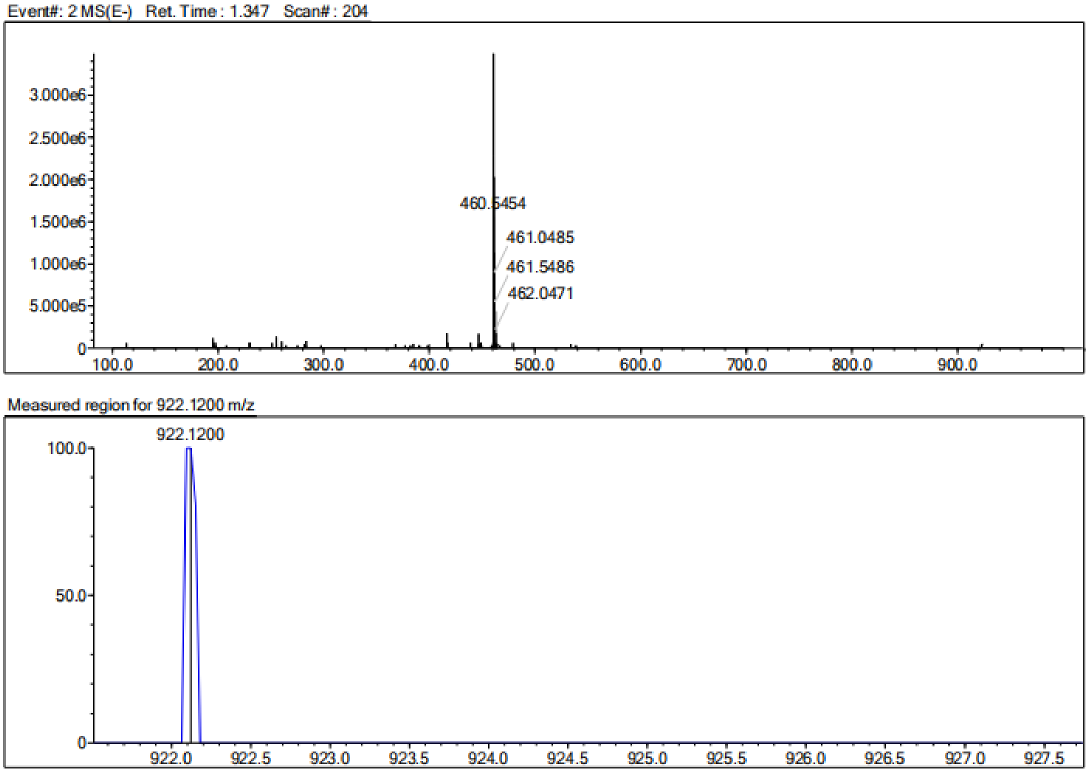


| Rank | Score | Formula (M) | Ion | Meas. m/z | Pred. m/z | Df. (mDa) | Df. (ppm) | Iso | DBE |
|---|---|---|---|---|---|---|---|---|---|
| 1 | 0.00 | C45 H37 N3 O11 S4 | [M-H]- | 922.1200 | 922.1238 | -3.8 | -4.12 | 0.00 | 29.0 |

*3'-(3-(3-carboxy-4-((5-(3,4-difluorophenyl)thiophene)-2-sulfonamido)phenyl)propanamido)-4-((5-(3,4-difluorophenyl)thiophene)-2-sulfonamido)-[1,1'-biphenyl]-3-carboxylic acid (**Y-11**).*

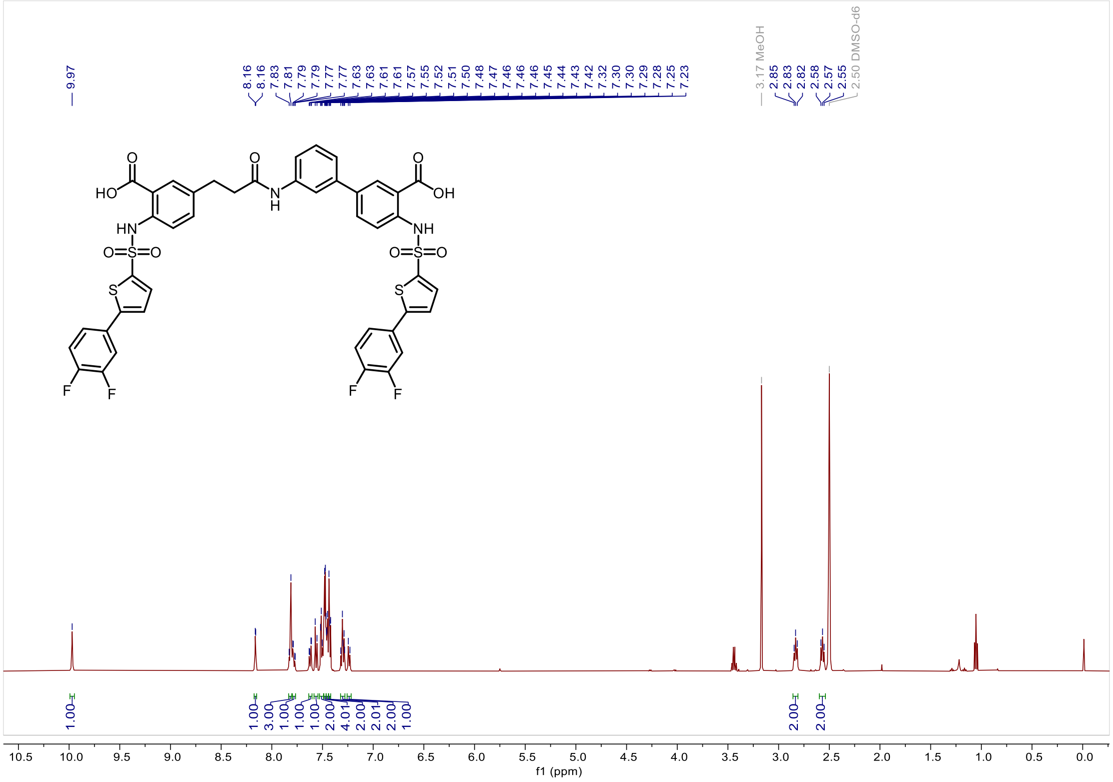

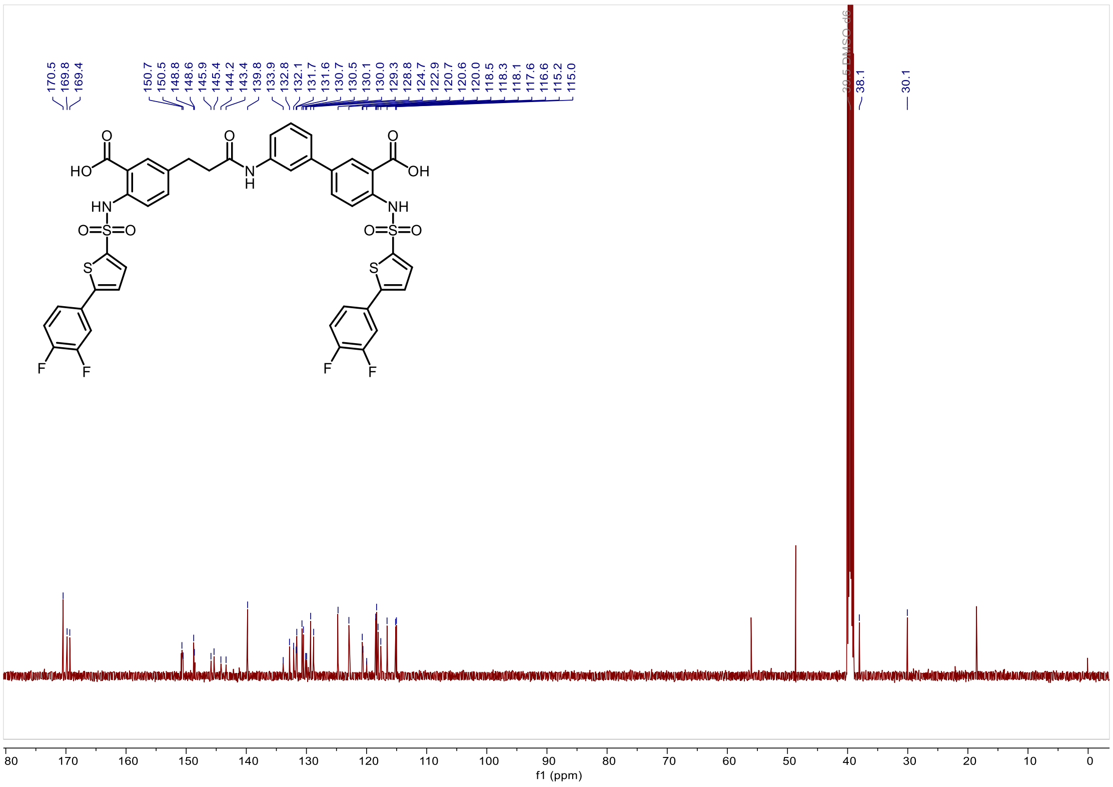

*3'-(3-(3-carboxy-4-((5-(3,4-difluorophenyl)thiophene)-2-sulfonamido)phenyl)propanamido)-4-((5-(3,4-difluorophenyl)thiophene)-2-sulfonamido)-[1,1'-biphenyl]-3-carboxylic acid (**Y-11**).*

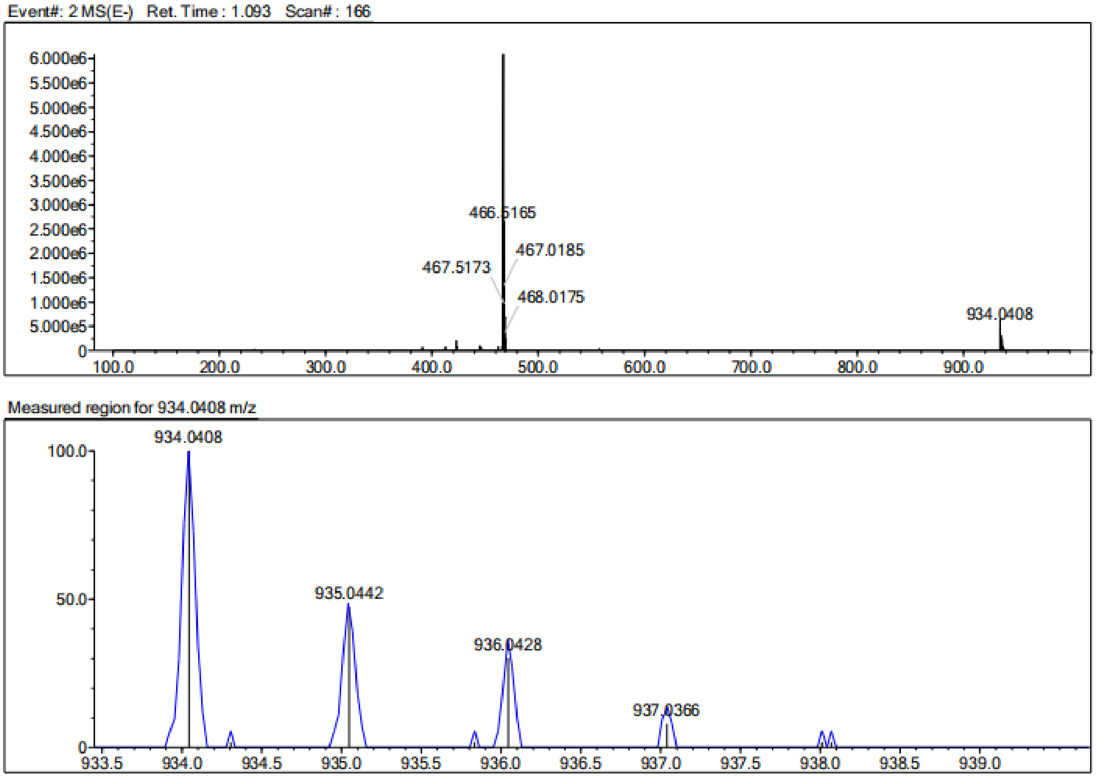


| Rank | Score | Formula (M) | Ion | Meas. m/z | Pred. m/z | Df. (mDa) | Df. (ppm) | Iso | DBE |
|---|---|---|---|---|---|---|---|---|---|
| 1 | 0.00 | C43 H29 N3 O9 F4 S4 | [M-H]- | 934.0408 | 934.0650 | -24.2 | -25.91 | 23.69 | 29.0 |

*3'-(3-(3-carboxy-4-((5-(3,4-dichlorophenyl)thiophene)-2-sulfonamido)phenyl)propanamido)-4-((5-(3,4-dichlorophenyl)thiophene)-2-sulfonamido)-[1,1'-biphenyl]-3-carboxylic acid* ***(Y-12)***.

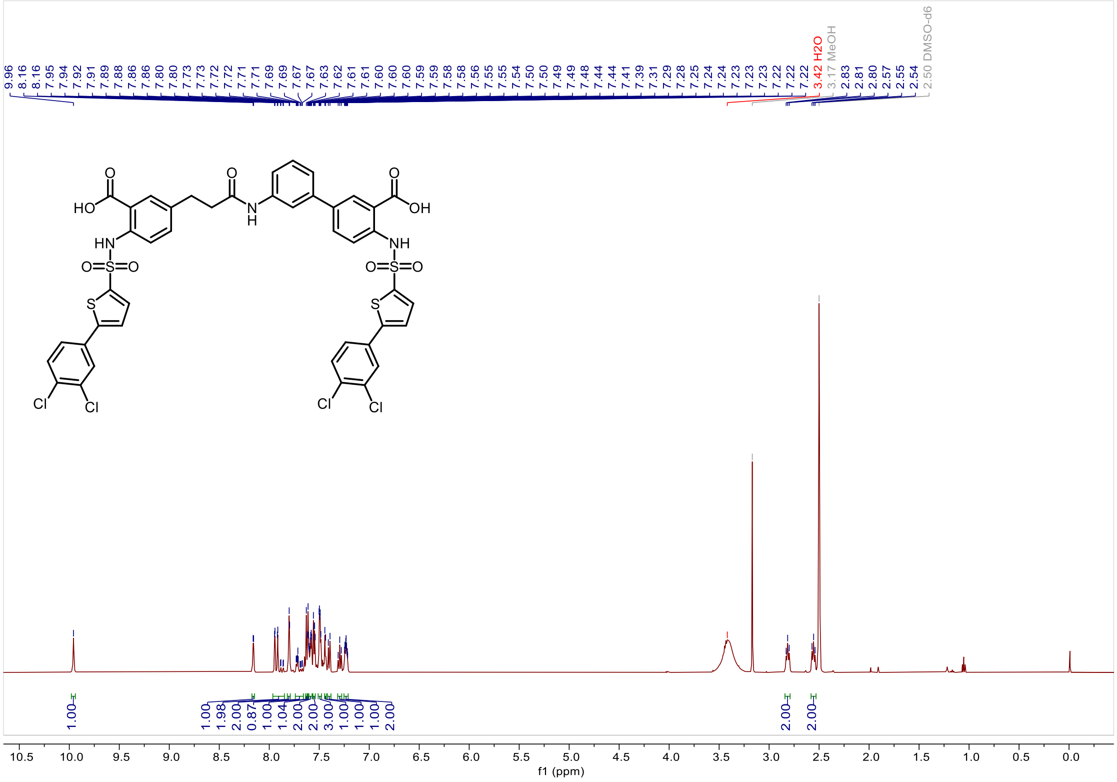

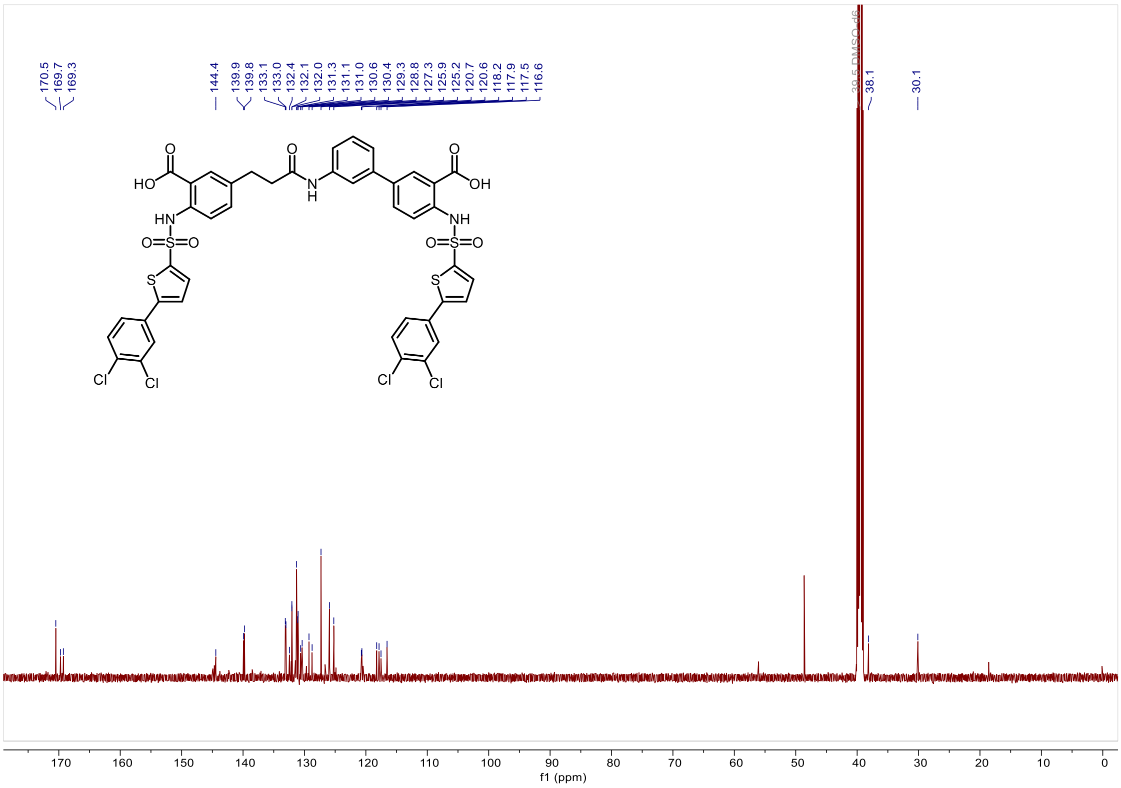

*3'-(3-(3-carboxy-4-((5-(3,4-dichlorophenyl)thiophene)-2-sulfonamido)phenyl)propanamido)-4-((5-(3,4-dichlorophenyl)thiophene)-2-sulfonamido)-[1,1'-biphenyl]-3-carboxylic acid* ***(Y-12)***.

Event#: 2 MS(E-) Ret. Time : 0.840 Scan# : 128

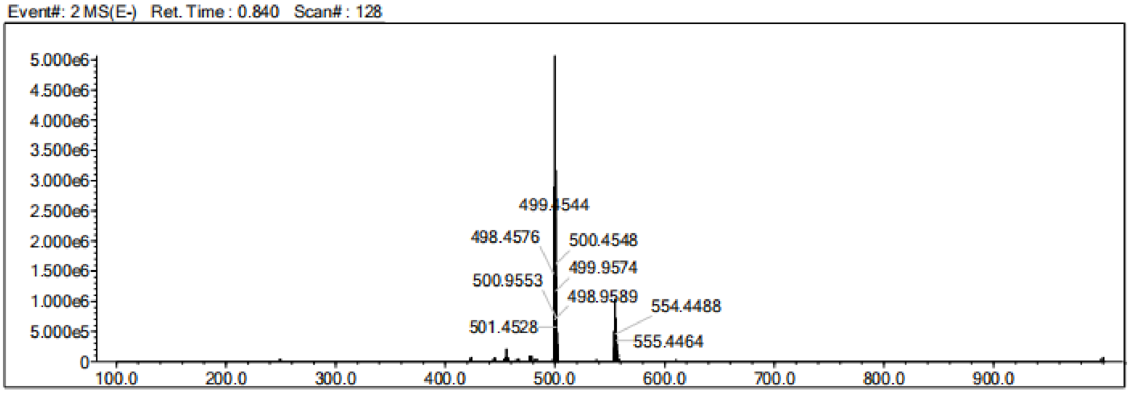


Measured region for 997.9420 m/z

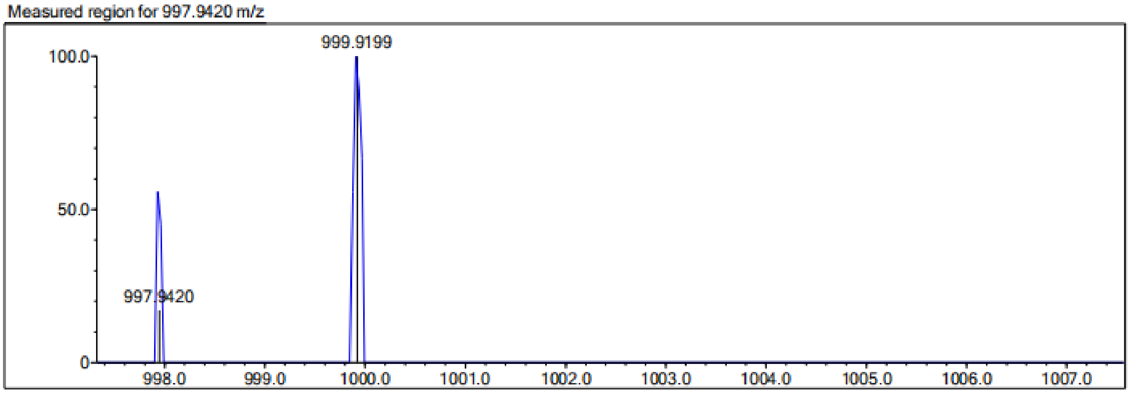


| Rank | Score | Formula (M) | Ion | Meas. m/z | Pred. m/z | Df. (mDa) | Df. (ppm) | Iso | DBE |
|---|---|---|---|---|---|---|---|---|---|
| 1 | 0.00 | C43 H29 N3 O9 S4 Cl4 | [M-H]- | 997.9420 | 997.9468 | -4.8 | -4.81 | 0.00 | 29.0 |

*3'-(3-(3-carboxy-4-((5-(3,4-dimethylphenyl)thiophene)-2-sulfonamido)phenyl)propanamido)-4-((5-(3,4-dimethylphenyl)thiophene)-2-sulfonamido)-[1,1'-biphenyl]-3-carboxylic acid (**Y-13**).*

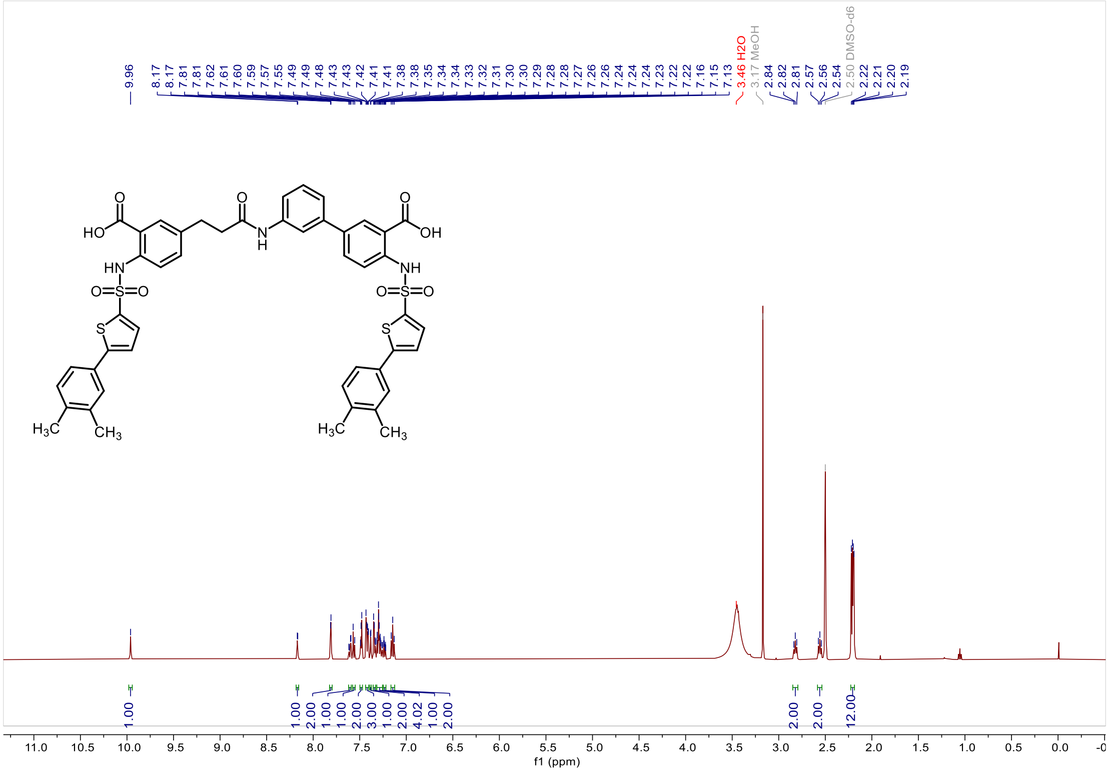


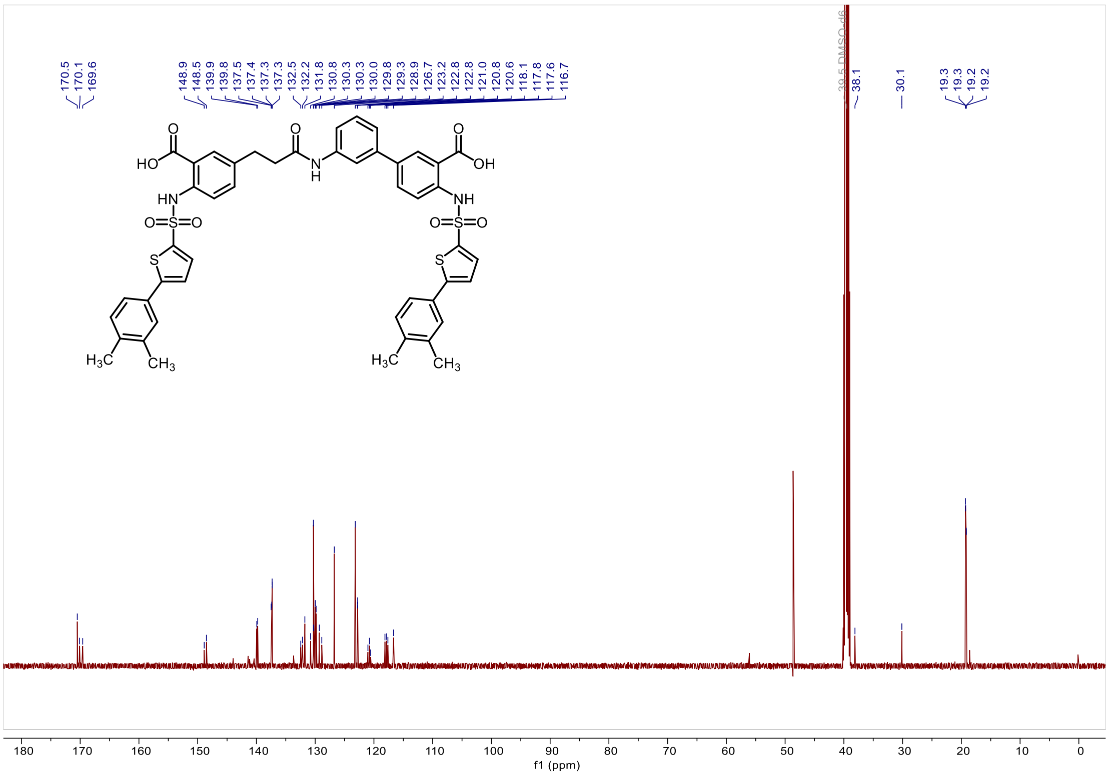

*3'-(3-(3-carboxy-4-((5-(3,4-dimethylphenyl)thiophene)-2-sulfonamido)phenyl)propanamido)-4-((5-(3,4-dimethylphenyl)thiophene)-2-sulfonamido)-[1,1'-biphenyl]-3-carboxylic acid (**Y-13**).*

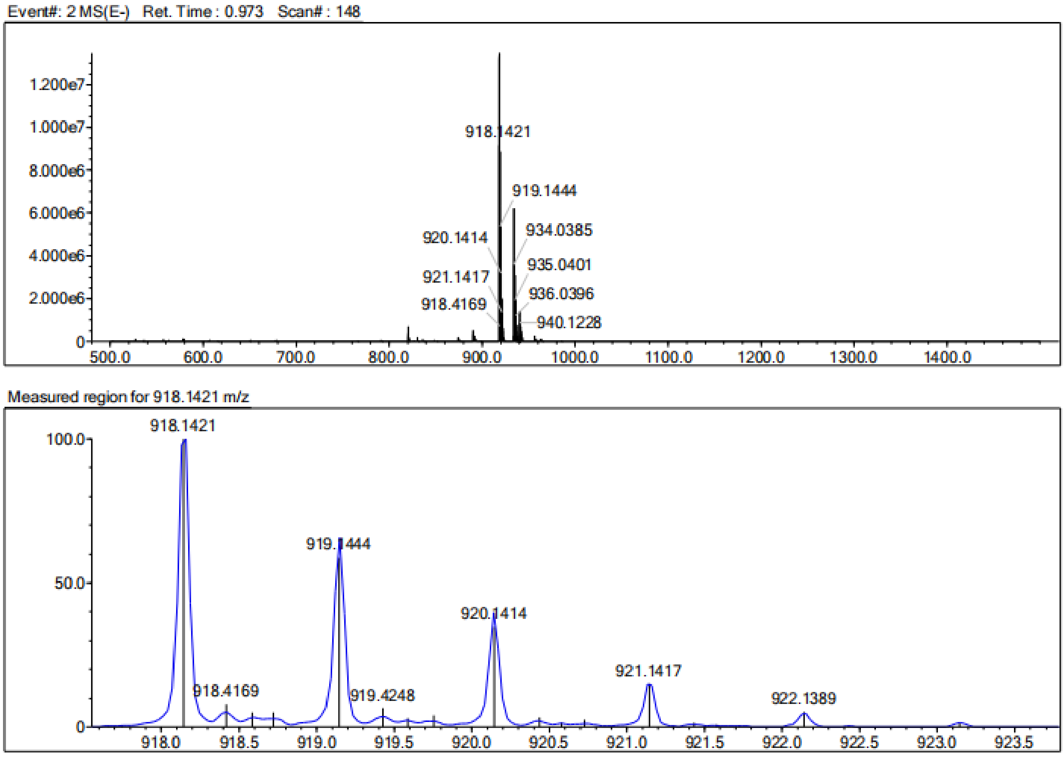


| Rank | Score | Formula (M) | Ion | Meas. m/z | Pred. m/z | Df. (mDa) | Df. (ppm) | Iso | DBE |
|---|---|---|---|---|---|---|---|---|---|
| 1 | 0.00 | C47 H41 N3 O9 S4 | [M-H]- | 918.1421 | 918.1653 | -23.2 | -25.27 | 43.87 | 29.0 |

*3'-(3-(3-carboxy-4-((5-(2,4-difluorophenyl)thiophene)-2-sulfonamido)phenyl)propanamido)-4-((5-(2,4-difluorophenyl)thiophene)-2-sulfonamido)-[1,1'-biphenyl]-3-carboxylic acid* ***(Y-14)***.

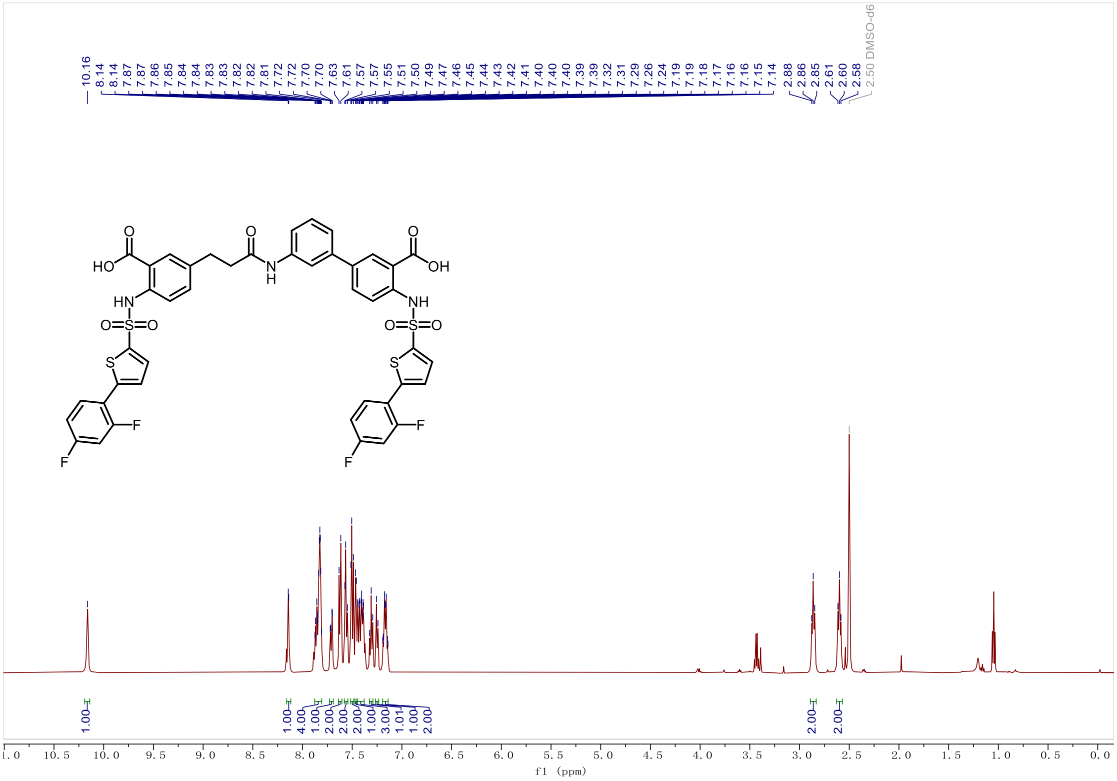


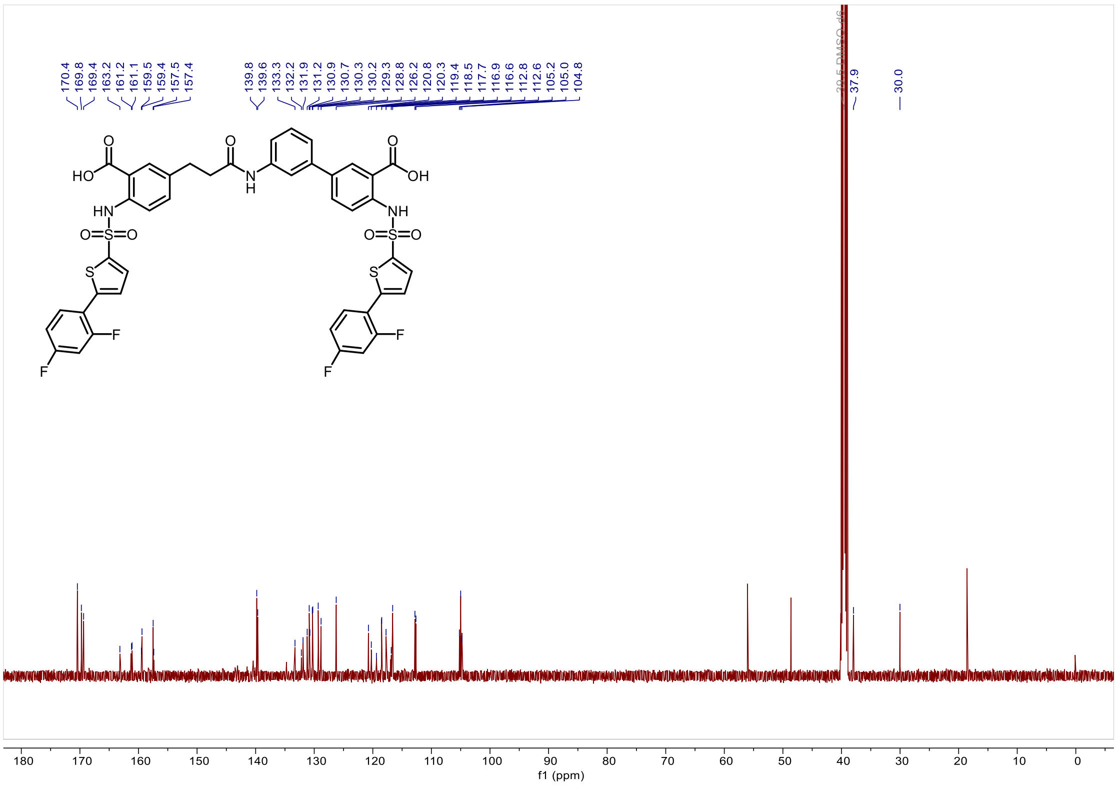

*3'-(3-(3-carboxy-4-((5-(2,4-difluorophenyl)thiophene)-2-sulfonamido)phenyl)propanamido)-4-((5-(2,4-difluorophenyl)thiophene)-2-sulfonamido)-[1,1'-biphenyl]-3-carboxylic acid* ***(Y-14)***.

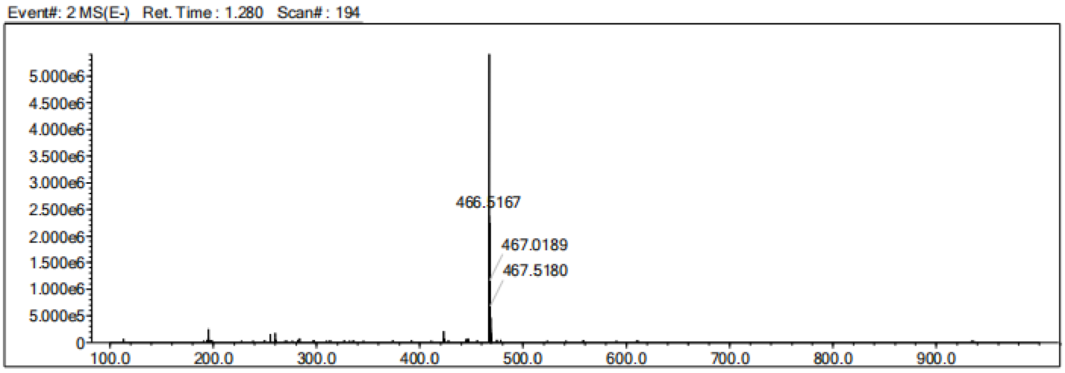


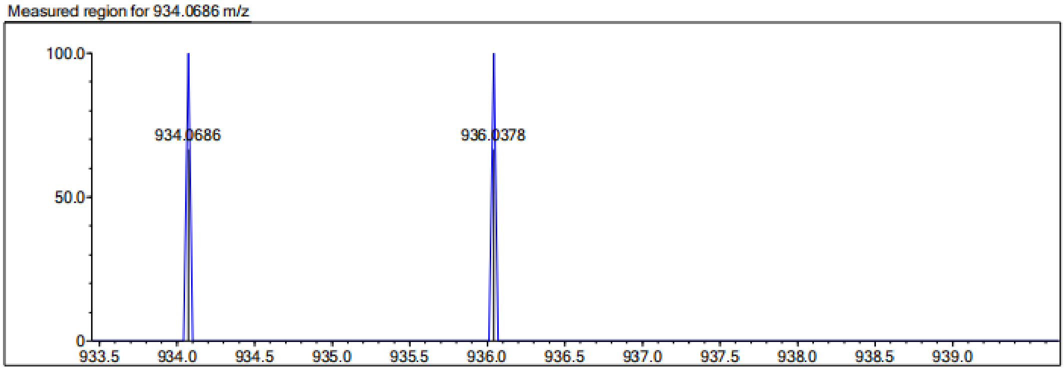


| Rank | Score | Formula (M) | Ion | Meas. m/z | Pred. m/z | Df. (mDa) | Df. (ppm) | Iso | DBE |
|---|---|---|---|---|---|---|---|---|---|
| 1 | 0.00 | C43 H29 N3 O9 F4 S4 | [M-H]- | 934.0686 | 934.0650 | 3.6 | 3.85 | 0.00 | 29.0 |

*3'-(3-(3-carboxy-4-((5-(3,5-difluorophenyl)thiophene)-2-sulfonamido)phenyl)propanamido)-4-((5-(3,5-difluorophenyl)thiophene)-2-sulfonamido)-[1,1'-biphenyl]-3-carboxylic acid (**Y-15**).*

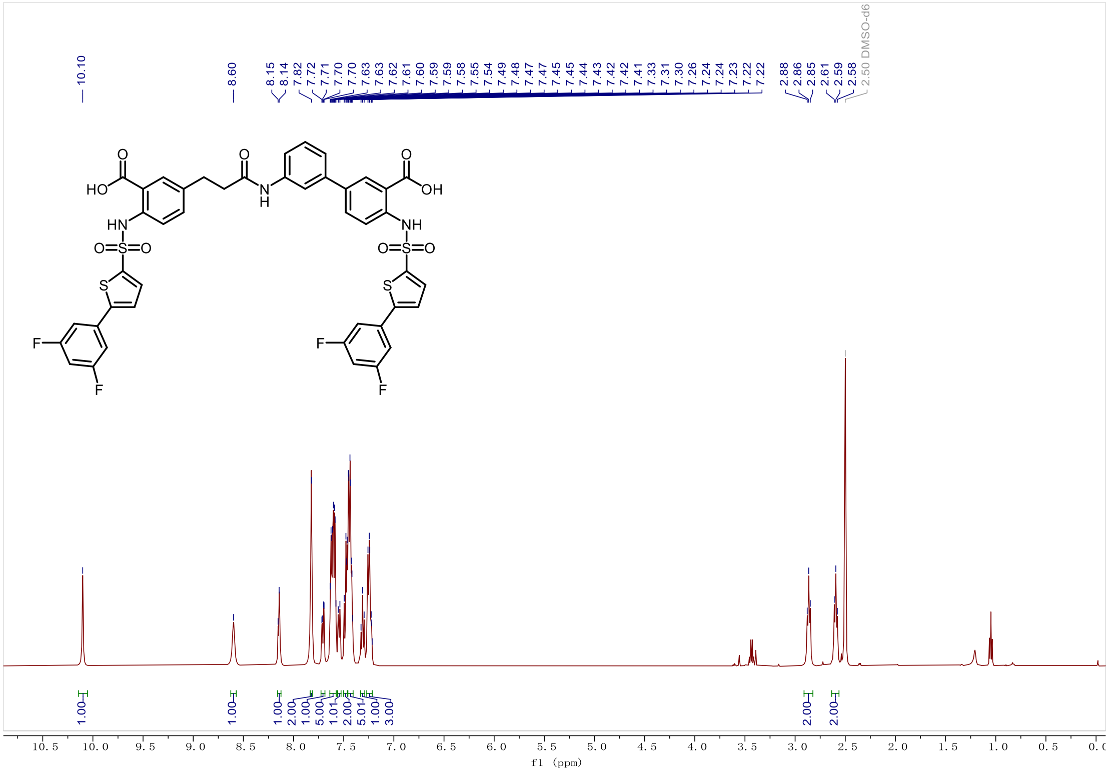

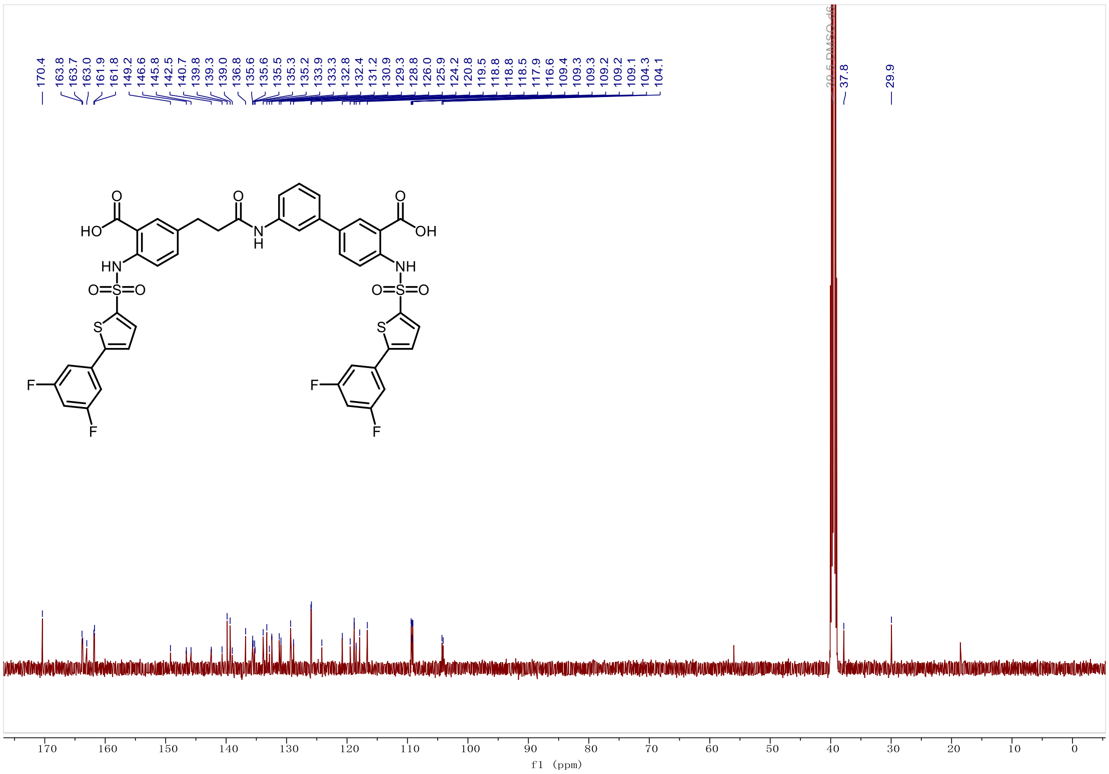

*3'-(3-(3-carboxy-4-((5-(3,5-difluorophenyl)thiophene)-2-sulfonamido)phenyl)propanamido)-4-((5-(3,5-difluorophenyl)thiophene)-2-sulfonamido)-[1,1'-biphenyl]-3-carboxylic acid* ***(Y-15)****.*

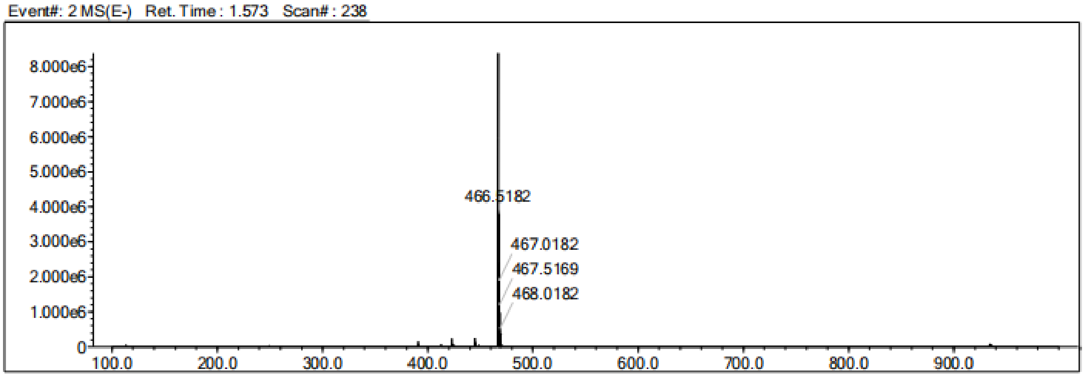


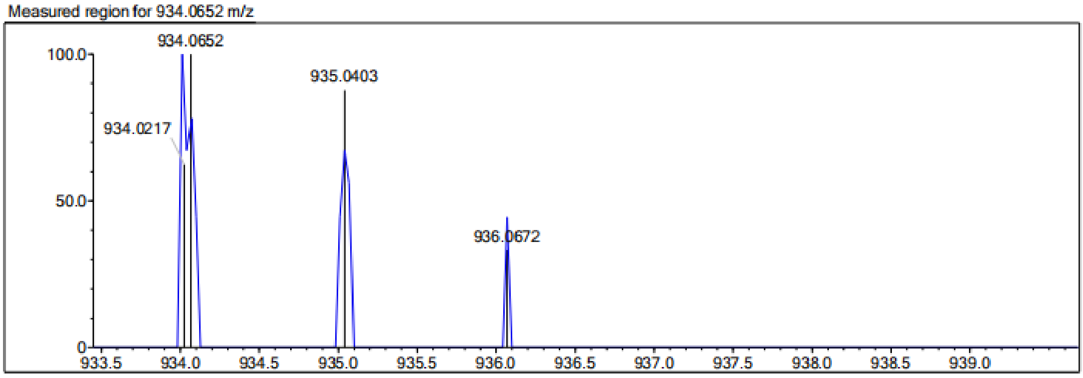


| Rank | Score | Formula (M) | Ion | Meas. m/z | Pred. m/z | Df. (mDa) | Df. (ppm) | Iso | DBE |
|---|---|---|---|---|---|---|---|---|---|
| 1 | 1.84 | C43 H29 N3 O9 F4 S4 | [M-H]- | 934.0652 | 934.0650 | 0.2 | 0.21 | 1.84 | 29.0 |

*3'-(3-(3-carboxy-4-((5-(3,4,5-trifluorophenyl)thiophene)-2-sulfonamido)phenyl)propanamido)-4-((5-(3,4,5-trifluorophenyl)thiophene)-2-sulfonamido)-[1,1'-biphenyl]-3-carboxylic acid (**Y-16**).*

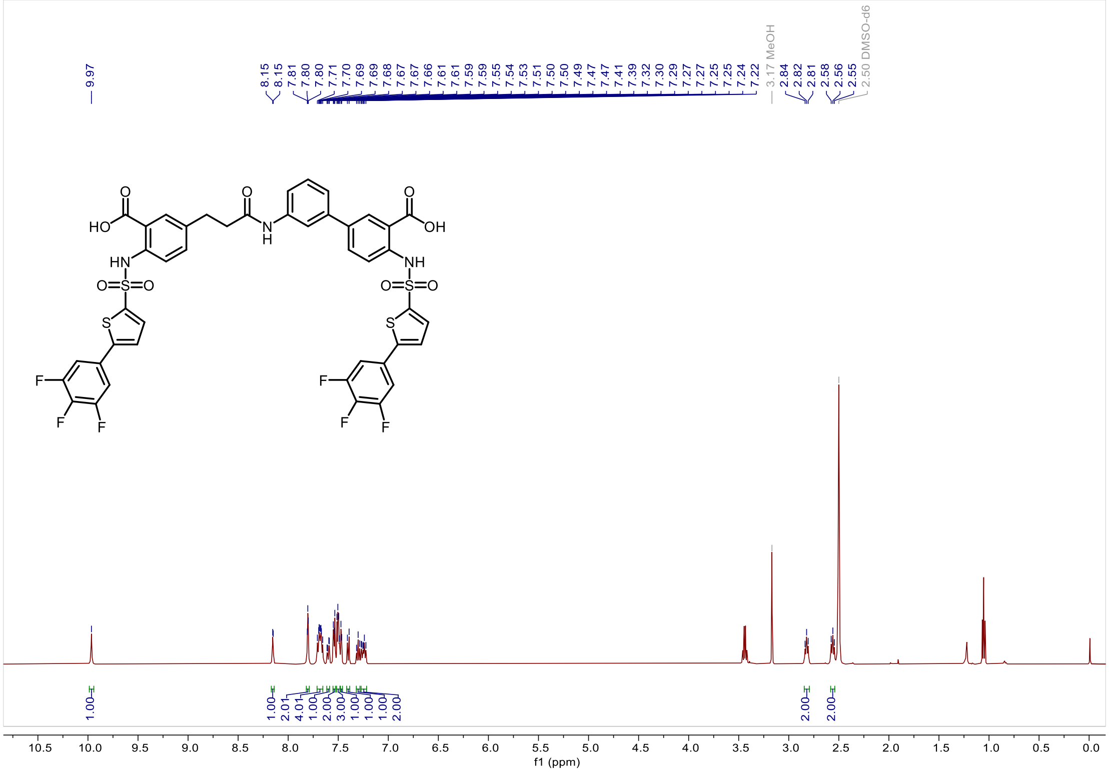


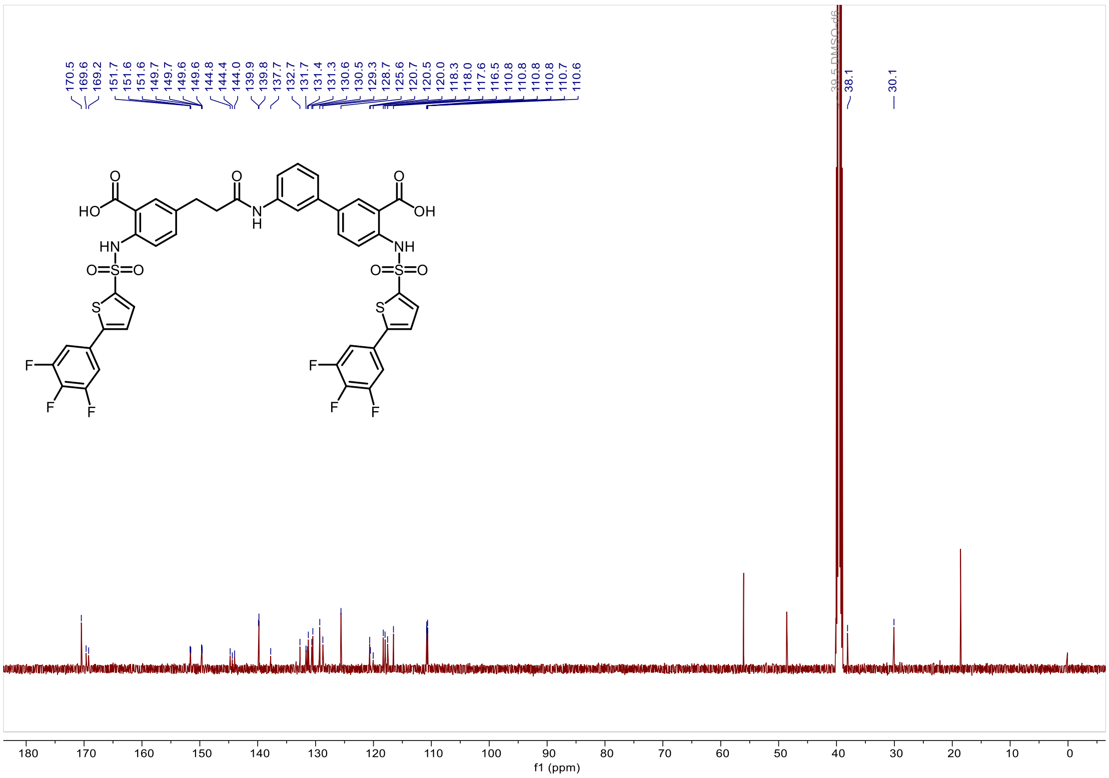

*3'-(3-(3-carboxy-4-((5-(3,4,5-trifluorophenyl)thiophene)-2-sulfonamido)phenyl)propanamido)-4-((5-(3,4,5-trifluorophenyl)thiophene)-2-sulfonamido)-[1,1'-biphenyl]-3-carboxylic acid (**Y-16**).*

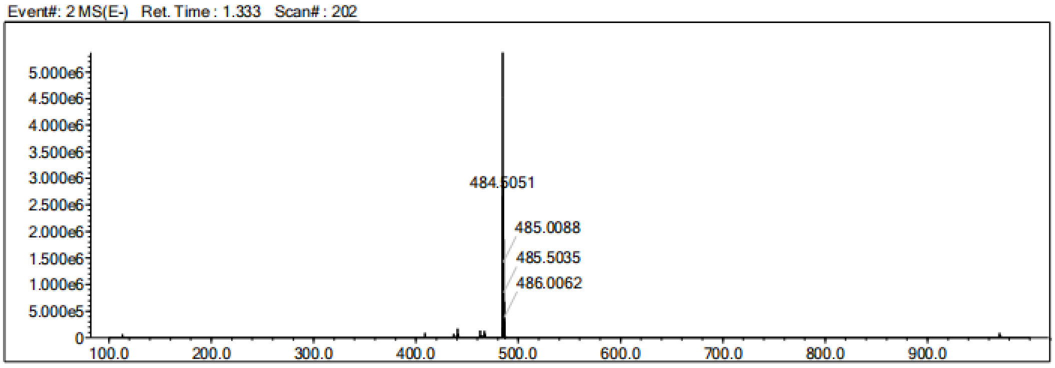


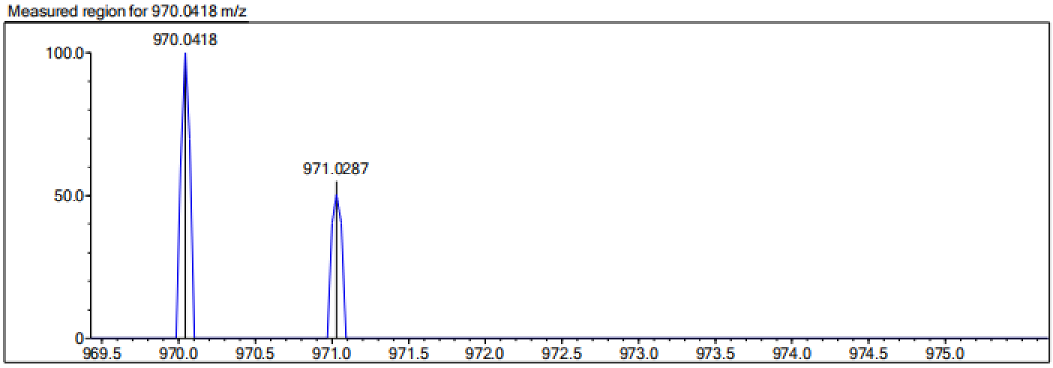


| Rank | Score | Formula (M) | Ion | Meas. m/z | Pred. m/z | Df. (mDa) | Df. (ppm) | Iso | DBE |
|---|---|---|---|---|---|---|---|---|---|
| 1 | 0.00 | C43 H27 N3 O9 F6 S4 | [M-H]- | 970.0418 | 970.0462 | -4.4 | -4.54 | 0.00 | 29.0 |

*3'-(3-(3-carboxy-4-((5-(2,3,4-trifluorophenyl)thiophene)-2-sulfonamido)phenyl)propanamido)-4-((5-(2,3,4-trifluorophenyl)thiophene)-2-sulfonamido)-[1,1'-biphenyl]-3-carboxylic acid (**Y-17**).*

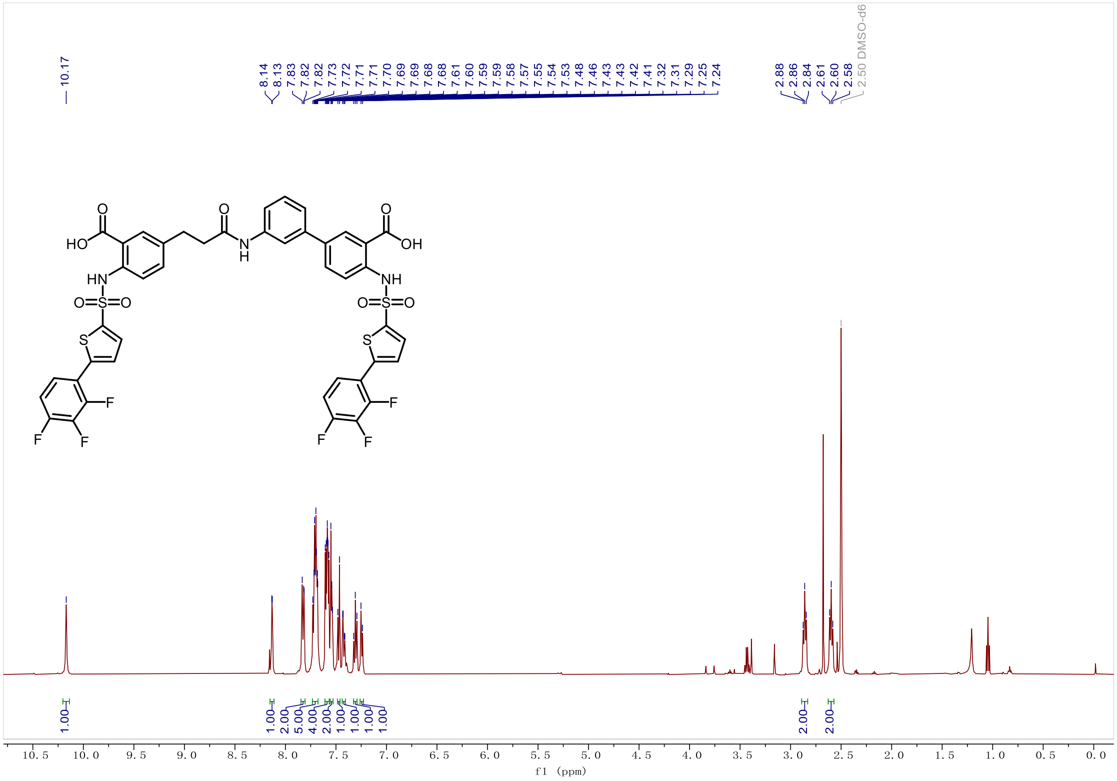

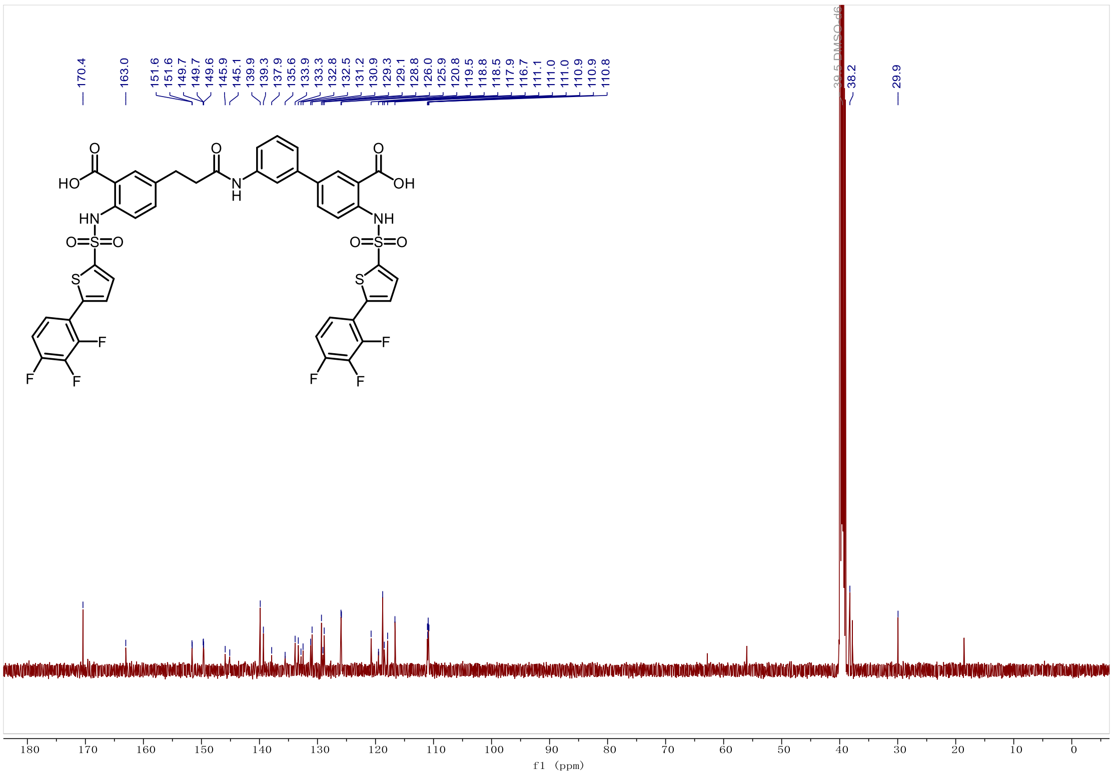

*3'-(3-(3-carboxy-4-((5-(2,3,4-trifluorophenyl)thiophene)-2-sulfonamido)phenyl)propanamido)-4-((5-(2,3,4-trifluorophenyl)thiophene)-2-sulfonamido)-[1,1'-biphenyl]-3-carboxylic acid (**Y-17**).*

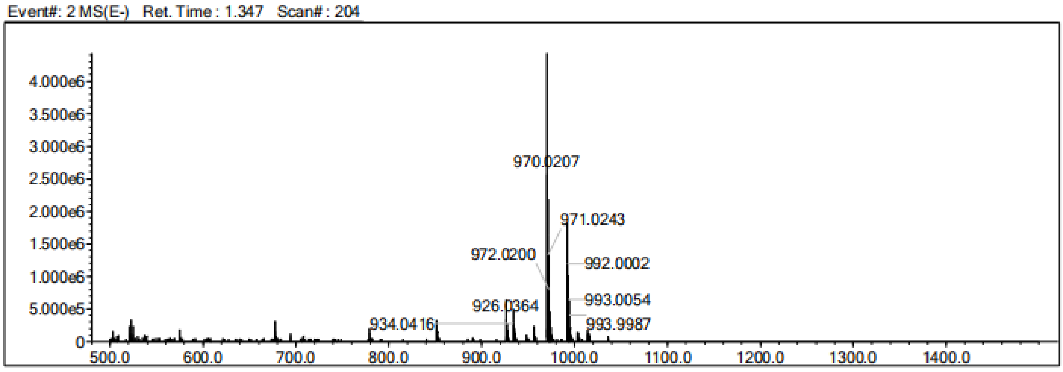


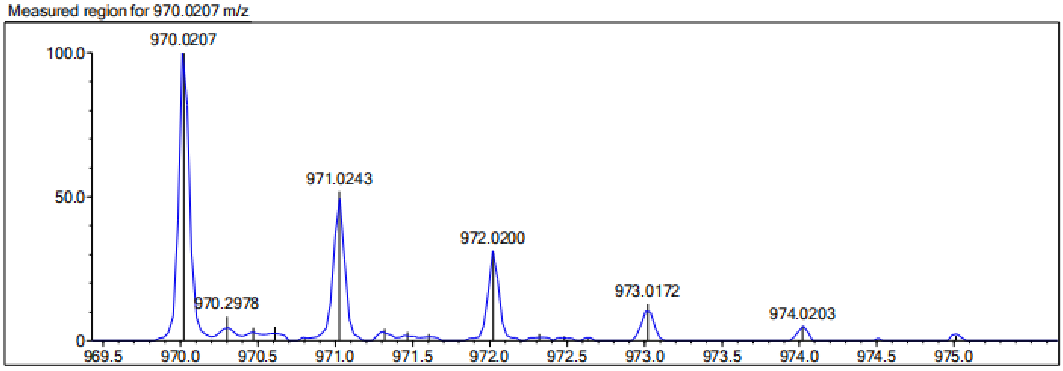


| Rank | Score | Formula (M) | Ion | Meas. m/z | Pred. m/z | Df. (mDa) | Df. (ppm) | Iso | DBE |
|---|---|---|---|---|---|---|---|---|---|
| 1 | 0.00 | C43 H27 N3 O9 F6 S4 | [M-H]- | 970.0207 | 970.0462 | -25.5 | -26.29 | 29.07 | 29.0 |

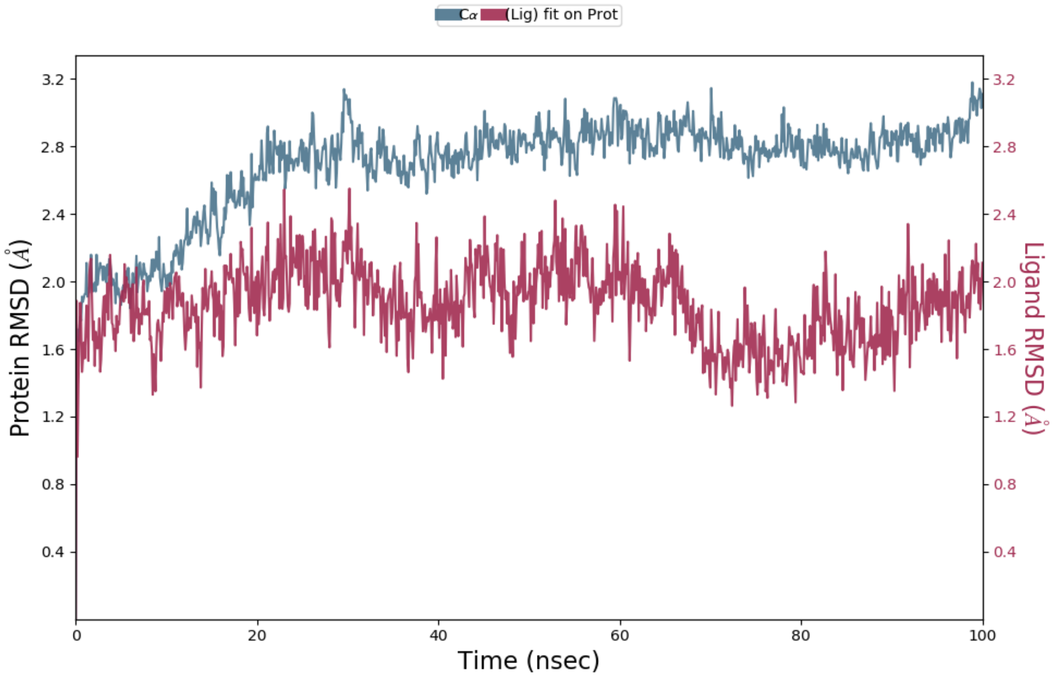


**Figure S5.** The RMSD plot of MsbA (Cα atom) and compound **Y-11** (heavy atoms) during 100-ns molecular dynamics simulation.

**Figure S6.** HPLC purity data of **Y-11** and **Cpd 4**.

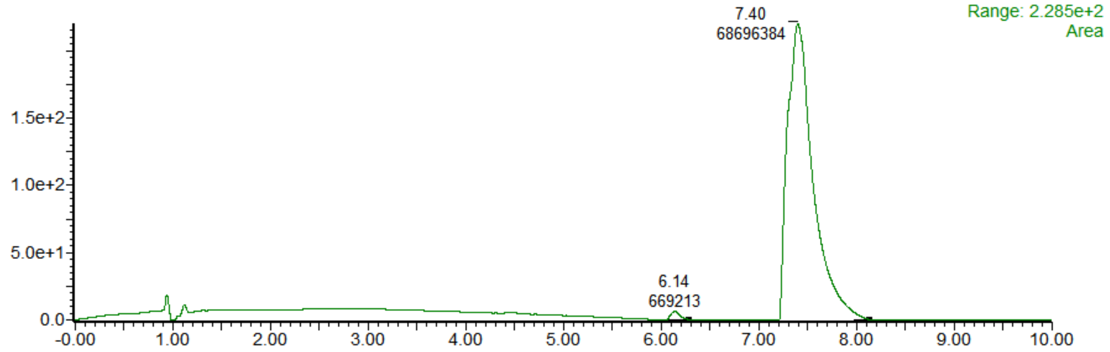


HPLC purity data of **Y-11**.

| No. | Retention Time(min) | Area(AU*s) | Area(%) |
|---|---|---|---|
| 1 | 6.14 | 669,213 | 0.96 |
| 2 | 7.40 | 68,696,384 | 99.03 |

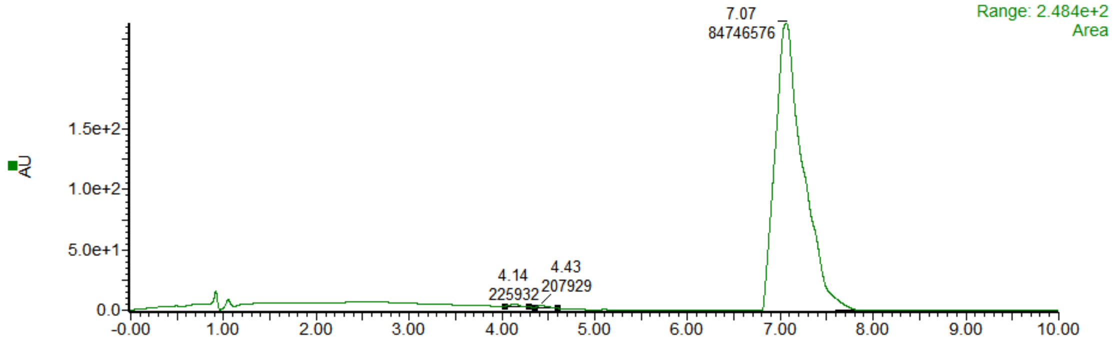


HPLC purity data of **Cpd 4**.

| Peak | Retention Time(min) | Area(AU*s) | Area(%) |
|---|---|---|---|
| 1 | 4.14 | 225,932 | 0.26 |
| 2 | 4.43 | 207,929 | 0.24 |
| 3 | 7.07 | 84,746,576 | 99.49 |
